# Sequestration-based neural networks that operate out of equilibrium

**DOI:** 10.1101/2025.05.16.654484

**Authors:** Eiji Nakamura, Frank Britto Bisso, Ignacio Gispert, Hossein Moghimianavval, Sota Okuda, Alicja Przybyszewska-Podstawka, Samuel Chisholm, Varunaa Sri Hemanth Kumar, Valerie Perez Medina, Ming-Ru Wu, Christian Cuba Samaniego

**Author notes:** Synthetic Biology Summer School at Cold Spring Harbor 2024 (Eiji Nakamura, Frank Britto Bisso, Ignacio Gispert, Sota Okuda, Alicja Przybyszewska-Podstawka, Samuel Chisholm, Varunaa Sri Hemanth Kumar, Valerie Perez Medina, Christian Cuba Samaniego). Equal contribution (Eiji Nakamura, Frank Britto Bisso, Ignacio Gispert).

## Abstract

Classification of high-dimensional information is a ubiquitous computing paradigm across diverse biological systems, including organs such as the brain, down to signaling between individual cells. Inspired by the success of artificial neural networks in machine learning, the idea of engineering genetic circuits that operate as neural networks emerges as a strategy to expand the classification capabilities of living systems. In this work, we design these biomolecular neural networks (BNNs) based on the molecular sequestration reaction, and experimentally characterize their behavior as linear classifiers for increasing levels of complexity. Initially, we demonstrate that a static, DNA-based system can effectively prototype a linear classifier, though we also identify its limitations to easily tune the slope of the decision boundary it generates. We then propose and experimentally validate a BNN at the protein level using a cell-free transcription–translation (TXTL) system, which overcomes the DNA-based system’s limitation and behaves as a linear classifier even before it reaches its steady state (or, equivalently, out-of-equilibrium). Ultimately, we test a CRISPR-based design and its out-of-equilibrium behavior in a biological context by successfully constructing a linear classifier within mammalian cells. Overall, by leveraging mathematical modeling and experimental automation, we establish molecular sequestration as a universal scheme for implementing neural networks within living systems, paving the way for transformative advances in synthetic biology and programmable biocomputing systems.

## 1 Introduction

Classification involves assigning labels to patterns based on characteristics shared among instances with that same label. Cells are no exception: they sense their environment, activate signaling pathways that process this input, and ultimately drive a phenotypic response. For instance, neural progenitor cells integrate morphogen signals from the Sonic Hedgehog and BMP pathways to shape the neural tube [1], while immune cells decode combinations of cytokines to determine whether to proliferate, initiate cell death, or activate or suppress their function [2]. Since the earliest reports of genetic circuits [3], synthetic biology has aimed to rewire the logic behind these cellular classification tasks in several ways: developing tools to detect new types of patterns (e.g., CAR-T cells that use chimeric surface receptors to recognize new — cancerous — patterns [4]), associating known patterns with new outputs (e.g., triggering cell death upon toxin detection [5]), or both (e.g., SynNotch receptors that generate custom input/output responses [6]). This kind of biocomputation was initially approached using digital logic, where gene expression was conceptualized as either “on” or “off” [7, 8]. However, biological patterns are far more complex: they are analogue [9, 10, 11], high-dimensional (e.g., molecular profiles that define a cell’s phenotype can involve dozens [12] to hundreds [13] of biomarkers), and even of diverse nature — not just chemical, but also mechanical, thermal, or optical [14]. Capturing this complexity using digital (Boolean) circuits is then limited by design.

Early work on the computational capabilities of gene regulatory networks [15, 16] inspired the design of synthetic circuits that mimic artificial neural networks. The core unit of these systems is the perceptron, a single information-processing element that computes a weighted sum of its inputs and feeds it through a nonlinear activation function to generate an output. These perceptrons have been implemented in cell-free systems for pattern recognition tasks of varying complexity, ranging from 4-bit inputs [17] to recent designs processing 100-bit inputs through hundreds of parallel DNA hybridization reactions [18, 19, 20, 21, 22]. Similar architectures have also been implemented in living cells, computing at transcriptional [23, 24], post-translational [25], and metabolic [26] levels. One notable example used the network’s output to trigger programmed cell death [25], highlighting its potential for diagnostic and therapeutic applications [27, 28]. However, all of these implementations have been studied at equilibrium, after the underlying reactions reached steady-state, where the production rate of each chemical species is equal to its degradation.

In previous work, we theoretically showed that molecular sequestration, where two chemical species reversibly bind into an inactive complex, approximates a subtraction operation, and thus can be exploited to realize a biomolecular neural network [29]. Here, we took a “bottom-up” approach to experimentally characterize this chemical reaction at different levels of complexity, with the ultimate goal of building an in-vivo sequestration-based biomolecular perceptron. We begin by analyzing the simplest sequestration reaction: DNA hybridization, which enables the rapid prototyping of a linear classifier once the reaction reaches its steady-state. We then mathematically study the molecular sequestration reaction’s behavior as a linear classifier before the reaction reaches steady-state (i.e., out-of-equilibrium), and experimentally validate this behavior in a cell-free transcription-translation (TXTL) system using proteins. Finally, we extended the mathematical analysis into a more realistic context by considering the degradation of the perceptron’s elements, and conducted a proof-of-concept experiment in mammalian cells, showing that linear classification is feasible within a biological context. In this way, and by combining mathematical models and high-throughput experiments, we systematically characterized the temporal behavior of a sequestration-based perceptron in three different biochemical implementations across a hierarchy of cellular contexts.

## 2 Results

### 2.1 Rapid prototyping of a sequestration-based perceptron using DNA hybridization

We begin by introducing a simple molecular perceptron architecture that uses a molecular sequestration reaction, followed by its experimental implementation using DNA hybridization in which two complementary, single-stranded DNA molecules bind to form a duplex. Figure 1A presents a diagram of a molecular sequestration reaction. A molecular sequestration reaction is a bimolecular process in which one species binds its target to form an inactive complex, thereby reducing the concentration of the free target. First, we mathematically validate that this chemical reaction system functions as a molecular perceptron. The fundamental chemical reaction in the system is as follows:

**Figure 1.**
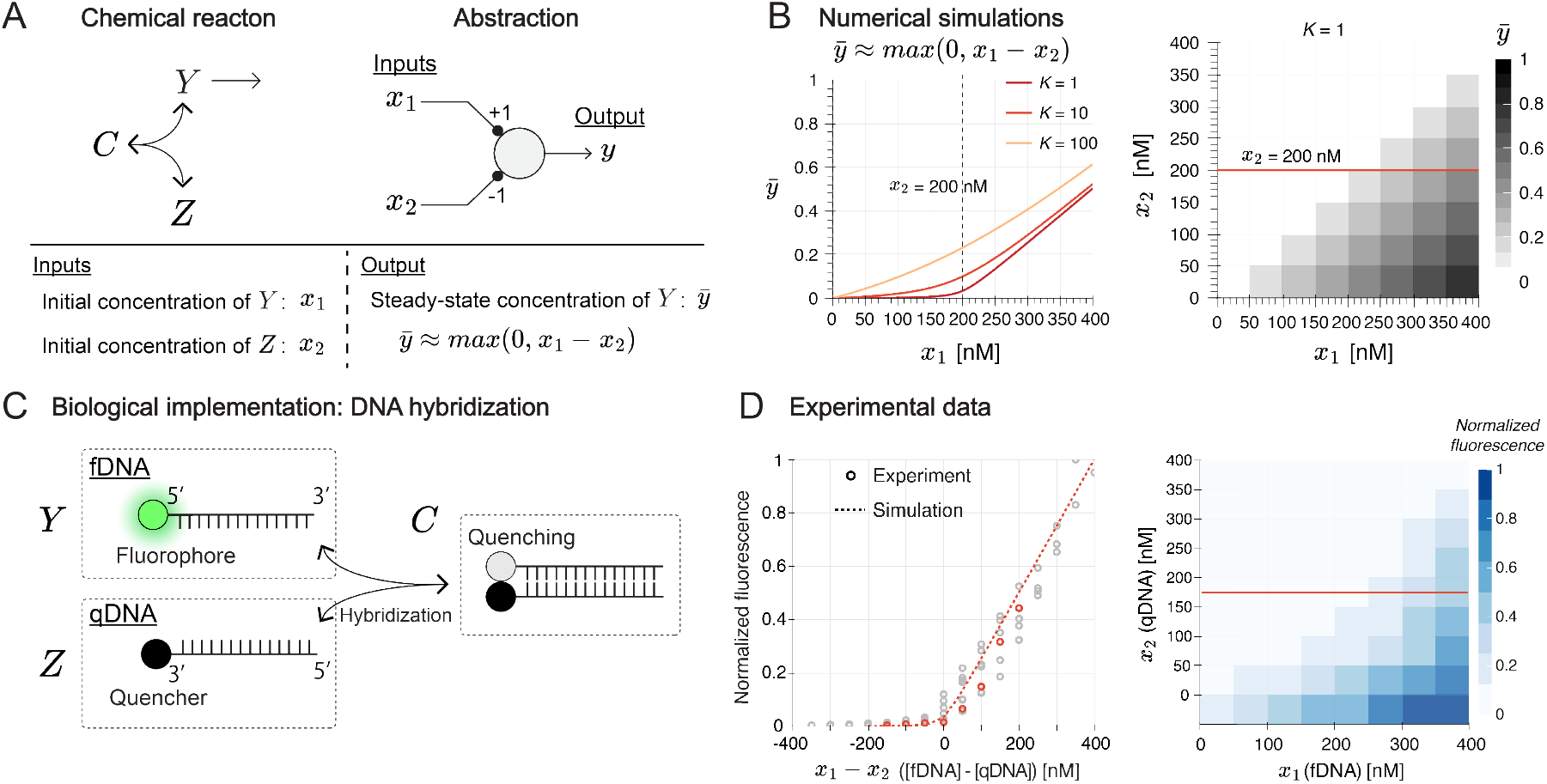
A biomolecular perceptron based on DNA hybridization. (A) Schematic diagrams of a sequestration reaction and the resulting perceptron computation. In the system, species Y serves as an output, and species Z inactivates Y through the formation of an inactive complex C. We take the initial concentrations of Y and Z as inputs x_1_ and x_2_, and the steady-state concentration of 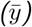 as the output. (B) Numerical simulation results of sequestration-based perceptron. The left panel shows 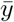 as a function of x_1_ for x_2_ = 0.5 (constant), with varying dissociation constant K (1, 10, 100). The right panel shows a contour map of 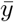 on the x_1_-x_2_ plane with K = 1. The red horizontal line indicates x_2_ value, along which the relationship between x_1_ and 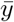 gives the curve for K = 1 shown in the left panel. (C) Biochemical implementation based on DNA hybridization. The system has two complementary DNA species: fDNA with 5’-end fluorophore (TYE563), and qDNA with 3’-end quencher (Iowa Black RQ). We represents fDNA as Y, qDNA as Z, and fDNA-qDNA complex as C. We take Y as the output, which can be monitored by the fluorescence from single-stranded fDNA. The initial concentrations of fDNA and qDNA are defined as x_1_ and x_2_ respectively. (D) The results of fluorescence measurement by a plate reader after 1-hour incubation at 37^°^C. Fluorescence intensity was normalized from 0 to 1 based on the minimum and maximum fluorescence measured in a single plate. The left panel shows scatter plots of the fluorescence data as a function of x_1_ − x_2_ (i.e., [fDNA] [qDNA]) with x_2_ = 200 nM (constant). The dashed lines show numerical simulation results of the normalized fluorescence as a function of x_1_ − x_2_ with varying x_2_ The red dots are experimental results with x_2_ = 200 nM, and the red dashed line corresponds the simulation results with x_2_ = 200 nM. The right panel shows a contour map of normalized fluorescent intensity on the x_1_-x_2_ plane. The red horizontal line indicates x_2_ value, along which the relationship between x_1_ − x_2_ and the fluorescence intensity corresponds to the red dots in the left panel.

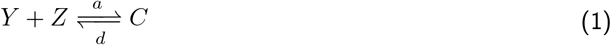

Species *Y* and *Z* reversibly bind to form an inactive complex *C*. By considering mass conservation (*y* + *c* = *x*_1_ and *z* + *c* = *x*_2_), the ordinary differential equation for *y* with respect to time (denoted as 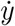) is obtained as follows:

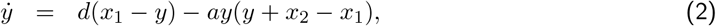

where *y, z*, and *c* are the concentrations of species *Y, Z, C*, and 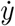 denotes the derivative of *y* with respect to time. Parameters *a, d* are the association rate constant and the dissociation rate constant of the binding reaction between *Y* and *Z*, respectively. *x*_1_ and *x*_2_ are initial concentrations of *Y* and *Z*. At steady-state, by solving 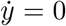 with a small dissociation constant (*K* = *d/a*), the steady-state value of *y*, denoted by 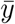, is derived as follows:

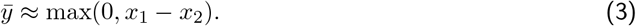

We interpret this equation as a perceptron computation [29], assigning a positive weight to *x*_1_ and a negative weight to *x*_2_. Figure 1A (right) shows an abstract representation of the perceptron. Note that the magnitudes of weights for *x*_1_ and *x*_2_ are one because we do not have any transformation or production reaction in the system. A more detailed mathematical derivation of the approximate steady-state solutions is provided in Section 1 of the Supporting Information.

Figure 1B (left) shows numerical simulations of the steady-state value of *y* (named 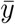) versus *x*_1_ with *x*_2_ = 200 nM and three different dissociation constants (*K* = 1, 10 and 100). As *K* decreases, the 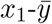 curve approaches a Rectified Linear Unit (ReLU) function-like behavior, where the output remains zero below a threshold input and increases linearly beyond it [30]. This trend is supported by the fact that the molecular sequestration mechanism executes a subtraction operation at a high sequestration reaction rate [31]. i.e., at each set of values (*x*_1_, *x*_2_), the perceptron computes 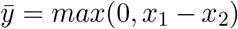. Figure 1B (right) shows a heatmap of the 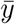 value on the *x*_1_-*x*_2_ plane with *K* = 1. There is a clear boundary on *x*_1_ = *x*_2_, along which *X*_1_ and *X*_2_ cancel each other out through the sequestration reaction. We call this a decision boundary. 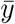 values along the red line in Figure 1B (right) correspond to the curve with *K* = 1 in Figure 1B (left). These results show that the biomolecular perceptron presented here executes a linear classification operation and a faster sequestration rate (i.e., smaller dissociation constant *K*) makes the boundary sharper.

For an experimental proof-of-concept, we propose the biochemical realization of a molecular perceptron utilizing DNA hybridization. Figure 1C shows a simplified schematic illustration of a proposed system. A more accurate and detailed representation of this system is provided in Supplementary Figure 1. Here, we have two DNA strands: a DNA strand with a fluorophore on its 5’-end, which we call fDNA, and a DNA strand with a quencher on its 3’-end, which we call qDNA. fDNA and qDNA have complementary sequence domains, so they can bind to each other if they are present in the same solution. When fDNA and qDNA bind, the fluorophore and the quencher are in close proximity, which results in fluorescence quenching. Therefore, we can monitor the binding state of the fDNA and qDNA by measuring the fluorescence. This DNA hybridization system can be associated with the chemical reaction model shown in Figure 1A, by assigning the single-stranded fDNA to *Y*, the single-stranded qDNA to *Z*, and the double-stranded fDNA-qDNA to *C* respectively. The DNA hybridization-based molecular perceptron can be described by the same mathematical model as that of the abstract chemical perceptron presented in Figure 1A with appropriate scaling of variables and parameters. To experimentally characterize the proposed system, we first set initial concentrations of fDNA and qDNA (*x*_1_ and *x*_2_ respectively), then mix them in a single solution to trigger the hybridization reaction. After incubating at 37^*°*^C for one hour, fluorescence was measured.

The experimental results presented in Figure 1D successfully validates the operation of the DNA hybridization-based perceptron. The left panel of Figure 1D shows experimental data and numerical simulation results of normalized fluorescence signals versus *x*_1_ *− x*_2_ with *x*_2_ = 200 nM. The right panel of Figure 1D shows a heatmap of normalized fluorescence signals on *x*_1_-*x*_2_ plane. Figure 1D (left) corresponds to the data in Figure 1D (right) along with the red horizontal line at *x*_2_ = 200 nM. Overall, the experimental data and numerical simulation results are in a good agreement, which demonstrate that the DNA hybridization-based system enables rapid prototyping of a molecular perceptron.

Next, we evaluated whether additional patterns of decision boundary could be constructed by changing how DNA species were assigned to the inputs. Figure 2 presents numerical simulations and experimental demonstrations of two different boundaries. The left schematic of each panel represents a biomolecular perceptron, in which we have two inputs, *X*_1_ and *X*_2_, and a constant bias *X*_0_. Figure 2 Panel A and C correspond to biochemical implementations for different combinations of weight signs. Note that in these cases *X*_1_ and *X*_2_ represent the same DNA strand but have independently varying concentrations, allowing them to be treated as two separate variables. In Figure 2B, the results with two different biases, *x*_0_ = 300 nM and 500 nM, are shown. As *X*_1_ and *X*_2_ are identical, the heat maps are symmetrical with respect to *x*_1_ = *x*_2_. The numerical simulations in the top row show that an increased bias moves the decision boundary toward the upper right because an increased *x*_0_ requires a greater concentration of fDNA to achieve the same output level. This trend was demonstrated experimentally, as shown in Figure 2B (bottom). In the decision boundary pattern presented in Figure 2C, *X*_1_ and *X*_2_ are qDNA corresponding to the negative weights and *X*_0_ is fDNA corresponding to the positive weight. An increased *x*_0_ moves the decision boundary to the upper right, which is observed in both numerical simulations and experimental demonstrations.

**Figure 2.**
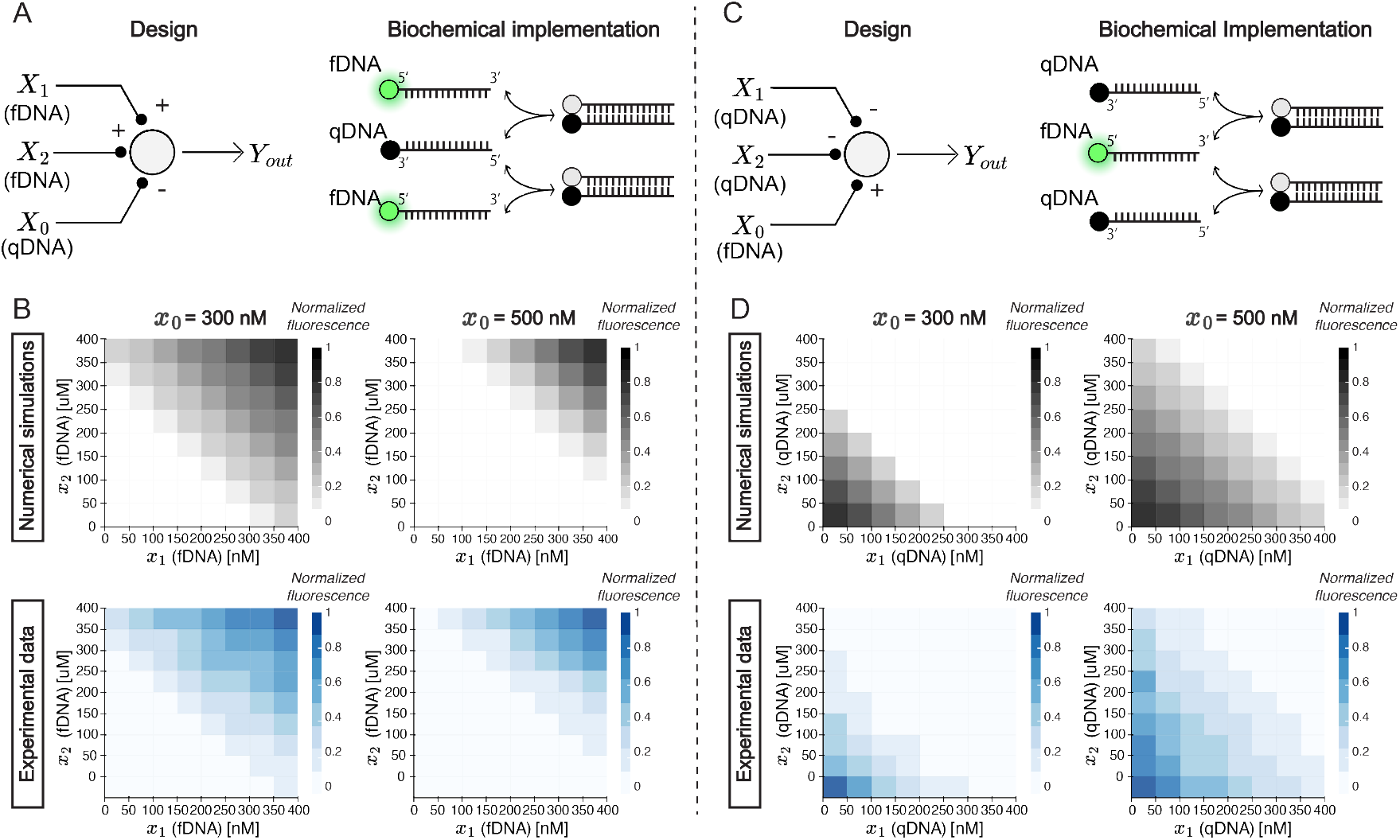
Different sign assignments for weights and a bias give various decision boundary patterns. In each panel, the top-left schematic shows an perceptron computation, which is implemented by a chemical reaction system shown in the top-right schematic diagram. The middle gray scale heat map shows numerical simulation results, and the bottm blue heat map shows experimental results. (A) Assigning two fDNA concentrations to inputs (x_1_ and x_2_, which are individually varied) and one qDNA concentration (x_0_) to bias implements positive weights and a negative bias. (B) Assigning two qDNA concentrations to inputs (x_1_ and x_2_, which are individually varied) and one fDNA concentration to bias (x_0_) implements positive weights and a negative bias.

Altogether, the experimental results show good agreement with the numerical simulations and demonstrate that the DNA hybridization-based system is well-suited for rapid prototyping of the sequestration-based linear classifier, thanks to its simplicity and programmability. However, it does not offer a practical way of weight tuning. One feasible way of weight tuning in the DNA hybridization-based system is adding dummy fDNA or dummy qDNA to decrease effective concentrations of regular fDNA or qDNA respectively. Dummy fDNA/qDNA has the identical sequence to the regular fDNA/qDNA, but the dummy fDNA/qDNA does not have a fluorophore/quencher. Therefore, adding dummy fDNA/qDNA dilute the effective concentrations of fDNA/qDNA. Supplementary Figure 3 shows experimental results and numerical simulations of DNA hybridization-based systems containing dummy qDNA (Panel A) or dummy fDNA (Panel B). While the results successfully demonstrate weight tuning in this system, this tunability is due to the unintended difference of the binding affinities of the regular binding (i.e., fDNA-qDNA) and binding with dummy strands (i.e., binding with dummy fDNA or dummy qDNA), which is most possibly derived from fluorophore-quencher interaction (Supplementary Section 2). The detailed mathematical analysis is provided in Section 2 of the Supplementary Information. In addition, the dynamic range of the tuned weights were small even if we added a significant fraction of the dummy strands. The lack of a practical way of weigh tuning is a disadvantage of the approach using only DNA hybridization reaction to form a sequestration reaction without any production of the signal species. It is not the case for some of the other DNA-based molecular classifiers. For example, Cherry and Qian reported a neural network based on DNA strand displacement, in which each perceptron executes a linear classification task [18]. Their system incorporates a double-stranded DNA complex for each weight assignment, which enables more drastic weight tuning. Hence, to overcome this limitation and realize a sequestration-based BNN with a wider range of tuned weights, in the following section, we introduce a molecular classifier based on TXTL reaction, which provides a more straightforward way of weight tuning.

**Figure 3.**
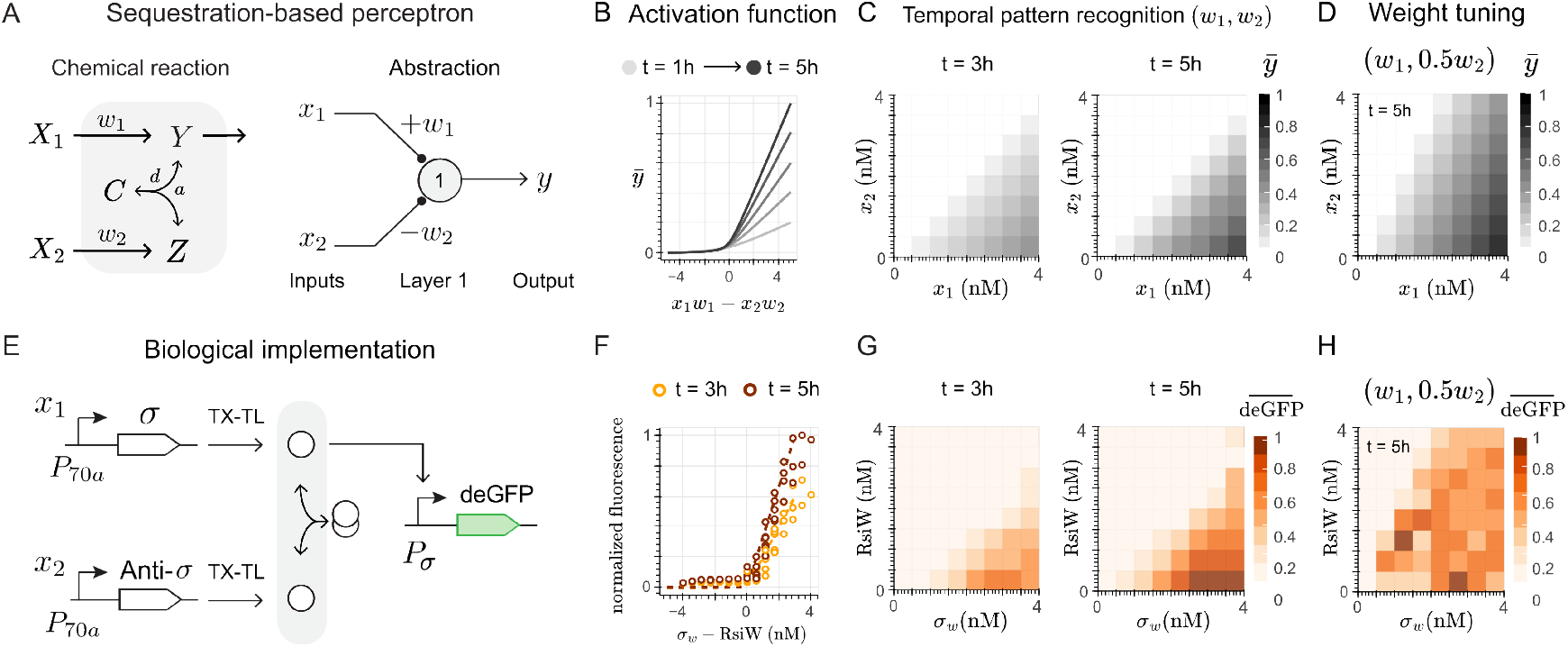
Molecular perceptron implementation using the molecular sequestration reaction between sigma (σ) and anti-sigma (anti-σ) factors in a cell-free (TXTL) system. (A) The chemical reactions describing the molecular perceptron (left), and its abstract schematics reflecting the signs of each weight (right). (B) The activation function obtained from the dynamics of the chemical reactions of Panel A, color-coded from t = 1 hour (light gray) to t = 5 (black).(C) The output of numerical simulations for w_1_ = w_2_ = 1 (/h) as heatmaps of the x_1_-x_2_ plane, highlighting the normalized value of output 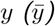 at two time points: t = 3 hours, and t = 5 hours. (D) The same heatmaps as (B) for t = 5 hours, but considering w_1_ = 1 and w_2_ = 0.5 (/h). (E) The biochemical realization of the molecular classifier using pairs of σ_w_/anti-σ factors; namely σ_28_/FlgM (blue) and σ_w_/RsiW (orange). (F) The experimental results as a scatter plot of the difference between the σ_w_ and RsiW plasmid concentrations with respect to the fluorescent readout at t = 3 hours (orange) and t = 5 hours (red). (G) The experimental results as heatmaps for the same time points as (C), considering 4 nM as the highest plasmid concentration for each species. (H) The experimental results for t = 5 hours, considering 4 nM of σ_w_ and 2 nM of RsiW as the highest plasmid concentrations. Normalizations for y at any time point for numerical simulations were done by considering the highest y value at t = 5, and the following kinetic parameters: a = 100 (/h/µM) and d = 1 (/h). For the experimental demonstrations, normalization of fluorescence intensity 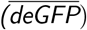 were done considering the highest fluorescence value at t = 5 hours.

### 2.2 Out-of-equilibrium linear classification using protein-level sequestration

Building on our previous work on molecular sequestration [32], we have constructed a TXTL-based linear classifier, as TXTL enables rapid prototyping in a cell-free environment and offers direct control over input concentrations (via titration), along with easy quantification of protein output [33]. Unlike the previous implementation (Figure 1A), here the inputs *X*_1_ and *X*_2_ are explicitly defined as chemical species, representing the number of plasmids encoding the production of proteins *Y* and *Z* at rates *w*_1_ and *w*_2_, respectively. We focus on the kinetic regime where resources in the TXTL mix are not depleted. Then, since ongoing chemical reactions do not reach steady-state in TXTL [34], we analyzed the output of the perceptron over time (*t*). First, under the assumption of a small dissociation constant, we derive an analytical approximation for the dynamics of species *Y*, given by

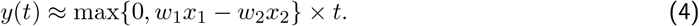

Here, *w*_1_ and *w*_2_ correspond to the perceptron’s weights, and as illustrated in the abstract schematics of Figure 3A, each weight is associated with a sign. In our molecular perceptron, *w*_1_ results in a positive action, and *w*_2_ in a negative one, as a result of the approximated subtraction. A detailed mathematical model is provided in Section 3 of the Supporting Information.

When analyzing the dynamics of *y*(*t*) over a simulation lasting 6 hours for varying inputs *x*_1_ and *x*_2_, and assuming production rates *w*_1_ = *w*_2_ = 1 (/h), the curve between the difference of the inputs and the normalized output 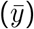 approaches a ReLU function-like behavior for a small dissociation constant value (*K* = 0.01). As suggested in equation (4) and confirmed in Figure 3B, this behavior holds across all time points (see also Supplementary Figure 4), suggesting that a decision boundary at *x*_1_ = *x*_2_ is present from the very beginning of the simulation (*t ≥* 1 hour). Figure 3C shows heatmaps in the *x*_1_-*x*_2_ plane constructed from these simulations at *t* = 3 and *t* = 5 hours. Both time points represent out-of-equilibrium conditions and exhibit a decision boundary, with the only difference being that the earlier time point shows a lower output intensity. We call this behavior temporal pattern recognition, as the linear classification function does not need to reach steady-state to appear.

**Figure 4.**
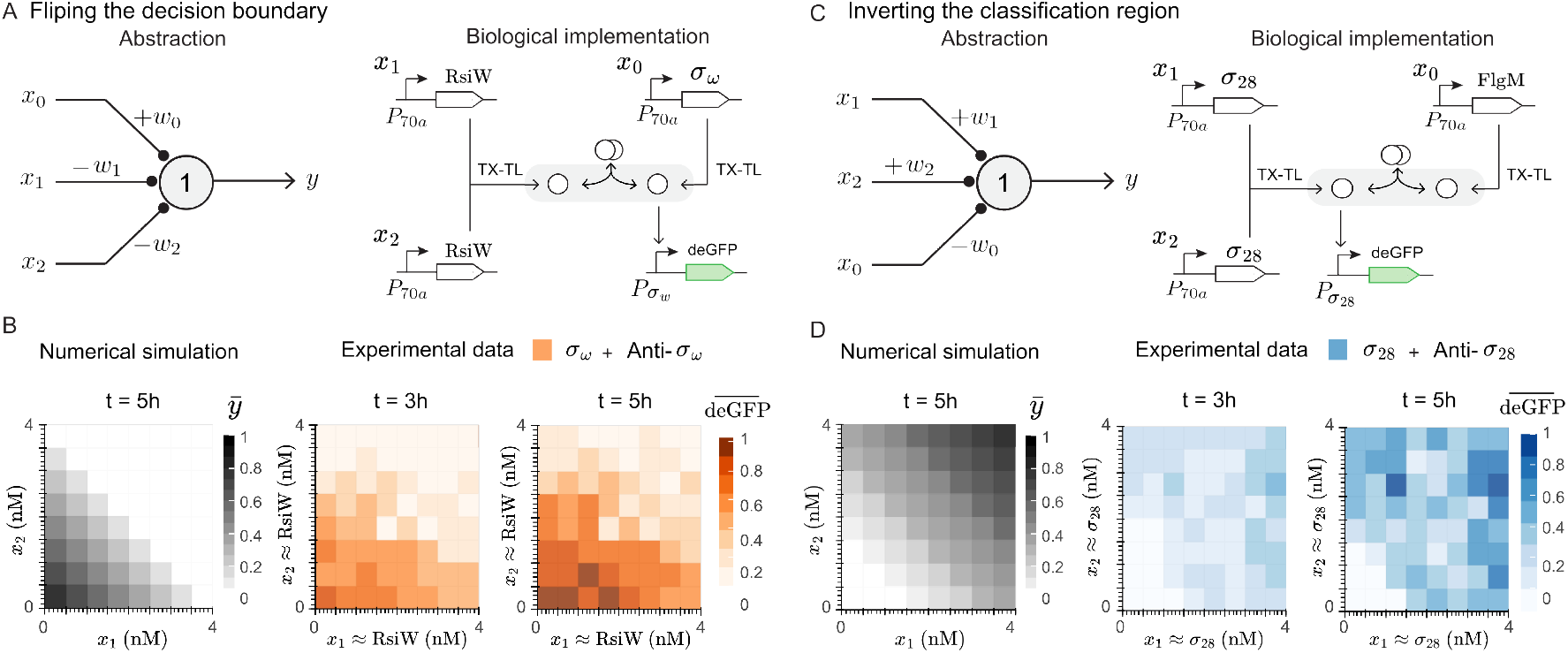
Adjusting the classification region by introducing a third input. (A) The design of a three-input linear classifier with a flipped decision boundary, with the classification region defined at the bottom-left corner of the boundary, which, at steady-state, exhibits negative weights (w_1_ and w_2_) associated with the inputs x_1_ and x_2_, and the positive weight (w_0_) associated with the third input x_0_ (left); along with its biochemical implementation based on the σ_w_/RsiW factors (right). (B) Comparison between the numerical simulations derived from the designed displayed in (A), showing the heatmap in the x_1_-x_2_ plane and the normalized y value 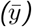 for t = 5 hours and w_1_ = w_2_ = 0.25 (/h) (left), along with the experimental results (right) for t = 3 and t = 5 hours. (C)A similar three-input classifier, but with the classification region defined at the upper-right corner relative to the decision boundary, with the positive weights (w_1_ and w_2_) associated with the inputs x_1_ and x_2_, and the negative weight (w_0_) associated with the third input x_0_ (left); along with the biochemical implementation using the σ_28_/FlgM factors (right). (D) Comparison between the numerical simulations for w_1_ = w_2_ = 1 (/h, left) with the experimental results for the same time points as Panel B. Normalizations for y at any time point for numerical simulations were done by considering the highest y value at t = 5, and the following kinetic parameters: a = 100 (/h/µM) and d = 1 (/h). For the experimental demonstrations, normalization of fluorescence intensity 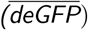 were done considering the highest fluorescence value at t = 5 hours.

Moreover, the mechanism for weight tuning is facilitated in our TXTL-based implementation: adjusting the production rate of either the *Y* or *Z* species allows for changes in the slope of the decision boundary, as illustrated in Figure 3D. As suggested by equation (4), reducing *w*_2_ to half of its nominal value increases the slope of the decision boundary, since a higher *x*_2_ concentration is now required for the *Z* and *Y* species to cancel each other out. In contrast, reducing *w*_1_ to half of its nominal value decreases the slope, as a lower *x*_2_ concentration is sufficient to achieve cancellation. Importantly, the temporal feature of pattern recognition is maintained in both cases.

For the biochemical realization of this molecular classifier, we used sigma (*σ*) factors - transcription factors that regulate protein expression. A *σ* factor binds to the RNA polymerase and activates transcription, whereas the anti-*σ* factor orthogonally binds to its corresponding *σ* factor and stops transcription by preventing the formation of the *σ*-RNA polymerase complex [35]. We considered two *σ*/anti-*σ* pairs for this implementation: the *σ*_*w*_ factor from *Bacillus subtilis* and its corresponding anti-*σ*_*w*_ (RsiW) [36], shown in orange throughout this section; and the *σ*_28_ factor from *Escherichia Coli* and its corresponding anti-*σ*_28_ named FlgM [37], shown in blue throughout this section. The reactions were carried out in a TXTL mix, with a constant concentration of DNA, to maintain the same resource consumption rate. To achieve this, we added a dummy plasmid that does not express any species involved in the sequestration reaction whenever the concentration of functional plasmids was reduced. Also, a constant concentration of a reporter gene (deGFP) with a corresponding *σ* factor-responsive promoter (P_*sω*_ and P_*s*28_) was used to monitor the dynamics of the *σ* factor (i.e., the *Y* species). An illustration of the biological implementation is shown in Figure 3E.

The experimental results in Figure 3F show that the *σ*_*w*_-based implementation behave as a linear classifier at every time point after *t* = 3 hours (see also Supplementary Figure 5), and are in good agreement with the temporal pattern recognition behavior observed in the numerical simulations (Figure 3G. See also Supplementary Figures 7 and 8). Prior to the *t* = 3 time point, although the decision boundary was theoretically predicted, the accumulation of fluorescent protein in the 2 *µ*L reaction well remained too low to be detected by the plate reader. Moreover, weight tuning was enabled by adjusting plasmid copy numbers, rather than modifying promoter strength (TX) or ribosome binding sites (TL). In particular, by diluting the RsiW plasmid concentration by half, we effectively increased the slope of the decision boundary (Figure 3H. See also Supplementary Figure 9 and 11). The same behavior was consistently observed for the *σ*_28_-based implementation: both the generation of a decision boundary (Supplementary Figures 6 to 8) and the feasibility of weight tuning, where halving the concentration of the *σ*_28_ plasmid resulted in a decrease in the slope of the decision boundary (Supplementary Figures 10 and 11). For both implementations, the weight-tuning operation does not rely on the system reaching steady-state.

**Figure 5.**
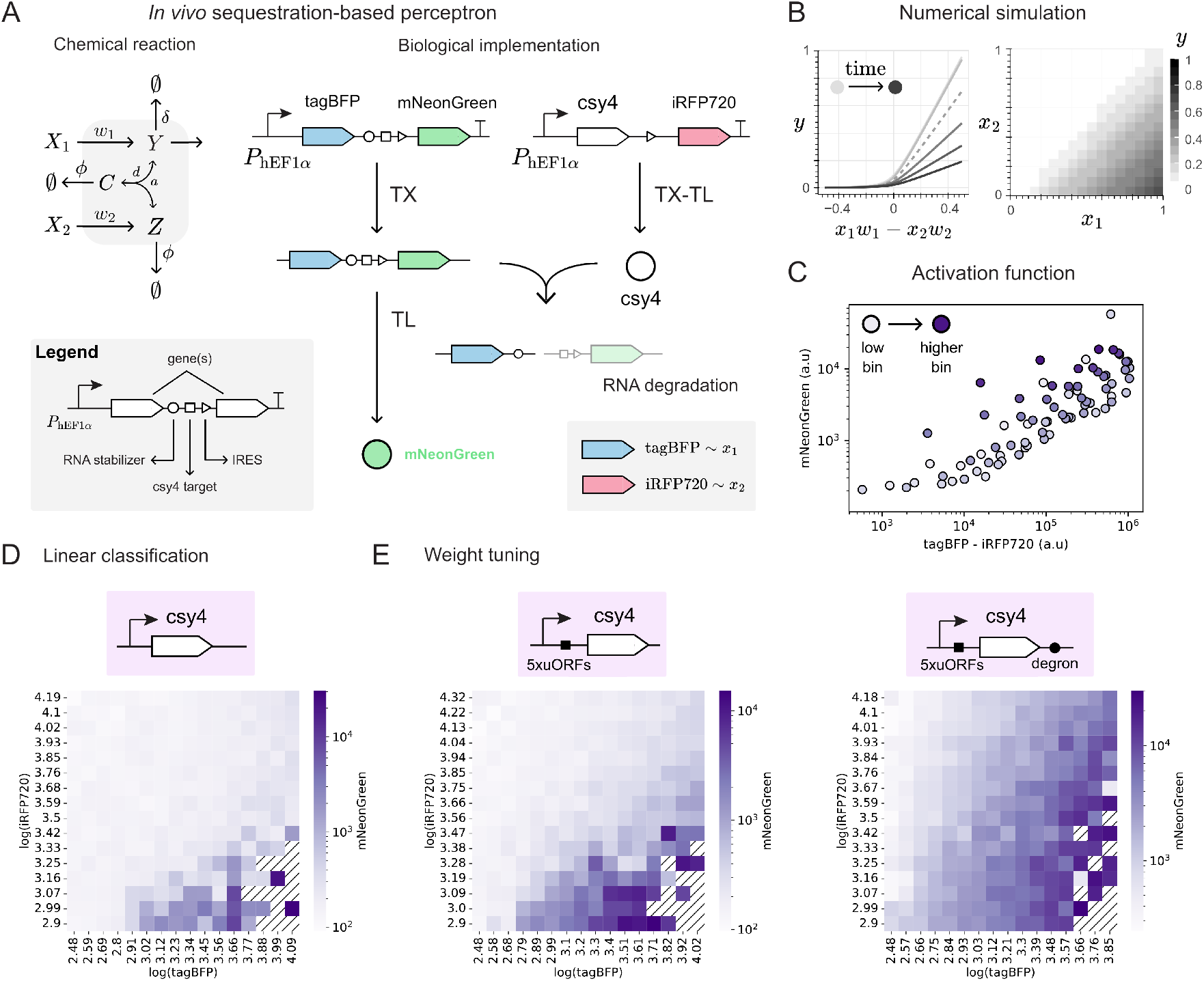
Endoribonuclease-based implementation of the molecular perceptron in mammalian cells. (A) The chemical reactions describing the molecular perceptron within a cellular context (left), and the biochemical realization using the Csy4 CRISPR-specific endoribonuclease (endoRNase). The sequestration reaction involves the Csy4 endoRNase and its target sequence (square) located upstream of the mNeonGreen gene. Both components of the reactions are encoded on independent plasmids, which co-express the tagBFP and iRFP720 fluorescent proteins as a proxy for the inputs x_1_ (i.e., mNeonGreen concentration) and x_2_ (i.e., Csy4 endoRNase concentration) using an IRES sequence (triangle). The mNeonGreen construct also includes an RNA stabilizer sequence (circle) to avoid degradation of the tagBFP protein upon cleavage of the Csy4 target sequence. (B) The activation function (left), obtained from the dynamics of the chemical reactions of (A), for 6 different time points and considering exponentially decaying inputs x_1_ and x_2_ with the shape x(t) = xe^−λt^. The third time point (dashed lines) was used to generate the heatmap on the x_1_-x_2_ plane (right). The following kinetic parameters were used for the numerical simulations: a = 100 (/h/µM), d = 1 (/h), w_1_ = w_2_ = 1 (/h), δ = 0.5 (/h), ϕ = 1 (/h); and for the inputs, x = 1 (µM) and λ = 1 (/h). (C) The experimental activation function, obtained with the median difference between tagBFP and iRFP720 versus the median mNeonGreen, calculated after dividing the data into bins, and color-coded by their location, from low (light purple) to high (dark purple). Black stripes indicate bins without a statistically significant number of cells. (D) and (E) show the experimental results as heatmaps, for the Csy4 endoRNase construct without any extra regulatory layer (C), with 5 repetitions of an upstream open reading frame (uORF) element (D, left), and with an additional downstream PEST degron sequence (D, right), exhibiting the linear classification and weight tuning features, respectively.

Furthermore, to explore the same type of decision boundaries shown in Figure 2 for the DNA hybridization system, we extended the model to include a new species *X*_0_ named bias, associated with a weight *w*_0_. To flip the decision boundary—mirrored along the vertical axis compared to Figure 3C—we relied onthe *σ*_*w*_/RsiW factor-based implementation. In the left schematics of Figure 4A, the weights of the *X*_1_ and *X*_2_ species are negative, as they correspond to two independent plasmids encoding the RsiW factor, which is expected to decrease the output (i.e., fluorescence intensity). In contrast, the weight of the *X*_0_ species is positive, as it represents the *σ*_*w*_ factor that drives deGFP expression. Under this framework, our numerical simulations, shown in Figure 4B (left), illustrate how the decision boundary can be flipped based on this design. For the experimental implementation, the plasmid encoding the bias species (i.e., *σ*_*w*_) was added at a concentration of 1 nM to the TXTL master mix. As seen in Figure 4B (right), the experimental results are in good agreement with the numerical simulations and exhibit the temporal pattern recognition feature, captured at *t* = 3 and *t* = 5 hours (see also Supplementary Figure 13).

The realization of the other decision boundary pattern is shown in Figure 4C, and was done based on the *σ*_28_/FlgM factors. To invert the classification region, so that fluorescence is observed in the uppermost right corner of the *x*_1_-*x*_2_ plane, we inverted the extended model’s inputs. Hence, now the species *X*_1_ and *X*_2_ are associated with a positive sign since they represent the production of the *σ*_28_ factor, whereas the bias species *X*_0_ is now associated with a negative sign, as it represents the production of FlgM. As in the previous analysis, we validated numerically this design for inverting the classification region, which can be observed in Figure 4D (left). For the experimental implementation, we also added the bias species (i.e., FlgM) to the TXTL master at a concentration of 1 nM. In Figure 4D (right), we observe our experiment results are in good agreement with the numerical simulations, as well, and display the temporal pattern recognition feature, captured also at *t* = 3 and *t* = 5 hours (see also Supplementary Figure 13).

### 2.3 In vivo implementation of a sequestration-based perceptron

Ultimately, we scale up the complexity of the molecular perceptron—not in its design, but in the biological context—by transfecting the plasmids encoding the sequestering species *Y* and *Z* into mammalian cells. The mathematical model that describes this implementation is quite similar to that of the TXTL-based implementation, with the difference that the species *Y* and *Z* have a “degradation” component to account for their dilution due to cell division and active protein degradation (Figure 5A, left). Following the small dissociation constant assumption, we can analytically derive the approximation for the dynamics of the *Y* species, given by

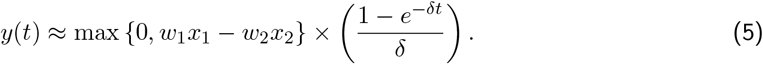

Here, *w*_1_ and *w*_2_ correspond to the perceptron’s weights, and *δ* represents the degradation rate of the species *Y*. A detailed mathematical model is provided in Section 4 of the Supporting Information.

For the biochemical realization of this in vivo molecular classifier, we rely on the reaction between a CRISPR-specific endoribonuclease (endoRNase) and its target sequence, which leads to RNA degradation [38]. We use the Csy4 endoRNase that forms a complex with its target sequence, which we added upstream of a fluorescent protein gene (mNeonGreen), and facilitates the degradation of the RNA transcript (Figure 5A, right). Since the association rate between the Csy4 and its target is strong [39], the binding between both species can be approximated as molecular sequestration in a fast sequestration regime, and so the system’s dynamics can be described by the same mathematical model as that of the abstraction presented in Figure 5A.

Importantly, we decided to take a high-throughput approach to characterize our design in a single experiment, and hence rely on the poly-transfection method proposed in [40]. By encoding the Csy4 endoRNase and the mNeonGreen gene with the target sequence in separate plasmids, and mixing each one independently with the transfection reagents—rather than mixing the plasmids before adding the reagents—we can generate a variety of plasmid copy numbers per cell, resulting in different stoichiometries between the two species across the population. Moreover, we co-express a fluorescent reporter protein on each plasmid using an internal ribosomal entry site (IRES, see Supplementary Figure 15), so its readout will serve as a proxy for the number of plasmid copies and, thus, as an estimate of the inputs. Specifically, we use tagBFP as a proxy for the input *x*_1_, and iRFP720 as a proxy for the input *x*_2_. Ultimately, to prevent the action of the Csy4 endoRNase from affecting the expression of tagBFP, we included a sequence downstream of the gene that encodes a triple-helix RNA-stabilizing structure from the MALAT1 RNA that reduces RNA degradation (cf. [38], and see Supplementary Figure 16).

Moreover, since the design relies on post-transcriptional regulation, we considered different degradation rates for each species to account for the differences in half-lives between the mNeonGreen mRNA (degradation described by *δ*) and the Csy4 endoRNase (degradation described by *ϕ*). Moreover, because poly-transfection is a transient transfection method, the system does not reach steady-state, as each plasmid is diluted over time with each cell division. We account for these details in our numerical simulations, which show the ReLU-like activation function and the decision boundary in the heatmap on the *x*_1_-*x*_2_ plane (Figure 5B), as suggested by equation (5)(see also Supplementary Figure 17), despite the out-of-equilibrium conditions and the time-decaying nature of the inputs.

After implementing the system and measuring between 80,000 and 110,000 HEK-293 cells per sample using flow cytometry, the experimental results validate the endoRNase-based implementation as an in vivo molecular perceptron. Following pre-processing and dividing the data into a 16×16 grid (Supplementary Figure 18), the scatter plot of the difference between the median tagBFP and iRFP720 versus the median mNeonGreen fluorescence, calculated per bin and shown in Figure 5D, closely resembles the ReLU activation function, compared with Figure 5B (see also Supplementary Figure 19). Moreover, since we used fluorescent reporter proteins to estimate the inputs *x*_1_ (with tagBFP) and *x*_2_ (with iRFP720), we identified the subset of data that maximizes the sampled area, ensuring that, after 16×16 binning, each bin contains a statistically significant number of cells (see Supplementary Figure 20). This enabled us to reconstruct the heatmaps in the *x*_1_-*x*_2_ plane, revealing a linear decision boundary (Figure 5C; see also Supplementary Figure 21).

Furthermore, the mechanism for weight tuning is similar to that of the TXTL-based implementation: adjusting the production rate of either the *Y* (i.e., mNeonGreen mRNA) or the *Z* (i.e., Csy4 endoRNase) species, which was confirmed for this implementation using numerical simulations (Supplementary Figure 22). Then, to experimentally increase the slope of the decision boundary, particularly by reducing the production rate of the Csy4 endoRNase (i.e., *w*_2_), we used 5 repetitions of an upstream open reading frame (uORF) sequence to reduce the translational rate of the Csy4 endoRNase (Supplementary Figure 23), which indeed resulted in the increased slope of the decision boundary, observed when comparing the circuit without uORFs (Figure 5D) and with the repetitions (Figure 5E, left. See also Supplementary Figure 25).

Ultimately, as an alternative strategy to modify the decision boundary, we explore fine-tuning the degradation rate. Ideally, for a very small dissociation constant (*K →* 0), equation (5) suggests that the output of the perceptron is decoupled from the degradation of species *Z*, as *y*(*t*) no longer depends on *ϕ*. However, numerical simulations show that when this assumption is relaxed, the degradation of *Z* (i.e., the Csy4 endoRNase) does influence the slope of the decision boundary (see Supplementary Figure 22). Intuitively, increasing the degradation of *Z* reduces the probability of sequestration. This means more of species *Y* is needed to cancel out species *Z*, which in turn increases the slope of the decision boundary. To achieve this effect experimentally, we further reduce the effective concentration of Csy4, beyond the five uORF copies, by attaching a PEST degron—a short peptide that promotes rapid protein degradation and has been characterized previously [34]. By targeting the Csy4 endoRNase gene with this degron, we observe an increase in the slope of the decision boundary (Figure 5E, right. See also Supplementary Figure 26).

## 3 Discussion

In this work, we experimentally demonstrated sequestration-based perceptrons at several levels of systems complexity, starting with a DNA-only system, progressing through a cell-free TX-TL system and scaling up to living cells. As the simplest realization, DNA hybridization provides a programmable platform for implementing a sequestration-based linear classifier at steady-state. Each complementary DNA strand is associated with a signed input, and experiments show that a linear decision boundary is generated, in good agreement with numerical simulations. However, adjusting the slope of this decision boundary relies on additional DNA strands (i.e., “dummy” strands), which may limit its scalability. In response to this limitation, we proposed the TXTL system as it offered more direct control over the perceptron’s weights by adjusting the plasmid copy number. This setup also enabled us to experimentally test the out-of-equilibrium behavior predicted by the mathematical model.

An 8×8 input grid was sufficient to capture the behavior of a linear decision boundary in both cell-free implementations. The use of a liquid handler enabled rapid and reproducible grid preparation, facilitating high-throughput prototyping. For the DNA-based system, this setup resulted in low measurement noise, as reflected in the close match between simulated and experimental heatmaps (Figure 1-D). In the TXTL system, the reaction was initiated by manually adding the mixture to the plate, followed by kinetic measurements in a microplate reader. While this introduced higher variability (Figure 3-F to H), the behavior of the linear classifier remained consistent. Interestingly, for the TXTL system, kinetic measurements from the *σ*_29_–FlgM perceptron showed a saturation in fluorescence signal after *t* = 5 hours, which was not due to sensor saturation (Supplementary Figures 6 and 10). Previous studies on TXTL systems derived from cell lysates suggest that this plateau results from resource depletion—such as ATP, amino acids, or carbon sources [34]. Although the time points at *t* = 3 and *t* = 5 hours are representative of the temporal pattern recognition behavior of the TXTL-based perceptron, a fed-batch setup could be used to extend the measurement window if needed.

Ultimately, we tested the temporal pattern recognition feature of the sequestration-based neural networks in mammalian cells, allowing us to explore how the system behaves under biological constraints. In particular, we focused on time-varying inputs—modeled as exponentially decaying since we used a transient transfection method—and on the dilution of chemical species due to cell division. As predicted by numerical simulations, the implementation of the sequestration reaction using an endoribonuclease-based design showed a decision boundary, whose slope could be changed by adjusting production rates, similar to the TXTL system. We highlight the use of the poly-transfection method as a “one-pot” assay that enabled us to characterize this implementation in a single flow cytometry experiment. Nevertheless, we acknowledge the limitation of this method in terms of the noisy measurements used as a proxy for production rates, though the resolution was sufficient to observe the decision boundary. A more rigorous quantification and increased resolution toward real-world applications may require calibration [41], as well as more sophisticated processing algorithms to deconvolve the network’s decision boundary from stochastic fluorescent readouts [42].

In summary, we systematically characterized different implementations of the sequestration-based perceptron, demonstrating that the network operates out-of-equilibrium, as decision boundaries can be reconstructed from snapshots at specific time points. To guide the design and interpretation of experiments, we relied on low-dimensional mathematical models based on ordinary differential equations. Although these models are not intended for quantitative predictions, they capture key features of the network that were validated experimentally, including the conditions required to compute ReLU-like responses, the tunability of decision boundaries, and the feasibility of out-of-equilibrium computation. We highlight the value of these models for circuit characterization, as they support both experimental planning and troubleshooting, as illustrated in Supplementary Figure 14. As future work, we aim to scale up these designs by interconnecting individual perceptrons into layered architectures, enabling multi-layer networks that compute non-linear decision boundaries and support more complex decision-making tasks [29]. We also plan to extend the framework by increasing input dimensionality (*n >* 2) and exploring alternative biochemical implementations based on phosphorylation [43], enzymatic catalysis [44], and competitive binding [45], each offering distinct trade-offs in programmability, tunability, and scalability.

## 4 Methods

### 4.1 Mathematical modeling

The complete set of Ordinary Differential Equations (ODEs) describing each reaction (i.e., DNA hybridization, TXTL and in vivo) and its features (e.g., activation function, weight tuning mechanism) is available in the Supplementary Information (Sections 1 to 4). The activation functions and heatmaps labeled as “simulations” presented in this work were generated by numerically integrating the ODE models using the *integrate*.*odeint* function from the *scipy* package in Python. Initial conditions were set to zero in all cases, while parameters for each simulation are described in each figure’s caption.

### 4.2 DNA hybridization reactions

Supplementary Table 2 lists the DNA oligos for the molecular classifier using the DNA hybridization system. All the DNA oligos are purchased from IDT. All the sequences are adapted from the genelet sequences designed by Schaffter et al [46]. fDNA is a partially double-stranded DNA composed to two single-stranded DNAs, referred to as fDNA-nt and fDNA-t. and fDNA is obtained by annealing fDNA-t and fNDA-nt in 1X NEB RNAPol Reaction Buffer (held at 90^*°*^C for 5 minutes then cooled down to 20^*°*^C at −1^*°*^C/min). Supplementary Figure 1 shows a detailed schematic diagram of the DNA hybridization system. We prepared samples with individually varied fDNA concentration (0 - 400 nM) and qDNA (50 - 400 nM) concentration with steps of 50 nM, using an Echo 550 liquid handler (Beckman Coulter). For the liquid handling operation, Echo qualified 384-well source plates (PP-0200, Labcyte) were used, and 96-well Clear V-Bottom plates (3357, Corning Inc.) were used as destination plates with 96 round well microplate storage mats (3080, Costar). The total volume of each sample was 2 *µ*L, which include fDNA and qDNA at specified concentrations, and 1X NEB RNAPol Reaction Buffer. After preparing samples on the desination plate, the plate was incubated at 37^*°*^C for 1 hour (except for E1-004 in Table 5). After the incubation, the destination plate is loaded into a microplate reader (Synergy H1, Biotek) to obtain fluorescence signal at excitation wavelength of 539 nm and emission wavelength of 569 nm at 37^*°*^C.

### 4.3 TXTL reactions

The cell-free expression system used in this work is myTXTL from Arbor Biosciences. Linear DNA, purchased from IDT, was used for all reactions. Sequences are available in the Supplementary Data. Samples were prepared with individually varying concentrations of sigma (*σ*) factor plasmid and anti-sigma (anti-*σ*) factor plasmid, ranging from 0 to 4 nM for both, in steps of 0.57 nM, unless stated otherwise. These samples were prepared using the Echo 550 liquid handler (Beckman Coulter). For liquid handling, Echo qualified 384-well source plates (PP-0200, Labcyte) and destination 96-well V-bottom plates (3357, Corning Costar) with 96 round well microplate storage mats (3080, Costar) were used. The total volume of each sample was 2 *µ*L, which included a reporter plasmid at 4 nM, the *σ* and anti-*σ* factor plasmids at the specified concentrations, myTXTL Master Mix (507096, Arbor Biosciences), and myTXTL GamS Nuclease Inhibitor Protein (501038, Arbor Biosciences). After preparing the samples on the destination plate, the plate was either loaded into a microplate reader (Synergy H1, Biotek) and incubated at 30^*°*^C for 6 hours, or incubated in a separate incubator at the same temperature for 6 hours, with point measurements taken in the microplate reader at specified hours. In both cases, fluorescence signals were measured at an excitation wavelength of 479 nm and an emission wavelength of 520 nm.

### 4.4 Plasmid cloning and circuit design

Plasmids were constructed using the Golden Gate strategy described in [47, 48]. Individual parts—promoters, 5’-UTRs, coding sequences, 3’-UTRs, and terminators—referred to as Level 0s (pL0s), were generated via PCR (Q5 polymerase, New England Biolabs) or ordered as gblocks (IDT), then cloned into pL0 backbones by digestion and ligation. These pL0s were assembled into transcriptional units, or Level 1s (pL1s), using Bas1 Golden Gate reactions (10–50 cycles between 16^*°*^C and 37^*°*^C, T4 DNA ligase; enzymes from New England Biolabs). Constructs were transformed into Stellar E. coli competent cells (Takara Bio), plated on LB agar (VWR), and cultured in TB medium (Sigma-Aldrich). Spectinomycin (100 *µ*g/*µ*L), kanamycin (50 *µ*g/mL) or X-Gal (40 *µ*g/*µ*L) were added to the plates or media, depending on plasmid selection markers. Plasmids were extracted using QIAprep Spin Miniprep or QIAGEN Plasmid Plus Midiprep kits, and sequences were confirmed by Sanger sequencing at Genewiz. All plasmid and backbone files are available in Supplementary Data. Sequences were designed and annotated in Benchling.

### 4.5 Cell culture

HEK-293 cells (Lenti-X, Takara Bio) were maintained in DMEM containing 4.5 g/L glucose, L-glutamine, and sodium pyruvate (Corning) supplemented with 10% fetal bovine serum (FBS, obtained from VWR). Cells were grown in a humidified incubator at 37^*°*^C and 5% CO_2_.

### 4.6 Transfections

Transfections were conducted in a 24-well plate (Coaster) using FuGENE HD (Promega), according to the manufacturer’s protocol but with modifications for poly-transfection detailed in [40, 49]. Essentially, one plasmid (e.g., mNeonGreen construct in Figure 5) was mixed in one tube with 0.5 *µ*l of transfection reagent, while the other plasmid (e.g., Csy4 endoRNase construct in Figure 5) was mixed in another tube with 0.5 *µ*l of transfection reagent. DNA–lipid complexes were allowed to form, and both mixtures were then added simultaneously to the same well containing cells.

### 4.7 Flow cytometry

Sample preparation for flow cytometry was done following the exact same protocol detailed in [48], and collecting 75,000–120,000 cells per sample. For all experiments, samples were collected on a CytoFLEX LX flow cytometer (Beckman Coulter) equipped with a 488 nm laser with 525/40 nm filter for measuring tagBFP, 561 nm laser with 585/42 filter for measuring mNeonGreen, and 638 nm laser with 660/10 nm filter for measuring iRFP720. 500-2000 events/s were collected either in tubes via the collection port. Compensation was not applied since fluorescence spectra was proven orthogonal [40].

### 4.8 Flow cytometry data analysis

Flow cytometry data were analyzed using a custom pipeline in Python v3.10. Individual FCS files were processed with FlowCal [50]. Saturating events were removed, followed by density gating (first gate: 0.6; second gate: 0.8) using FlowCal’s built-in functions *gate*.*high_low* and *gate*.*density2d*. Data were then filtered based on a minimum fluorescence threshold in the tagBFP and iRFP720 channels, using baseline fluorescence from a control (no-plasmid) sample. To estimate the activation function, the filtered dataset was binned into a 16×16 grid according to tagBFP and iRFP720 fluorescence levels (Supplementary Figure 18). Median fluorescence values for tagBFP, iRFP720, and mNeonGreen were calculated for each bin, and a scatterplot was generated where each point represents the difference between median tagBFP and iRFP720 versus the corresponding median mNeonGreen.

To construct the heatmap in the *x*_1_–*x*_2_ plane, we first filtered the data using the aforementioned fluorescence thresholds. For notation, the measurement’s dynamic range was defined as the difference between the lowest and highest fluorescence values in the filtered data. We then defined a series of rectangular regions covering increasing percentages of this dynamic range. For each rectangular region, the data were divided into a 16×16 grid, and the number of cells per bin was calculated. To ensure each bin contained a statistically significant number of events, we set a minimum bin count threshold based on a Poisson model and a one-sided 95% confidence bound. The region that maximized the number of bins exceeding this threshold was selected to ensure that it contained sufficient events per bin. The final heatmap was then reconstructed by calculating the median mNeonGreen fluorescence for each bin. An illustrative example of this method, used to construct Panel D of Figure 5, is available in Supplementary Figure 20.

## Acknowledgments

We thank Nika Shakiba, Karmella Haynes, Jean-Baptiste Lugagne, Jeremie Marlhens, Enoch Antwi, Ross Jones, and Rongrong Du for their critical feedback. E.N. acknowledges support from National Science Foundation through award NSF-CCF 2107483.

## Notes

### Competing Interest Statement

The authors have declared no competing interest.

