## Supplementary Information for "Sequestration-based neural networks that operate out of equilibrium"

[1]Department of Mechanical and Aerospace Engineering, University of California at Los Angeles, Los Angeles, CA, USA
[2]Computational Biology Department, Carnegie Mellon University, Pittsburgh, PA, USA
[3]Department of Chemistry, Imperial College London, London, UK
[4]Biomedical Engineering, Boston University, Boston, MA, USA
[5]Biological Design Center, Boston University, Boston, MA, USA
[6]Department of Biochemistry and Systems Medicine, Juntendo University, Tokyo, Japan
[7]Department of Biochemistry and Molecular Biology, Medical University of Lublin, Lublin, Poland
[8]Department of Molecular Biology, Institute of Biochemistry, Faculty of Biology, University of Warsaw, Miecznikowa 1, Warsaw, Poland
[9]The Translational Plant Sciences Center, Virginia Tech, Blacksburg, Virginia, USA
[10]Fischell Department of Bioengineering, University of Maryland, College Park, Maryland, USA
[11]Department of Mechanical Engineering, Stanford University, Stanford, CA, USA
[12]Department of Cancer Immunology and Virology, Dana-Farber Cancer Institute, Boston, MA, USA
[13] Department of Immunology, Harvard Medical School, Boston, MA, USA
[+]Synthetic Biology Summer School at Cold Spring Harbor 2024
[*]Equal contribution

May 16, 2025

**5 Experimental details of DNA hybridization system 15**

**6 Weight tuning by the addition of dummy DNA strands in the DNA hybridization system 16**

**7 Experimental details of TXTL system 20**

In this manuscript, we use capital letters to denote chemical species (e.g., $Y$) and lowercase letters for their concentrations (e.g., $y$). We consider that all variables depend on time but omit the explicit notation for simplicity. For instance, we write $y$ instead of $y(t)$

### 1 DNA hybridization

First, we develop the mathematical model of the simplest realization of a sequestration-based perceptron using DNA hybridization. This implementation consists of two single DNA strands: a sense strand $Y$ and anti-sense strand $Z$. Both can form a complex at a rate constant $a$, and dissociate at a rate constant $d$. This interaction can be described by the following chemical reactions:

$$\begin{aligned} Y + Z &\xrightarrow{a} C, \\ C &\xrightarrow{d} Y + Z. \end{aligned} \tag{1}$$

Using the law of mass of action, we can write down the Ordinary Differential Equations (ODEs) that describes their dynamics,

$$\dot{y} = dc - ayz \tag{2}$$
$$\dot{z} = dc - ayz \tag{3}$$
$$\dot{c} = ayz - dc \tag{4}$$

with a mass conservation given by $y + c = x_1$, and $z + c = x_2$. where $x_1$ and $x_2$ are initial concentrations of $Y$ and $Z$, respectively.

#### 1.1 Steady State analysis

##### 1.1.1 Steady state of $y$

We can rewrite the equation (2) as

$$\dot{y} = d(x_1 - y) - ay(y - x_1 + x_2) \tag{5}$$

We can further find the steady-state solution of $y$ by making equation (5) equal to zero. We find

$$K(x_1 - \bar{y}) - \bar{y}(\bar{y} - x_1 + x_2) = 0, \tag{6}$$

where $K = d/a$ is the dissociation constant. This leads to a second order polynomial

$$\bar{y}^2 + A\bar{y} + B = 0,$$

where $A = -(x_1 - x_2) + K$, and $B = -Kx_1$. It admits a single positive real solution since $B$ is negative. Therefore,

$$\bar{y} = \frac{1}{2}\left(x_1 - x_2 - K + \sqrt{(x_1 - x_2 - K)^2 + 4Kx_1}\right) \tag{7}$$

We explore the regime of a small dissociated constant by defining $\xi_y = K/x_1 \to 0$, and finding the steady state of $\bar{y}$,

$$\lim_{\xi_y \to 0} \bar{y} = \frac{1}{2} \left\{ (x_1 - x_2) + |(x_1 - x_2)| \right\} \tag{8}$$

We can rewrite equation (8) as follows,

$$\lim_{\xi_y \to 0} \bar{y} = \max(0, x_1 - x_2) \tag{9}$$

###### 1.1.2 Steady state of $z$

We can rewrite the equation (3) as

$$\dot{z} \quad = \quad d(x_2 - z) - az(z - x_2 + x_1) \tag{10}$$

Following similar steps as Section 2.2.1, we can find the steady state value of $z$ as

$$\bar{z} = \frac{1}{2} \left( x_2 - x_1 - K + \sqrt{(x_2 - x_1 - K)^2 + 4Kx_2} \right) \tag{11}$$

We explore the regime of a small dissociation constant by defining $\xi_z = K/x_2 \to 0$, and finding the steady state of $\bar{z}$,

$$\lim_{\xi_z \to 0} \bar{z} = \max(0, x_2 - x_1) \tag{12}$$

###### 1.1.3 Steady state of $c$

We can rewrite the equation (4) as

$$\dot{c} \quad = \quad a(c - x_1)(c - x_2) - dc \tag{13}$$

Following similar steps as Section 2.2.1, we can find the steady state value of $c$ as

$$\bar{c} = \frac{1}{2} \left( x_2 + x_1 + K - \sqrt{(x_2 + x_1 + K)^2 - 4x_1x_2} \right) \tag{14}$$

We explore the regime of a small dissociation constant by defining $\xi_c = K/(x_1 + x_2) \to 0$ , and finding the steady state of $\bar{c}$,

$$\lim_{\xi_c \to 0} \bar{c} = \frac{1}{2} \left( x_2 + x_1 - \sqrt{(x_2 + x_1)^2 - 4x_1x_2} \right) = \frac{1}{2} (x_2 + x_1 - |x_2 - x_1|) \tag{15}$$

We can rewrite equation (15) as follows

$$\lim_{\xi_c \to 0} \bar{c} = \min(x_1, x_2) \tag{16}$$

#### 2 DNA hybridization with dummy strands

Here we modify the mathematical model of the DNA hybridization system in the previous section to incorporate the effect of dummy DNA strands, which do not have a fluorophore or quencher. Dummy fDNA, $Y_d$, has the identical sequences to $Y$, and it binds either $Z$ or $Z_d$ to form complexes $C$ or $C_{dy}$ respectively, but concentrations of $Y_d$ is not considered as output because it does not fluoresce. Similarly, dummy qDNA, $Z_d$, binds either $Y$ or $Y_d$ to form complexes $C$ or $C_{dz}$ respectively, and $Y$ and $C_{dz}$ are considered as outputs, because $Z_d$ does not inactivate $Y$ in the complex $C_z$. This interaction can be described by the following chemical reactions:

$$
\begin{aligned}
Y + Z &\xrightarrow{a} C, \\
C &\xrightarrow{d} Y + Z. \\
Y_d + Z &\xrightarrow{a_{dy}} C_{dy}, \\
C_{dy} &\xrightarrow{d_{dy}} Y_d + Z. \\
Y + Z_d &\xrightarrow{a_{dz}} C_{dz}, \\
C_{dz} &\xrightarrow{d_{dz}} Y + Z_d. \\
Y_d + Z_d &\xrightarrow{a_{dyz}} C_{dyz}, \\
C_{dyz} &\xrightarrow{d_{dyz}} Y_d + Z_d.
\end{aligned}
\tag{17}
$$

In practice, we do not add both $Y_d$ and $Z_d$ in the same sample because, if both are added, they cancel each other's dilution effects.

#### 2.1 With dummy fDNA

Here we assume that the system has dummy fDNA, but does not have dummy qDNA. Using the law of mass action, we can write down the modified ODEs as follows.

$$\dot{y} = dc - ayz \tag{18}$$

$$\dot{z} = dc + d_d c_d - ayz - a_d y_d z \tag{19}$$

$$\dot{c} = ayz - dc \tag{20}$$

$$\dot{y}_d = d_y c_d - a_d y_d z \tag{21}$$

$$\dot{c}_d = a_d y_d z - d_d c_d \tag{22}$$

Because we have only one dummy strand, $Y_d$, the subscripts of the constants and variables involving $Y_d$ are updated (e.g., $d_{dy} \to d_d$) to simplify the notations. The above ODEs satisfy mass conservations given by

$$y + c = \alpha x_1 \tag{23}$$

$$y_d + c_d = (1-\alpha)x_1 \tag{24}$$

$$z + c + c_d = x_2 \tag{25}$$

where $x_1$ and $x_2$ are inputs. At steady state,

$$\dot{c} = a\bar{y}\bar{z} - d\bar{c} = 0 \quad \Rightarrow \quad \bar{c} = \frac{\bar{y}\bar{z}}{K} \tag{26}$$

$$\dot{c}_d = a_d \bar{y}_d \bar{z} - d_d \bar{c}_d = 0 \quad \Rightarrow \quad \bar{c}_d = \frac{\bar{y}_d \bar{z}}{K'} \tag{27}$$

where $K = d/a$ and $K' = d_d/a_d$. Substituting these into the equation 25 and rearranging with respect of $z$, we obtain

$$\bar{z} = \frac{x_2}{1 + \bar{y}/K + \bar{y}_d/K'} \tag{28}$$

By using the equation 26, 27, and 28, the equations 23 and 24 can be rearranged as follows.

$$\bar{y} + \frac{\bar{y}/K}{1+\bar{y}/K+\bar{y_d}/K'}x_2 = \alpha x_1 \tag{29}$$

$$\bar{y_d} + \frac{\bar{y_d}/K'}{1+\bar{y}/K+\bar{y_d}/K'}x_2 = (1-\alpha)x_1 \tag{30}$$

We explore the regime of small dissociation constants, i.e., $K/\bar{y},\ K'/\bar{y_d} \to 0$, and rewrite the equations 29 and 30 as follows.

$$\bar{y} + \frac{\bar{y}/K}{\bar{y}/K+\bar{y_d}/K'}x_2 = \alpha x_1 \tag{31}$$

$$\bar{y_d} + \frac{\bar{y_d}/K'}{\bar{y}/K+\bar{y_d}/K'}x_2 = (1-\alpha)x_1 \tag{32}$$

Note that the assumption, $K/\bar{y},\ K'/\bar{y_d} \to 0$, is not valid if $x_1 \leq x_2$, because it gives a small value of $\bar{y}$ and $\bar{y_d}$ (i.e., most of fDNA and dummy fDNA is sequestered), leading to relatively large values of $K/\bar{y}$ and $K'/\bar{y_d}$. By adding both sides of the equations 31 and 32. we obtain

$$\bar{y} + \bar{y_d} = x_1 - x_2 \tag{33}$$

Using this equation, the equation 31 can rewritten as follows.

$$(K'-K)\bar{y}^2 + [Kx_1 + (\alpha x_1 - x_2)(K-K')]\bar{y} - \alpha K x_1(x_1-x_2) = 0 \tag{34}$$

In the system with dummy fDNA, the output is $\bar{y}$. Thus, in the following, we derive $\bar{y}$ for two cases.

###### 2.1.1 Case 1: $K' = K$

The equation 34 can be rewritten as

$$\bar{y} = \alpha(x_1 - x_2) \tag{35}$$

In this case, the decision boundary does not change, because each data point $\bar{y}(x_1, x_2)$ is just scaled by the same constant $\alpha$. The equation 35 is rewritten as follows

$$x_2 = x_1 - \bar{y}/\alpha \tag{36}$$

and we find $\alpha$ does not change the slope of $x_2$ as a function of $x_1$, which means the addition of the dummy fDNA does not tune the weight.

###### 2.1.2 Case 2 $K' \neq K$

Solving the equation 34 and choosing the positive root gives

$$\bar{y} = \frac{-[Kx_1 - \Delta K(\alpha x_1 - x_2)] + \sqrt{[Kx_1 - \Delta K(\alpha x_1 - x_2)]^2 + 4\alpha\Delta K K x_1(x_1-x_2)}}{2\Delta K} \tag{37}$$

where, $\Delta K = K' - K$. We can rearrange the equation 34 with respect of $x_2$ as follows.

$$x_2 = \frac{\alpha K x_1^2 + \alpha\Delta K \bar{y} x_1}{\alpha K x_1 + \Delta K \bar{y}} - \frac{\Delta K \bar{y}^2 + K x_1 \bar{y}}{\alpha K x_1 + \Delta K \bar{y}} \tag{38}$$

In this case, $x_2$ is a non-linear function of $x_1$, and $\alpha$ affects both terms in the equation 38. Although it is not a linear decision boundary, this analysis shows that the addition of the dummy fDNA provides additional tunability for the decision boundary of the DNA-hybridization system.

#### 2.2 With dummy qDNA

##### 2.2.1 Steady state of $y + c_{dz}$

Using the law of mass action, we can write the modified ODEs for the system with dummy qDNA as follows.

$$\dot{y} = dc + d_d c_d - ayz - a_d y z_d \quad (39)$$
$$\dot{z} = dc - ayz \quad (40)$$
$$\dot{c} = ayz - dc \quad (41)$$
$$\dot{z}_d = d_y c_d - a_d y z_d \quad (42)$$
$$\dot{c}_d = a_d y_d z - d_d c_d \quad (43)$$

with mass conservations given by

$$y + c + c_d = x_1 \quad (44)$$
$$z + c = \beta x_2 \quad (45)$$
$$z_d + c_d = (1-\beta)x_2 \quad (46)$$

At steady state,

$$\dot{c} = a\bar{y}\bar{z} - d\bar{c} = 0 \quad \Rightarrow \quad \bar{c} = \frac{\bar{y}\bar{z}}{K} \quad (47)$$
$$\dot{c}_d = a_d \bar{y}\bar{z}_d - d_d \bar{c}_d = 0 \quad \Rightarrow \quad \bar{c}_d = \frac{\bar{y}\bar{z}_d}{K'} \quad (48)$$

By substituting these into the equation 45 and 46, we obtain

$$\bar{z} = \frac{\beta x_2}{1 + \bar{y}/K} \quad (49)$$
$$\bar{z}_d = \frac{(1-\beta)x_2}{1 + \bar{y}/K'} \quad (50)$$

By using the equation 47, 48, 49, and 50, the equation 44 can be written as

$$\bar{y} + \frac{\bar{y}}{K}\frac{\beta x_2}{1 + \bar{y}/K} + \frac{\bar{y}}{K'}\frac{(1-\beta)x_2}{1 + \bar{y}/K'} = x_1 \quad (51)$$

Assuming $y/K,\ y/K' >> 1$ in the equation 51, the approximation of $\bar{y}$ can be obtained as follows.

$$\bar{y} \approx x_1 - x_2 \quad (52)$$

By substituting $\bar{y}$ into the equations 49,

$$\bar{z} = \frac{\beta x_2}{1 + (x_1 - x_2)/K} \quad (53)$$

From the equation 45,

$$\bar{c} = \beta x_2 - \bar{z} = \frac{\beta x_2 (x_1 - x_2)/K}{1 + (x_1 - x_2)/K'} \quad (54)$$

Substituting $\bar{c}$ into the equation 44 gives

$$\bar{y} + \bar{c}_d = x_1 - c = x_1 - \frac{\beta x_2 (x_1 - x_2)/K}{1 + (x_1 - x_2)/K'} \quad (55)$$

In this system, $C_d$ emits fluorescence signal because $Z_d$ does not have a quencher. Therefore, we take $Y = \bar{y} + \bar{c_d}$ as the output. Then, we can rearrange the equation 55 as follows.

$$\beta x_2^2 + (Y - (1+\beta)x_1)x_2 + (x_1 - Y)(x_1 + K) = 0 \tag{56}$$

Assuming $x_1 >> K$, we obtain an approximate solution of $x_2$ as

$$\begin{aligned} x_2 &= \frac{(1+\beta)x_1 - Y \pm \sqrt{(Y-(1+\beta)x_1)^2 - 4\beta(x_1 - Y)x_1}}{2\beta} && (57) \\ &= \frac{(1+\beta)x_1 - Y \pm |Y - (1-\beta)x_1|}{2\beta} && (58) \end{aligned}$$

$Y - (1-\beta)x_1 = x_1 - c - (1-\beta)x_1 = \beta x_1 - c = \beta(x_1 - x_2) + z > 0$ Therefore,

$$x_2 = x_1, \frac{x_1}{\beta} - \frac{Y}{\beta} \tag{59}$$

With finite values of $x_1$ and $x_2$, $y > 0$. Therefore, $x_2 = x_1$ is not valid. Thus,

$$x_2 = \frac{x_1}{\beta} - \frac{Y}{\beta} \tag{60}$$

This equation represents the decision boundary of the system with dummy qDNA.

### 3 Modeling sequestration reactions in TX-TL

Next, we develop the mathematical model of a protein-based implementation of the perceptron using $\sigma$ and anti-$\sigma$ factors that exhibit sequestration dynamics. We consider the species $X_1$ and $X_2$ that represent the DNA sequences that encode the $\sigma$ (denoted by the $Y$ species) and anti-$\sigma$ factors (denoted by the $Z$ species), and produce them at a rate $w_1$ and $w_2$, respectively. The binding between both proteins generates a complex $C$ at a rate constant $a$, and dissociates at a rate constant $d$. This interaction can be described by the following chemical reactions:

$$\begin{aligned} X_1 &\xrightarrow{w_1} X_1 + Y, \\ X_2 &\xrightarrow{w_2} X_2 + Z, \\ Y + Z &\underset{d}{\overset{a}{\rightleftharpoons}} C. \end{aligned} \tag{61}$$

Assuming that the reactions occur in a well-steered volume, we can construct the ODEs that describe their dynamics following the law of mass action, resulting in

$$\begin{aligned} \dot{y} &= w_1 x_1 + dc - ayz && (62) \\ \dot{z} &= w_2 x_2 + dc - ayz && (63) \\ \dot{c} &= ayz - dc && (64) \end{aligned}$$

with a mass conservation given by $y^T = y + c$ and $z^T = z + c$.

#### 3.1 Temporal analysis

Eqs. (62)–(64) suggest that the protein-based implementation of the perceptron within a cell-free system does not reach steady-state. Hence, we explore the temporal dynamics of the reaction. Using the mass conservation equations, we can rewrite equations (62)-(64) as

$$\begin{aligned} \dot{y}^T &= w_1 x_1 && (65) \\ \dot{z}^T &= w_2 x_2 && (66) \\ \dot{c} &= a(c - y^T)(c - z^T) - dc && (67) \end{aligned}$$

We define the dissociation constant $K = d/a$ and rewrite Eq. (67) as follows,

$$(\frac{K}{d})\frac{d}{dt}c = (c - y^T)(c - z^T) - Kc \quad (68)$$

We explore the regime of a small dissociation constant by defining $\xi = K/[(x_1w_1 - x_2w_2)/d] \to 0$, and therefore

$$\lim_{\xi \to 0} c(t) = \min(y^T(t), z^T(t)) \quad (69)$$

In other words, $c(t)$ approaches either $y^T(t)$ or $z^T(t)$ when $\xi \to 0$, which is consistent with previous derivations [1]. Then, recalling the equations from mass conservation, and since equation (65) does not depend on $\xi$,

$$\lim_{\xi \to 0} y(t) = y^T(t) - \lim_{\xi \to 0} c = y^T - \min(y^T(t), z^T(t)) = \max(0, y^T(t) - z^T(t)) \quad (70)$$

We can further find the analytical solutions to Eq. (65) and Eq. (66) by integration. That is,

$$\begin{aligned} \int dy^T &= \int w_1 x_1 dt && (71) \\ \int dz^T &= \int w_2 x_2 dt && (72) \end{aligned}$$

Solving,

$$\begin{aligned} y^T(t) &= (w_1 x_1)t + c_1 && (73) \\ z^T(t) &= (w_2 x_2)t + c_2 && (74) \end{aligned}$$

where $c_1$ and $c_2$ depend on the initial conditions. That is, the total concentration of the $Y$ and $Z$ species at the beginning of the reaction ($t = 0$). Since the reaction is done in a cell free system where the $\sigma_w$/RsiW nor the $\sigma_{28}$/FlgM proteins are endogenously expressed, we have that both $y^T(t = 0)$ and $z^T(t = 0)$ are 0 $\mu$M. Then, by substitution in Eq. (70), we can find the expression of $y(t)$ in the regime of a small dissociation constant, given by

$$\lim_{\xi \to 0} y(t) = \max\{0, (w_1x_1 - w_2x_2)t\} \quad (75)$$

Similarly, recalling the equations from mass conservation, and since Eq. (66) does not depend on $\xi$,

$$\lim_{\xi \to 0} z(t) = z^T(t) - \lim_{\xi \to 0} c = z^T(t) - \min(y^T(t), z^T(t)) = \max(0, z^T(t) - y^T(t)) \quad (76)$$

From the analytical expressions found in Eqs. (73) and (74), and by subtitution in Eq. (76), we can find the expression of $z(t)$ in the regime of a small dissociation constant, given by

$$\lim_{\xi \to 0} z(t) = \max \{0, (w_2 x_2 - w_1 x_1)t\} \tag{77}$$

### 4 Modeling sequestration reactions in mammalian cells

The molecular sequestration reaction within a living cell can be modeled with similar chemical reactions as in equation (61) but considering degradation and dilution due to cell division of the species $Y$, $Z$ and $C$. In this work, we propose an RNA-protein implementation of the perceptron based on the sequestration reaction between the Csy4 endoribonuclease (endoRNase) and its target sequence located at the 5' end of the mNeonGreen gene, which is cleaved after transcription (see Figure 5 in the Main Text for schematics). For notation, the $Y$ species represents the mNeonGreen mRNA, the $Z$ species represents the Csy4 endoRNase, and the $C$ species represents the RNA-protein complex. The following chemical reactions model this in vivo implementation,

$$\begin{aligned} X_1 &\xrightarrow{w_1} X_1 + Y, \\ X_2 &\xrightarrow{w_2} X_2 + Z, \\ Y + Z &\underset{d}{\overset{a}{\rightleftharpoons}} C, \\ Y &\xrightarrow{\delta} \emptyset, \\ Z, C &\xrightarrow{\phi} \emptyset \end{aligned} \tag{78}$$

Note that the mRNA species ($Y$) and the protein Csy4 ($Z$), as well as the protein-RNA complex ($C$), have different degradation rates to reflect the relative differences in their half-lives. Then, using the law of mass action, we can write down the ODEs that describe the dynamics of these reactions

$$\dot{y} = w_1 x_1 + dc - ayz - \delta y \tag{79}$$
$$\dot{z} = w_2 x_2 + dc - ayz - \phi z \tag{80}$$
$$\dot{c} = ayz - dc - \phi c \tag{81}$$

#### 4.1 Steady-state analysis

First, we notice that when $\dot{c} = 0$ in equation (81),

$$\bar{c} = (\frac{a}{d + \phi})\bar{y}\bar{z} \tag{82}$$

where $\bar{y}$, $\bar{z}$ and $\bar{c}$ are the steady-state values of $y(t), z(t)$ and $c(t)$, respectively. Then, when $\dot{y} = 0$ (that is, from the steady-state of equation (79)), we obtain:

$$w_1 x_1 + d(\frac{a}{d + \phi})\bar{y}\bar{z} - a\bar{y}\bar{z} - \delta\bar{y} = 0 \tag{83}$$

Similarly, when $\dot{z} = 0$ (that is, from the steady-state of Eq. (80)), we obtain

$$w_2 x_2 + d(\frac{a}{d + \phi})\bar{y}\bar{z} - a\bar{y}\bar{z} - \phi\bar{z} = 0 \tag{84}$$

###### 4.1.1 Steady state of $y$

From equation (83) and equation (84),

$$\bar{z} = \frac{w_1 x_1 - \delta \bar{y}}{\bar{y} a(\frac{\phi}{d+\phi})} = \frac{w_2 x_2}{\phi + \bar{y} a(\frac{\phi}{d+\phi})} \tag{85}$$

This leads to a second-order polynomial $P(\bar{y})$ of the shape

$$P(\bar{y}) = \bar{y}^2 + A\bar{y} + B = 0 \tag{86}$$

where $A = \frac{w_2 x_2 - w_1 x_1}{\delta} + \phi \delta K \left[\frac{1}{\phi} + \frac{1}{d}\right]$ and $B = -\phi(w_1 x_1) K \left[\frac{1}{\phi} + \frac{1}{d}\right]$, given the dissociation constant $K = d/a$. This polynomial admits a single real solution,

$$\bar{y} = \frac{1}{2}\left(\frac{w_1 x_1 - w_2 x_2}{\delta} - \phi \delta K \left[\frac{1}{\phi} + \frac{1}{d}\right] + \sqrt{\left(\frac{w_1 x_1 - w_2 x_2}{\delta} - \phi \delta K \left[\frac{1}{\phi} + \frac{1}{d}\right]\right)^2 + 4\phi(w_1 x_1) K \left[\frac{1}{\phi} + \frac{1}{d}\right]}\right) \tag{87}$$

We explore the regime of a small dissociation constant $\xi = K/([x_1 w_1 - x_2 w_2]/\delta) \to 0$. Therefore,

$$\lim_{\xi \to 0} \bar{y} = \frac{1}{2}\left(\frac{w_1 x_1 - w_2 x_2}{\delta} + \sqrt{\left(\frac{w_1 x_1 - w_2 x_2}{\delta}\right)^2}\right) = \frac{1}{2}\left(\frac{w_1 x_1 - w_2 x_2}{\delta} + \left|\frac{w_1 x_1 - w_2 x_2}{\delta}\right|\right) \tag{88}$$

We can rewrite equation (88) as follows:

$$\lim_{\xi \to 0} \bar{y} = \max(0, \frac{w_1 x_1 - w_2 x_2}{\delta}) \tag{89}$$

###### 4.1.2 Steady state of $z$

From Eq. (83) and Eq. (84),

$$\bar{y} = \frac{w_1 x_1}{\delta + \bar{z} a(\frac{\phi}{d+\phi})} = \frac{w_2 x_2 - \phi \bar{z}}{\bar{z} a(\frac{\phi}{d+\phi})} \tag{90}$$

This leads to a second-order polynomial $P(\bar{y})$ of the shape

$$P(\bar{y}) = \bar{y}^2 + A\bar{y} + B = 0 \tag{91}$$

where $A = \frac{w_1 x_1 - w_2 x_2}{\phi} + +\phi \delta K \left[\frac{1}{\phi} + \frac{1}{d}\right]$ and $B = -\delta(w_2 x_2) K \left[\frac{1}{\phi} + \frac{1}{d}\right]$, given the dissociation constant $K = d/a$. This polynomial admits a single real solution,

$$\bar{z} = \frac{1}{2}\left(\frac{w_2 x_2 - w_1 x_1}{\phi} - \phi \delta K \left[\frac{1}{\phi} + \frac{1}{d}\right] + \sqrt{\left(\frac{w_2 x_2 - w_1 x_1}{\phi} - \phi \delta K \left[\frac{1}{\phi} + \frac{1}{d}\right]\right)^2 + 4\delta(w_2 x_2) K \left[\frac{1}{\phi} + \frac{1}{d}\right]}\right) \tag{92}$$

We explore the regime of a small dissociation constant $\xi = K/([x_2 w_2 - x_1 w_1]/\phi) \to 0$. Therefore,

$$\lim_{\xi\to 0}\bar{z} = \frac{1}{2}\left(\frac{w_2x_2 - w_1x_1}{\phi} + \sqrt{\left(\frac{w_2x_2 - w_1x_1}{\phi}\right)^2}\right) = \frac{1}{2}\left(\frac{w_2x_2 - w_1x_1}{\phi} + \left|\frac{w_2x_2 - w_1x_1}{\phi}\right|\right) \quad (93)$$

We can rewrite equation (93) as follows:

$$\lim_{\xi\to 0}\bar{z} = \max(0, \frac{w_2x_2 - w_1x_1}{\phi}) \quad (94)$$

#### 4.2 Temporal analysis

To explore the temporal dynamics of the sequestration reaction, we first define the dissociation constant $K = d/a$ and apply the law of mass conservation: $y^T = y + c$ and $z^T = z + c$. Then, we can rewrite equations (79) to (81) as follows,

$$\begin{aligned}
\frac{d}{dt}y^T &= w_1x_1 - \delta y^T + (\delta - \phi)c && (95)\\
\frac{d}{dt}z^T &= w_2x_2 - \phi z^T && (96)\\
(\frac{K}{d})\frac{d}{dt}c &= (c - y^T)(c - z^T) - Kc - (\frac{K}{d})\phi c && (97)
\end{aligned}$$

Analyzing equation (97) in the regime of a small dissociation constants (or, equivalently, a high sequestration rate), defined by $\xi = K/[(x_1w_1 - x_2w_2)/\delta] \to 0$, we obtain

$$\lim_{\xi\to 0} c(t) = \min(y^T(t), z^T(t)) \quad (98)$$

Furthermore, for convenience, we substract equations (95) and (96), resulting in the following expression,

$$\dot{y}^T - \dot{z}^T = (w_1x_1 - w_2x_2) - \delta y^T + (\delta - \phi)c + \phi z^T \quad (99)$$

Since the analytical solution of equation (99) requires a value of $c(t)$, we can rely on the quasi steady-state approximation described by equation (98), which results in the following cases.

##### 4.2.1 Case N°1: $y^T(t) > z^T(t)$

When $y^T(t) > z^T(t)$, then $c(t) = \min\{y^T(t), z^T(t)\} = z^T(t)$. Therefore, equation (99) simplifies to the following expression,

$$\dot{y}^T - \dot{z}^T = (w_1x_1 - w_2x_2) - \delta(y^T - z^T) \quad (100)$$

Now, if we define $\Delta(t) = y^T(t) - z^T(t)$, so that $F = (\dot{\Delta}) - (w_1x_1 - w_2x_2) + \delta(\Delta) = 0$, we could apply the Laplace transform since $F$ describes a linear ODE. Therefore,

$$\mathcal{L}\{F\} = [\Delta(s)\cdot s - \Delta(0)] - (w_1\frac{x_1}{s} - w_2\frac{x_2}{s}) + \delta\cdot\Delta = 0 \quad (101)$$

where $\Delta(0)$ is defined by the initial conditions. Solving for $\Delta(s)$, we obtain

$$\Delta(s) = \frac{\Delta(0)}{s+\delta} + \frac{w_1x_1 - w_2x_2}{s(s+\delta)} \quad (102)$$

Taking the inverse Laplace transform $\mathcal{L}^{-1}\{\Delta(s)\}$ to recover $\Delta(t)$ yields

$$\Delta(t) = y^T(t) - z^T(t) = \frac{w_1x_1 - w_2x_2}{\delta} + \left(\Delta(0) - \frac{w_1x_1 - w_2x_2}{\delta}\right) e^{-\delta t} \tag{103}$$

##### 4.2.2 Case N°2: $y^T(t) < z^T(t)$

When $y^T(t) < z^T(t)$, then $c(t) = \min\{y^T(t), z^T(t)\} = y^T(t)$. Therefore, equation (99) simplifies to the following expression,

$$\dot{y}^T - \dot{z}^T = (w_1x_1 - w_2x_2) - \phi(y^T - z^T) \tag{104}$$

We take the same definition for $F = F(\Delta)$ as in Case N°1. Then, taking the Laplace transform,

$$\mathcal{L}\{F\} = [\Delta(s) \cdot s - \Delta(0)] - (w_1\frac{x_1}{s} - w_2\frac{x_2}{s}) + \phi \cdot \Delta = 0 \tag{105}$$

where $\Delta(0)$ is defined by the initial conditions. Solving for $\Delta(s)$, we obtain

$$\Delta(s) = \frac{\Delta(0)}{s + \phi} + \frac{w_1x_1 - w_2x_2}{s(s + \phi)} \tag{106}$$

Taking the inverse Laplace transform $\mathcal{L}^{-1}\{\Delta(s)\}$ to recover $\Delta(t)$ yields

$$\Delta(t) = y^T(t) - z^T(t) = \frac{w_1x_1 - w_2x_2}{\phi} + \left(\Delta(0) - \frac{w_1x_1 - w_2x_2}{\phi}\right) e^{-\phi t} \tag{107}$$

##### 4.2.3 Dynamics of $y(t)$

Recalling the equations of mass conservation and since equation (95) does not depend on $\xi$,

$$\lim_{\xi \to 0} y(t) = y^T(t) - \lim_{\xi \to 0} c = y^T(t) - \min(y^T(t), z^T(t)) = \max\{0, y^T(t) - z^T(t)\} \tag{108}$$

Analyzing equation (108) for Case N°2, we notice that $\Delta(t) < 0$ at all times since $y^T(t)$ is assumed to be less that $z^T(t)$. Hence,

$$\lim_{\xi \to 0} y(t) = \max\{0, \Delta(t)\} = 0$$

On the other hand, analyzing equation (108) for Case N°1 results in

$$\lim_{\xi \to 0} y(t) = \max\{0, \Delta(t)\} = \max\left\{0, \frac{w_1x_1 - w_2x_2}{\delta} + \left(\Delta(0) - \frac{w_1x_1 - w_2x_2}{\delta}\right) e^{-\delta t}\right\} \tag{109}$$

Since both the Csy4 endoRNase and the mNeonGreen fluorescent protein are endogenously expressed and aren't native to the HEK293 cell line used for the experiments, we will have that $\Delta(0) = y^T(t)|_{t=0} - z^T(t)|_{t=0} = 0$ $\mu$M. Then, equation (109) simplifies to

$$\lim_{\xi \to 0} y(t) = \max\left\{0, \frac{w_1x_1 - w_2x_2}{\delta}(1 - e^{-\delta t})\right\} \tag{110}$$

Now, let $\lim_{\xi \to 0} y(t) = L$. We observe that

$$\lim_{\delta \to 0} \frac{1 - e^{-\delta t}}{\delta} = t \tag{111}$$

Therefore,

$$\lim_{\delta \to 0} L = \max \{0, (w_1 x_1 - w_2 x_2)t\} \tag{112}$$

Thus, we demonstrate that when the dissociation constant is small and degradation is negligible, the TXTL experimental results emerge as a special case of the more general model described by equations (79) to (81), as equation (112) closely resembles equation (75).

###### 4.2.4 Dynamics of $z(t)$

Recalling the equations of mass conversation and since equation (96) does not depend on $\xi$,

$$\lim_{\xi \to 0} z(t) = z^T(t) - \lim_{\xi \to 0} c = y^T(t) - \min(y^T(t), z^T(t)) = \max\{0, z^T(t) - y^T(t)\} \tag{113}$$

For simplicity, given the definition of $\Delta(t)$ at the beginning of Section 4.2, let's consider $\Delta^*(t) = -\Delta(t) = z^T(t) - y^T(t)$. Then, analyzing equation (113) for Case N°1, we notice that $\Delta^*(t) < 0$ at all times since $z^T(t)$ is assumed to be less than $z^T(t)$ (i.e., the opposite case as for the dynamics of $y(t)$ shown in Section 4.2.3). Hence,

$$\lim_{\xi \to 0} z(t) = \max\{0, \Delta^*(t)\} = 0 \tag{114}$$

On the other hand, analyzing for equation (113) for Case N°2 results in

$$\lim_{\xi \to 0} z(t) = \max\{0, \Delta^*(t)\} = \max \left\{0, \frac{w_2 x_2 - w_1 x_1}{\phi} + \left(\Delta(0) - \frac{w_2 x_2 - w_1 x_1}{\phi}\right) e^{-\phi t}\right\} \tag{115}$$

Under the same assumptions as Section 4.2.3, we can simplify equation (115) as follows

$$\lim_{\xi \to 0} z(t) = \max \left\{0, \frac{w_2 x_2 - w_1 x_1}{\delta}(1 - e^{-\phi t})\right\} \tag{116}$$

Now, let $\lim_{\xi \to 0} z(t) = L$. We observe that

$$\lim_{\phi \to 0} \frac{1 - e^{-\phi t}}{\phi} = t \tag{117}$$

Therefore,

$$\lim_{\phi \to 0} L = \max\{0, (w_2 x_2 - w_1 x_1)t\} \tag{118}$$

Similarly, we demonstrate that when the dissociation constant is small and degradation is negligible, the conclusions drawn from the TXTL model emerge as a special case of the more general model described by equations (79) to (81), as equation (118) closely resembles equation (77).

### 5 Experimental details of DNA hybridization system

Supplementary Table 2 lists the DNA oligos for the molecular classifier using the DNA hybridization system. All the DNA oligos are purchased from IDT. All the sequences are adapted from the genelet system established by Schaffter et al [2]: the sequence of fDNA-nt, fDNA-nt_dummy is adapted from G2C1-nt, the sequence of fDNA-t is adapted from C1-t, and the sequence of qDNA and qDNA_dummy is adapted from dA2. fDNA-nt and fDNA-nt_dummy has the same sequence, but only fDNA-nt has a fluorophore (TYE563) on its 5' end. Similarly, qDNA and qDNA_dummy has the same sequence, but only qDNA has a quencher (Iowa Black RQ) on its 3' end. The annealed DNA complex of fDNA-nt (or fDNA-nt_dummy) and fDNA-t is referred to as fDNA (or fDNA_dummy). fDNA-nt (or fDNA-nt_dummy) and fDNA-t are annealed in 1X NEB RNAPol Reaction Buffer (held at 90°C for 5 minutes then cooled down to 20°C at -1°C/min). Supplementary Figure 1 shows a detailed schematic diagram of the DNA hybridization system. We prepared samples with individually varied fDNA concentration (0 - 400 nM) and qDNA (50 - 400 nM) concentration with steps of 50 nM, using Echo 550 liquid handler (Beckman Coulter). For the liquid handling operation, source plates (xxxxxxx), and destination plates (3357, Corning Inc.) were used. The total volume of each sample is 2 $\mu$L, which include fDNA and qDNA at specified concentrations, and 1X NEB RNAPol Reaction Buffer. After preparing samples on the desination plate, incubate the plate at 37°C for 1 hour (except for E1-004 in Table 5). After the incubation, the destination plate is loaded into a mcroplate reader (Biotek, Synergy H1) to obtain fluorescence signal at excitation wavelength 539 nm and emission wavelength 569 nm at 37°C.

***Table 1:*** *A list of the DNA sequences for the DNA hybridization system*

| Sequence ID | DNA Sequence (5' to 3') |
|---|---|
| fDNA-nt | /5TYE563/AGCCAAGATTCAGGTCAATAAGTGACCAAGTAATACGACTCACTATAGGGAGATTCGTCTCCCAAATCCTTCCATGAACGCCAAACCGTG |
| fDNA-nt_dummy | AGCCAAGATTCAGGTCAATAAGTGACCAAGTAATACGACTCACTATAGGGAGATTCGTCTCCCAAATCCTTCCATGAACGCCAAACCGTG |
| fDNA-t | GAGTAGGTCGTCGCCACGGTTTGGCGTTCATGGAAGGATTTGGGAGACGAATCTCCCTATAGTGAGTCG |
| qDNA | CATCCCACTATTACTTGGTCACTTATTGACCTGAA/3IABRQSp/ |
| qDNA_dummy | CATCCCACTATTACTTGGTCACTTATTGACCTGAA |

Table 5 is a list of the experiments using DNA hybridization systems, with detailed variable assignments. Figure 2 shows all the replicate data for each experiment. E1-002 and E1-003 have two replicates, while the rest of the experiments have three replicates. The rightmost column of Figure2 shows the averaged data for each experiment. Unless otherwise noted, the averaged data are included in the figures in the main and the supplementary text. E1-004 was a time-course measurement, while the other experiment have only one time point after one-hour incubation at 37°C. Figure **??** shows the time-course data of E1-004 and heatmap snapshots at 0 and 60 minutes. Because the initial measurement is performed just after a sample is loaded into the microplate reader, it is assumed that the initial measurement corresponds to the fluorescence data at the room temperature, while the last measurement at 60 minutes is assumed to correspond to the measurement at 37°C.

***Table 2:*** *A list of the experiments using the DNA hybridization system*

| Experiment ID | Experimental details |
|---|---|

| | |
|---|---|
| E1-002 | X1: 25% fDNA, X2: 100% qDNA |
| E1-003 | X1: 100% fDNA, X2: 25% qDNA |
| E1-004 | X1: 100% fDNA, X2: 100% qDNA |
| E1-010 | X1: 100% fDNA, X2: 100% fDNA, X0: 300 nM qDNA |
| E1-011 | X1: 100% fDNA, X2: 100% fDNA, X0: 500 nM qDNA |
| E1-012 | X1: 100% qDNA, X2: 100% qDNA, X0: 300 nM fDNA |
| E1-013 | X1: 100% qDNA, X2: 100% qDNA, X0: 500 nM fDNA |

### 6 Weight tuning by the addition of dummy DNA strands in the DNA hybridization system

We evaluate additional patterns of decision boundary by changing the assignment of DNA species to input and bias (Figure 3). In Figure 3A, qDNA without a quencher, which is called qDNA_dummy, is added. The fraction of qDNA_dummy is 25%. The addition of qDNA_dummy dilutes the effective fraction of qDNA with a quencher, which is expect to increases the slope of the decision boundary. In Figure 3B, fDNA without a fluorophore, which is called fDNA_dummy, is added. The fraction of the dummy strand is 25% for both cases. The addition of the dummy strands dilutes the effective fraction of qDNA or fDNA, which is expected to increases or decreases the slope of the decision boundary respectively, and the experimental results showed this trend. The numerical simulations fit the experimental results better with lower affinity between a regular strand and a dummy strand compared with the regular hybridization with both a fluorophore and a quencher; i.e., the association rate constant for dummy qDNA $a_d = 30$, and the dissociation rate constant for qDNA $d_d = 30$, while the regular association rate constant $a = 100$, and dissociation rate constant $d = 3$. These parameter adjustments suggest that the incorporation of the fluorophore/quencher pair stabilizes the DNA duplex, which is consistent with a prior study that reported the stabilizing effect of particular fluorophore/quencher pairs [3].

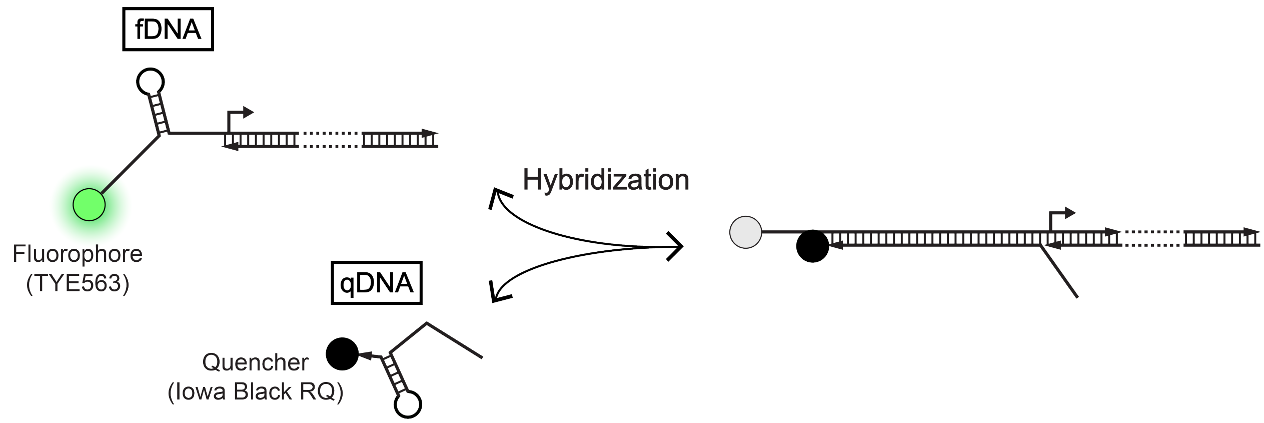

***Figure 1:*** *The schematic diagram of fDNA, qDNA, and fDNA-qDNA complex in the DNA hybridization system. fDNA has a fluorophore (TYE563), and qDNA has a quencher (Iowa Black RQ).*

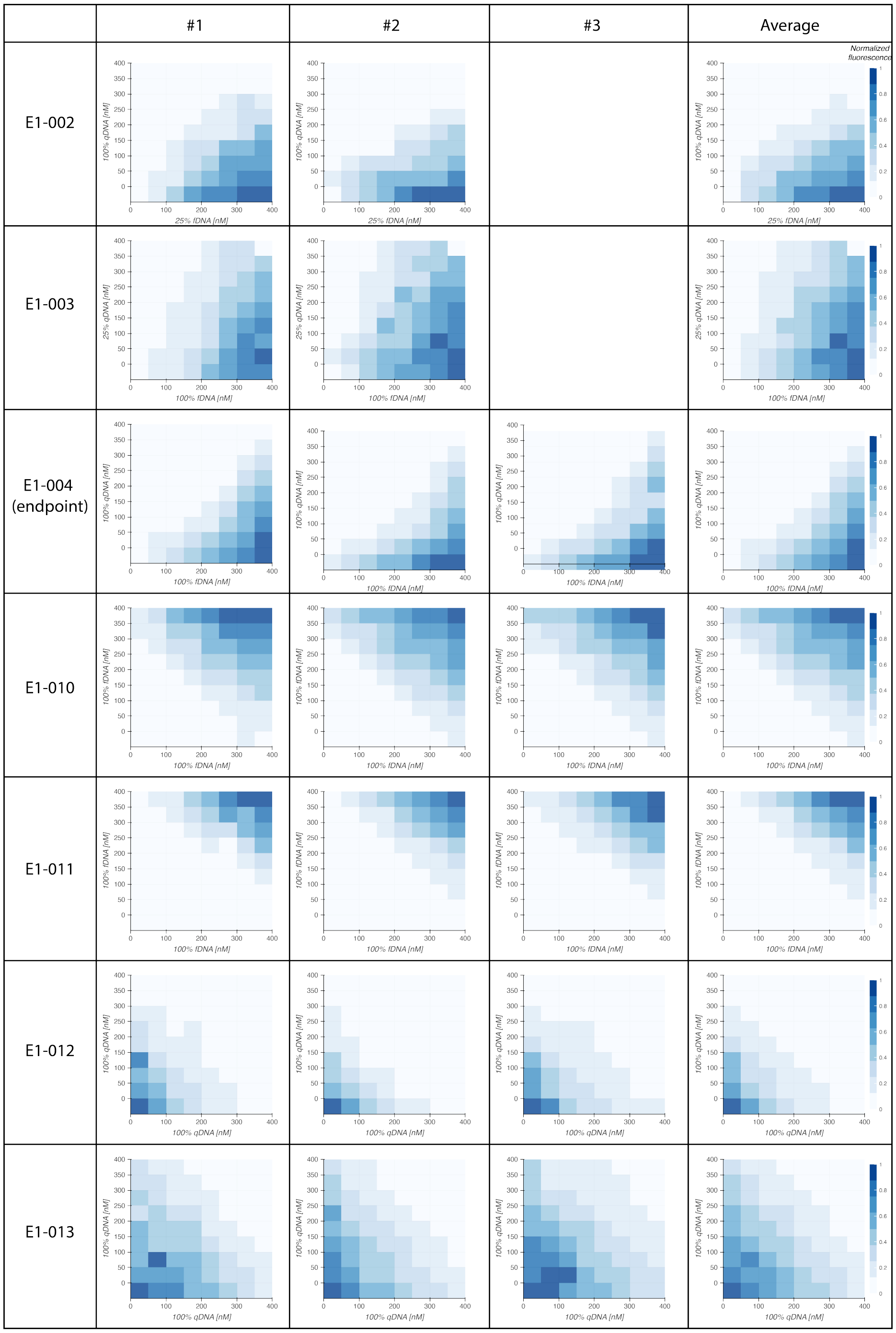

***Figure 2:*** *Replicates of all the experiments presented in the main text and the supplementary text. The leftmost cell in each row indicates the experiment ID for each experiment, which corresponds to the ID listed in Table 5. The E1-002, and E1-003 have two replicates, while the other experiments have three replicates. The rightmost column indicates the averaged data for each experiment.*

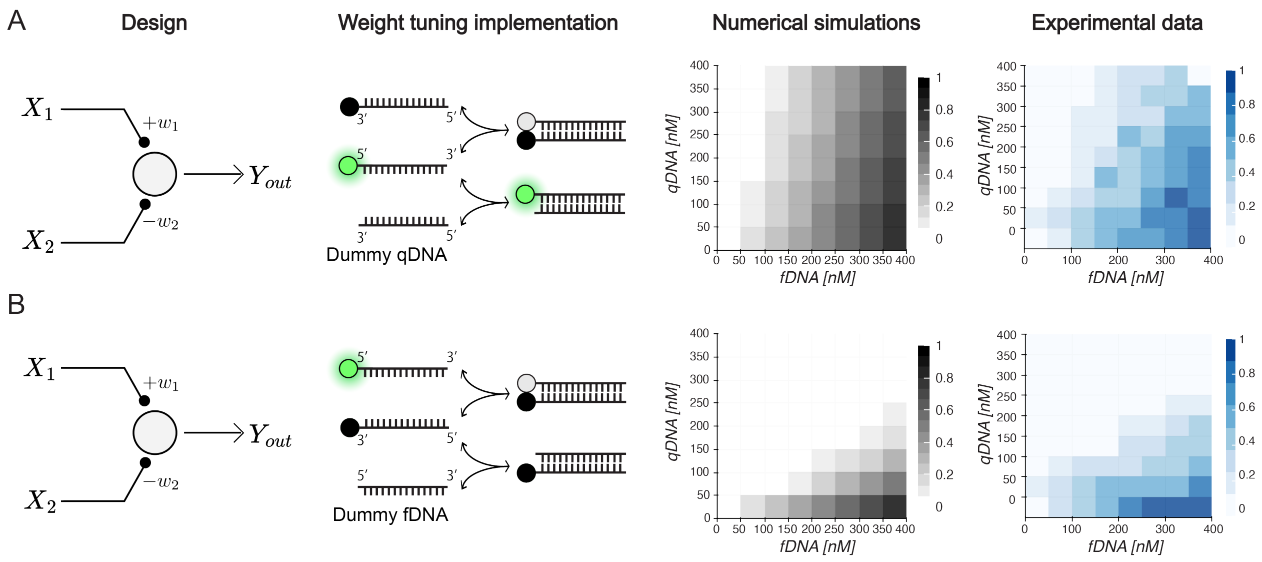

***Figure 3:*** *Decision boundaries with dummy DNA strands. From left to right, diagrams of the classifiers, hybridization reaction diagrams, numerical simulations, and experimental results of DNA hybridization system with (A) dummy qDNA strand and (B) dummy fDNA strand. Kinetic parameter values for the numerical simulations are as follows.* $a = 100$, $a_d = 30$, $d = 1$, $d_d = 3$.

### 7 Experimental details of TXTL system

#### 7.1 Activation function

Equation (75) predicts a time-dependent Rectified Linear Unit (ReLU)-like activation function, assuming a sufficiently small dissociation constant. In our case study of the $\sigma_w$-RsiW protein-based molecular perceptron, we analyze how the concentrations of $\sigma_w$ ($x_1$) and RsiW ($x_2$) — the inputs — map to the fluorescent readout — the output — over time. The expected behavior is absent fluorescence whenever $w_1x_1 < w_2x_2$ and fluorescence whenever $w_1x_1 > w_2x_2$, where $w_1$ and $w_2$ are the weights assigned to each input. This corresponds to the decision boundary observed in the heatmap shown in Figure 4-A.

Fitting the data captured at $t = 6$ h from the main text (Main Figure 3) to the mathematical model described by equation (75) reveals that $w_1 = w_2 = 1$ (/h). As a result, the activation function simplifies: fluorescence remains zero when RsiW is more abundant than $\sigma_w$ ($x_1 < x_2$), and turns on otherwise ($x_1 > x_2$), with intensity proportional to the difference $x_1 - x_2$. This behavior is illustrated in Figure 4-B. For better visualization, normalizing the x-axis ($\sigma_w$) relative to RsiW further highlights the fluorescence response, which closely resembles the ReLU activation function (Figure 4-C).

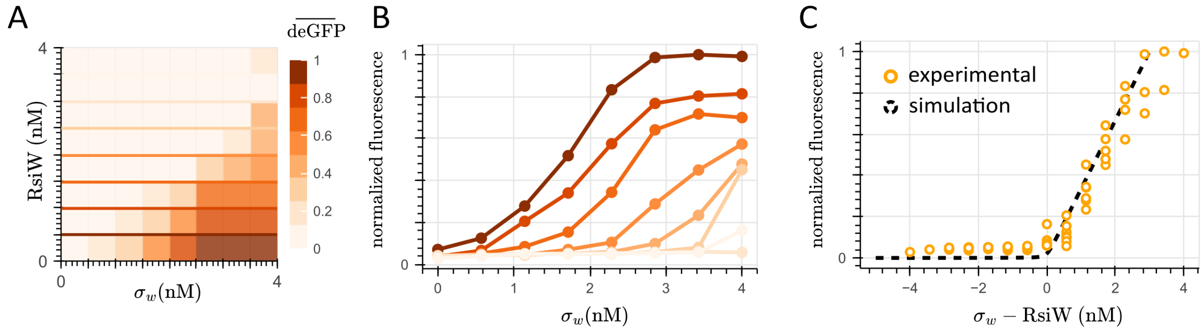

*Figure 4: **Input-output map of the molecular sequestration reaction between $\sigma_w$ and RsiW proteins**. Panel A shows the heatmap constructed from snapshots at $t = 6$ h from the time-course measurement shown in Figure 5. The fluorescence signals were normalized using the maximum fluorescence value obtained for $(\sigma_w, \text{RsiW}) = (4, 0)$ nM. Horizontal lines are shown to represent the 8 bins from which Panel B is constructed, showing the input ($\sigma_w$)-output (fluorescence readout) of the TXTL reaction. Panel C normalizes the horizontal axis by substracting the RsiW concentration with respect to the $\sigma_w$ concentration, and displays this difference with respect to the fluorescence readout. The numerical simulation is shown as black dashed lines, obtained by numerically integrating equations (62) to (64) using the following nominal parameters: $a = 100$ (/h/$\mu$M), $d = 1$ (/h), and $w_1 = w_2 = 1$ (/h).*

#### 7.2 Temporal pattern recognition

We conducted kinetic experiments to validate the time-dependency of the decision boundary predicted from equation (75). Figure 5 show the time-course fluorescent readout for a 8×8 grid of $\sigma_w$/RsiW proteins arranged in a well-plate, starting with no plasmid at the bottom-left and increasing up to 4 nM (horizontal: $\sigma_w$, vertical: RsiW) in increments of 0.57 nM. During the 6-hour measurement period, the system does not reach steady-state. However, a well-defined decision boundary emerges as early as $t = 3$ hours. By normalizing the fluorescence readout at each hour to the maximum measurement at $t = 6$ hours, we constructed heatmaps in the $x_1$-$x_2$ plane (i.e., $\sigma_w$-RsiW in this example) and confirmed the presence of the decision boundary at $t = 3$ hours (Fig. 7–A and B). Replicates of the heatmap showing the linear decision boundary are shown in Figure 8–A and B, and were constructed from endpoint fluorescence measurements after 3, 4 and 5 hours of incubation at $30°$C.

Likewise, Figure 6 shows the time-course fluorescent readout for a 6×6 grid of $\sigma_{28}$/FlgM proteins arranged similarly to the previous implementation, but with a maximum plasmid concentration of 1 nM and increments of 0.2 nM. For this example, the fluorescence signal plateaus around $t = 5$ hours; however, despite this plateau, the system does not reach a steady state, but all available resources have been consumed. The decision boundary is however observed as early as $t = 3$ hours, as well. This statement is further confirmed by the presence of the decision boundary in the heatmap at $t = 3$ hours of Fig. 7–C and D, as well. Replicates of the heatmap showing the linear decision boundary are shown in Figure 8–C and D. The heatmaps in Panel C were constructed using snapshots from a time-course measurement at 3, 4 and 5 hours, while the heatmaps in Panel D were constructed from endpoint fluorescence measurements after 3, 4 and 5 hours of incubation at $30°$C.

#### 7.3 Weight tuning

We then evaluate if we can tune the slope of the decision boundary by adjusting the inputs. As shown in the numerical simulations of the main text (Main Figure 3), we can increase the slope of the decision boundary by either increasing the weight associated to the $X_1$ species (e.g., the $\sigma$ factor) or decreasing the weight associated to the $X_2$ species (e.g., the anti-$\sigma$ factor). In Figure 9, we used a dummy plasmid (i.e., a plasmid that expresses a protein not involved in the sequestration reaction) to dilute the total effective concentration of the RsiW protein (anti-$\sigma_w$), which is expected to increase the slope of the decision boundary. Similar to Figure 5, the system does not reach steady-state in the 6-hour measurement time, but a decision boundary with an increased slope can be observed since $t = 3$ hours. This observation is further validated in the heatmaps shown in Figure 11–A and B. Replicates of this behavior can be found in Figure 12–A and B, in which the heatmaps were constructed from two endpoint fluorescence measurements at 3 and 5 hours of incubation time at $30°$C.

Following a similar logic, in Figure 10, we used another dummy plasmid to dilute the total concentration of FlgM protein (anti-$\sigma_{28}$), which is expect to decrease the slope of the decision boundary. Similar to Figure 6, the system does reach steady-state in the 6-hour measurement time, but the decision boundary with the decreased slope can be observed as early as $t = 3$ hours. This observation is further validated in the heatmap snapshots shown in Figure 11–C and D. Replicates of this behavior can be found in Figure 12–C and D. The heatmap in Panel C was constructed from endpoint fluorescence measurements after 3, 4 and 5 hours of incubation at $30°$C, while Panel D was constructed using snapshots from a time-course measurement at 3, 4 and 5 hours.

#### 7.4 Flipping the decision boundary and inverting the classification region

From equation (75), we notice that if we invert the inputs $x_1$ and $x_2$, we will obtain the same decision boundary, but an inverted classification region, so that the output of the molecular perceptron will be proportional to the difference between $w_2x_2 - w_1x_1$, whenever that difference is positive. We then evaluate other patterns of decision boundary by considering another constant input to the molecular sequestration reaction, also named "bias". If we intend to parametrize the decision boundary for when the weights are equal ($w_1 = w_2$), we could roughly approximate it by a linear equation $y = x$. Then, to flip the decision boundary (so that $y = -x$ is what we observe), we added a constant plasmid concentration of $\sigma_w = 1$ nM to the TXTL master mix, which in the main text (Main Figure 4) is represented by the $X_0$ species and assigned to a positive weight $w_0$. The remaining species ($X_1$ and

$X_2$, encoded in two plasmids that express the RsiW protein) are assined negative weights. The resulting heatmaps can be observed in Figure 13–A.

Furthermore, to invert the classification region (i.e., cover the uppermost region of the $x_1 - x_2$ plane), we added a constant plasmid concentration of FlgM = 1 nM to the TXTL master mix, which is also represented by the $X_0$ species in the main text (Main Figure 4), and is associated with a negative weight $w_0$. The remaining species are encoded in two plasmids that express the $\sigma_{28}$ factor, and whose expression is associated with positive weights. The resulting heatmaps can be observed in Figure 13–B. All heatmaps were constructed using endpoint fluorescent measurements after 3, 4 and 5 hours of incubation at $30°$C.

#### 7.5 Troubleshooting

The unexpected patterns observed in the heatmaps of the middle column of Figure 14, which deviate from the expected behavior, can be explained by adjusting the weights associated with the production of the $X_1$ and $X_2$ species, as well as the initial concentration of the $X_0$ species. These adjustments suggest possible dilution errors and/or inaccuracies in the preparation of the TXTL master mix. We identified three representative cases where the mathematical model, described by equations (62) to (64), provided insights into these inconsistencies.

The first case is a three-input implementation using the $\sigma_{28}$ and FlgM proteins, designed to invert the classification region. As shown in Figure 14–A, this case suggests an incorrect plasmid concentration in the TXTL master mix. The classification region aligns with the biochemical implementation; however, the expanded region with minimal to no fluorescence indicates an excess of FlgM plasmid (anti-$\sigma_{w8}$), which sequesters $\sigma_{28}$ and prevents fluorescence gene expression. Additionally, the increased slope of the decision boundary suggests an incorrect dilution of the plasmid encoding $\sigma_{28}$, particularly the one associated with the $w_2$ weight, whose decrease leads to a higher decision boundary. Overall, this experimental outcome can be explained by the following parameter values: $w_1 = 0.6$ (/h), $w_2 = 0.35$ (/h), $w_0 = 1$ (/h), and an increased constant RsiW concentration in the TXTL master mix of approximately 2 nM.

The second case follows the same design as before. Similarly, as shown in Figure 14–B, this case suggests an incorrect plasmid concentration in the TXTL master mix, as the region with minimal to no fluorescence is also extended, indicating an incorrect dilution of the plasmid encoding $\sigma_{28}$. However, unlike the previous case, we observe a decrease in the decision boundary, which suggests an incorrect dilution of the plasmid associated with the $w_1$ weight. Overall, this experimental outcome can be explained by the following parameter values: $w_1 = 0.3$ (/h), $w_2 = 0.6$ (/h), $w_0 = 1$ (/h), and an increased constant FlgM concentration in the TXTL master mix of approximately 2.5 nM.

The third and final case involves a three-input implementation using $\sigma_w$ and RsiW proteins, designed to flip the decision boundary. In this experimental setup, the fluorescent output is highly dependent on the concentration of $\sigma_w$ added to the TXTL master mix, as it enables the expression of the fluorescent gene. Consequently, the offset observed in Figure 14–C toward the upper-right corner can be attributed to an increased concentration of $\sigma_w$ in the master mix. By slightly adjusting the weights in the expected simulation for a better fit to the experimental data—specifically, $w_1 = 0.25$ (/h), $w_2 = 0.2$ (/h), and $w_0 = 1$ (/h)—the experimental outcome can be explained by an increased constant $\sigma_w$ concentration in the TXTL master mix of approximately 1.6 nM.

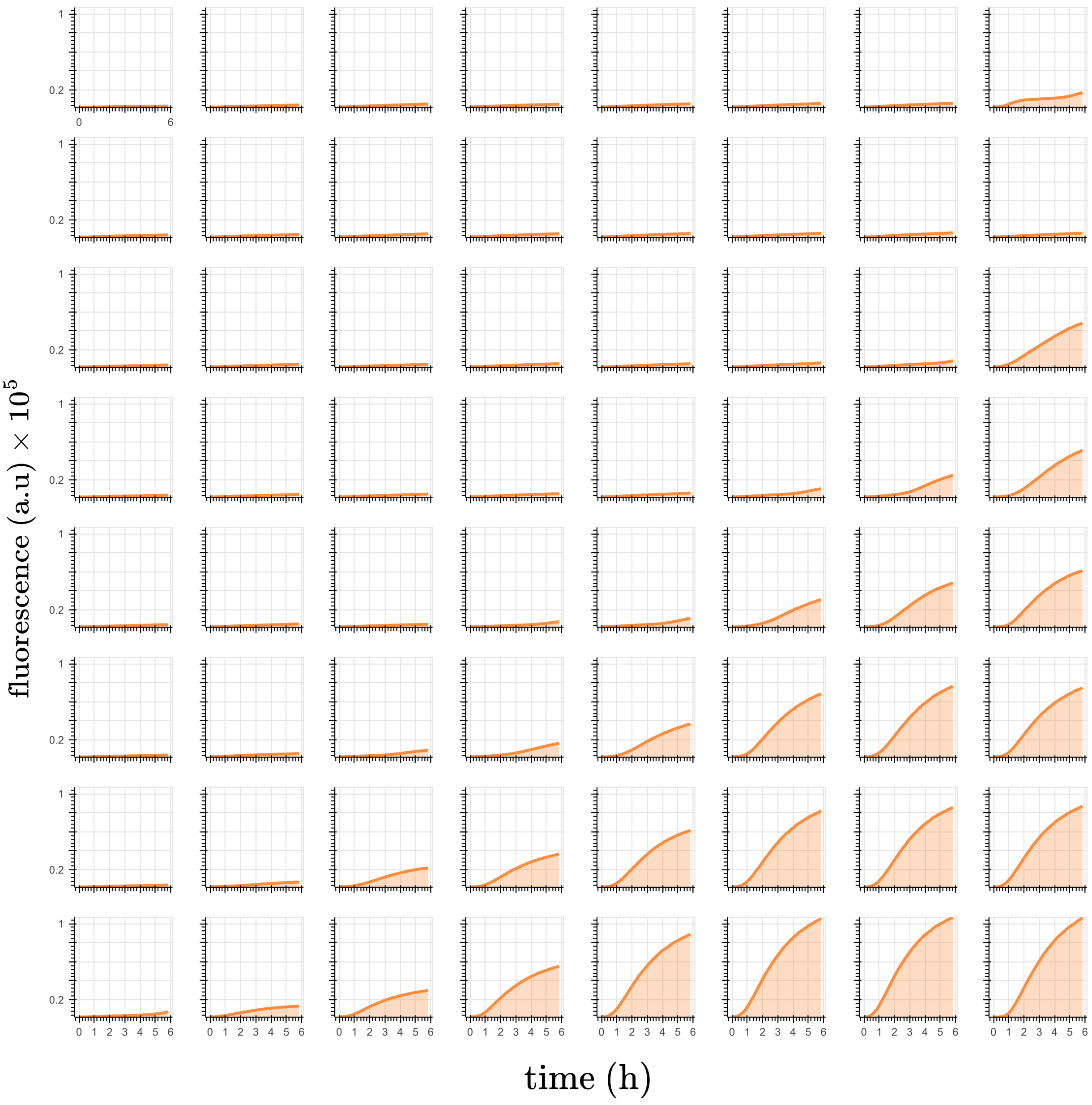

***Figure 5: Time-course measurement of the molecular sequestration reaction between $\sigma_w$ and RsiW proteins***. *The concentration of $\sigma_w$ increases from 0 nM on the left to 4 nM on the right, while the concentration of RsiW increases from 0 nM at the bottom to 4 nM at the top, both in increments of 0.57 nM.*

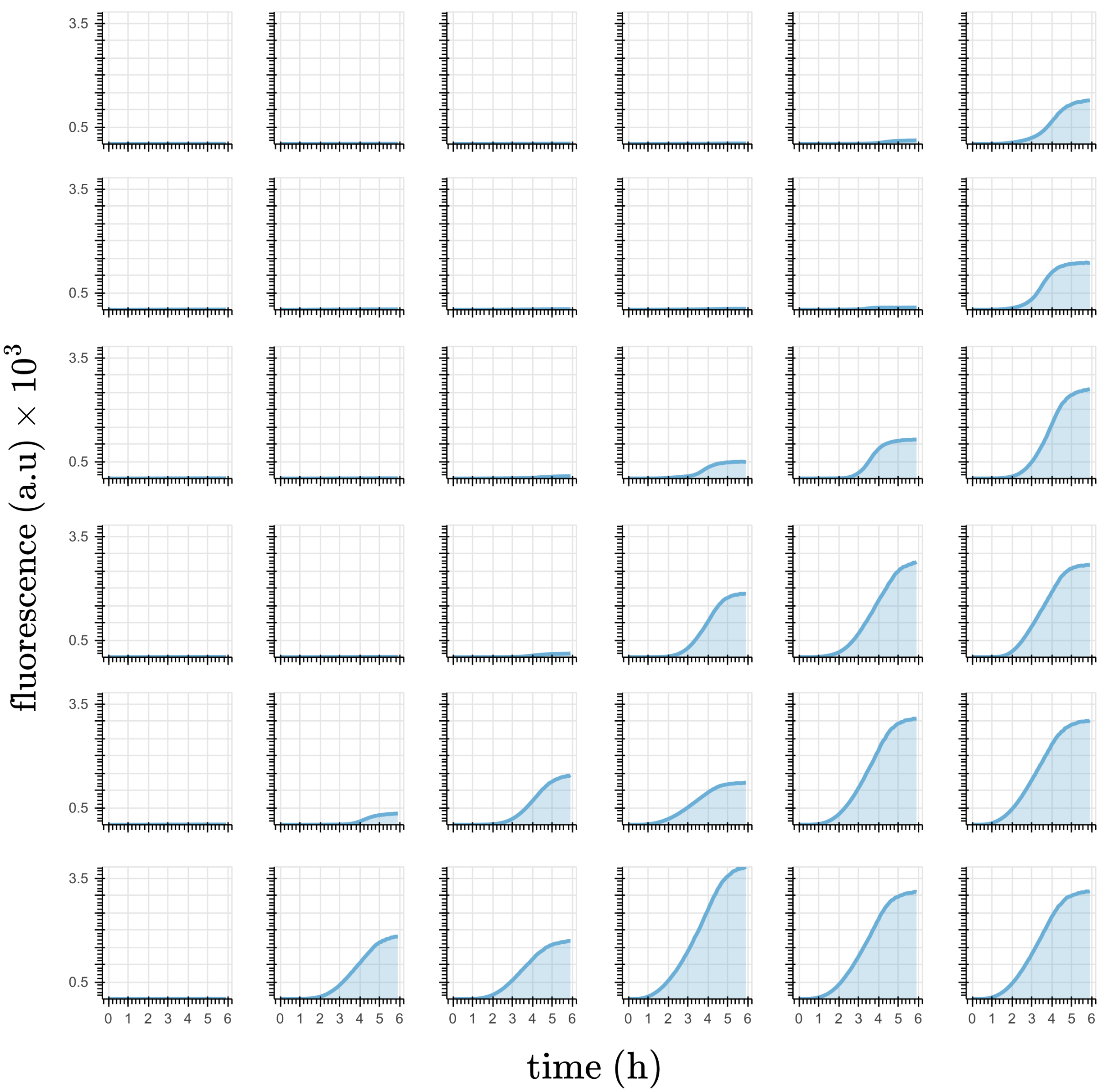

***Figure 6: Time-course measurement of the molecular sequestration reaction between $\sigma_{28}$ and FlgM***
*The concentration of $\sigma_{28}$ increases from 0 nM on the left to 1 nM on the right, while the concentration of FlgM protein increases from 0 nM at the bottom to 1 nM at the top, both in increments of 0.2 nM.*

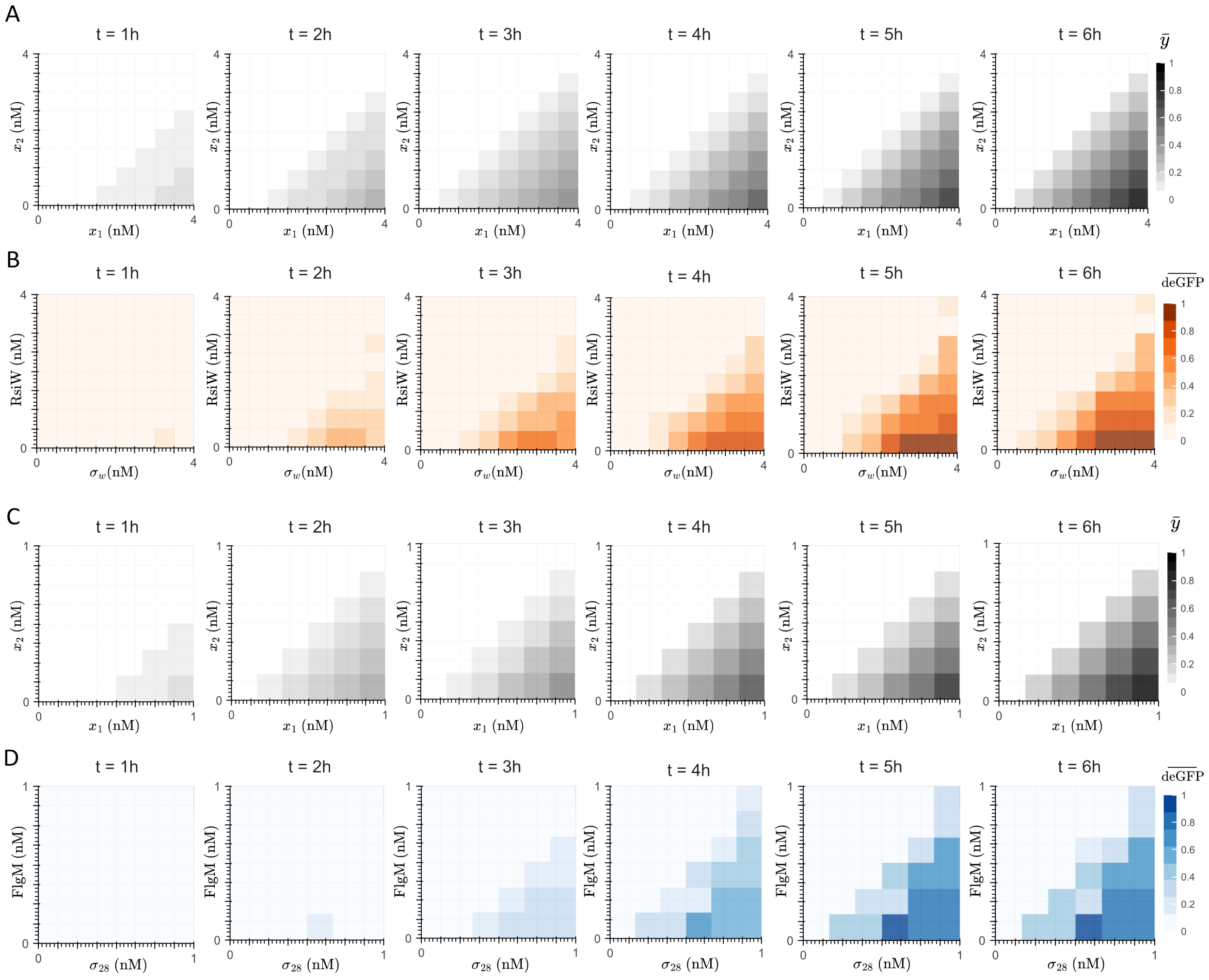

***Figure 7: Temporal pattern recognition.*** *For the biochemical implementation based on* $\sigma_w$ *and RsiW proteins, Panel A shows the heatmaps at 6 time points obtained by numerically solving equations* (62) *to* (64) *with an 8x8 resolution, while Panel B shows the experimental results. For the biochemical implementation based on* $\sigma_{28}$ *and FlgM proteins, Panel C shows the heatmaps obtained by solving the aforementioned equations with an 6x6 resolution, while Panel D shows the experimental results. For both implementations, the fluorescence signals in the heatmaps were normalized using the maximum fluorescence value recorded at the 6-hour time point. Kinetic parameter values for the numerical simulations are as follows:* $a = 100$ *(/h/*$\mu M$*)* , $d = 1$ *(/h),* $w_1 = w_2 = 1$ *(/h)*

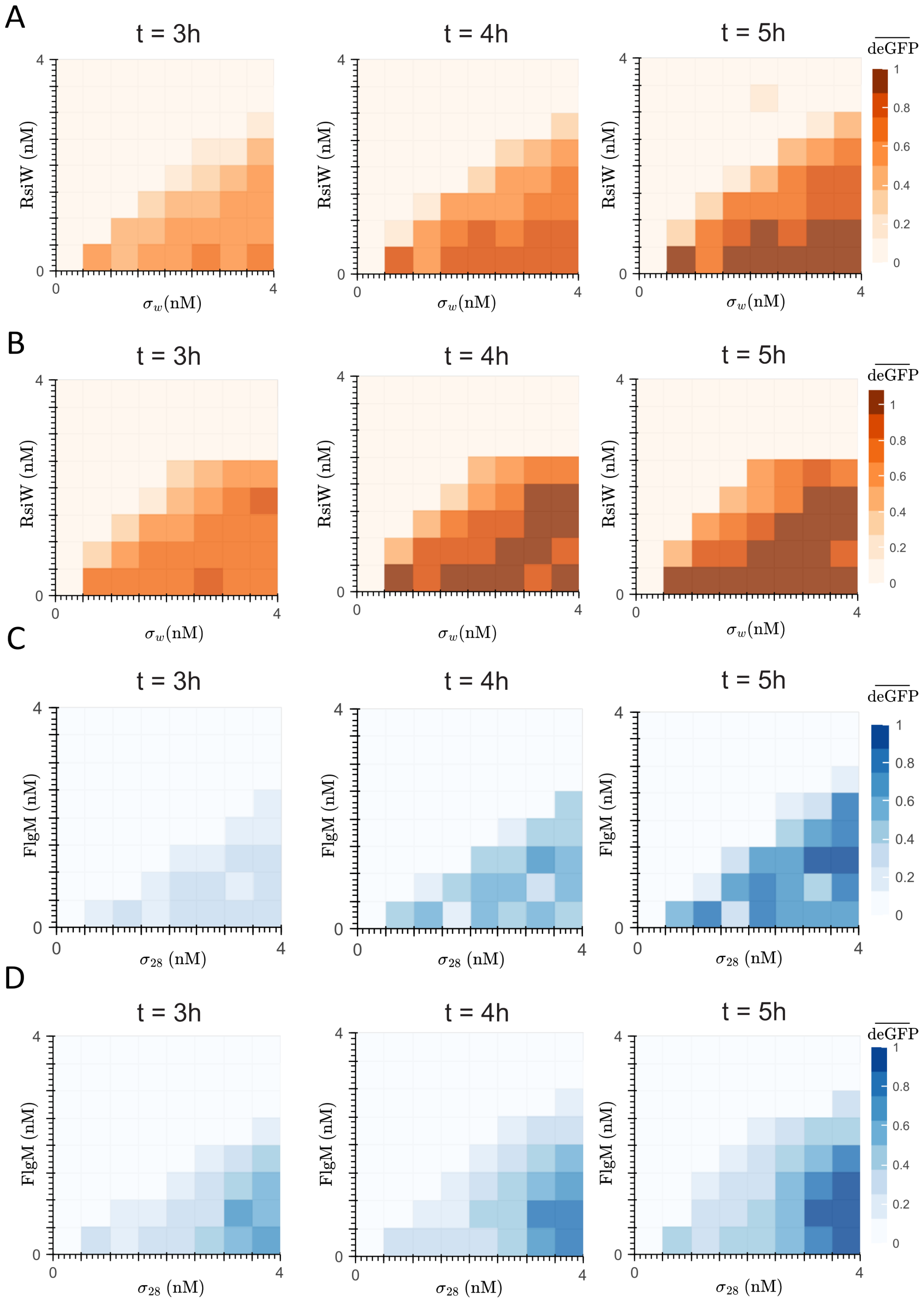

***Figure 8: Temporal pattern recognition - Replicates*** *Panel A and B show the heatmaps for the biochemical implementation based on* $\sigma_w$ *and RsiW factors. Panel C and D show the biochemical implementation based on* $\sigma_{28}$ *and FlgM factors. For both implementations, the fluorescence signals in the heatmap were normalized using the maximum fluorescence value recorded at the 5-hour time point.*

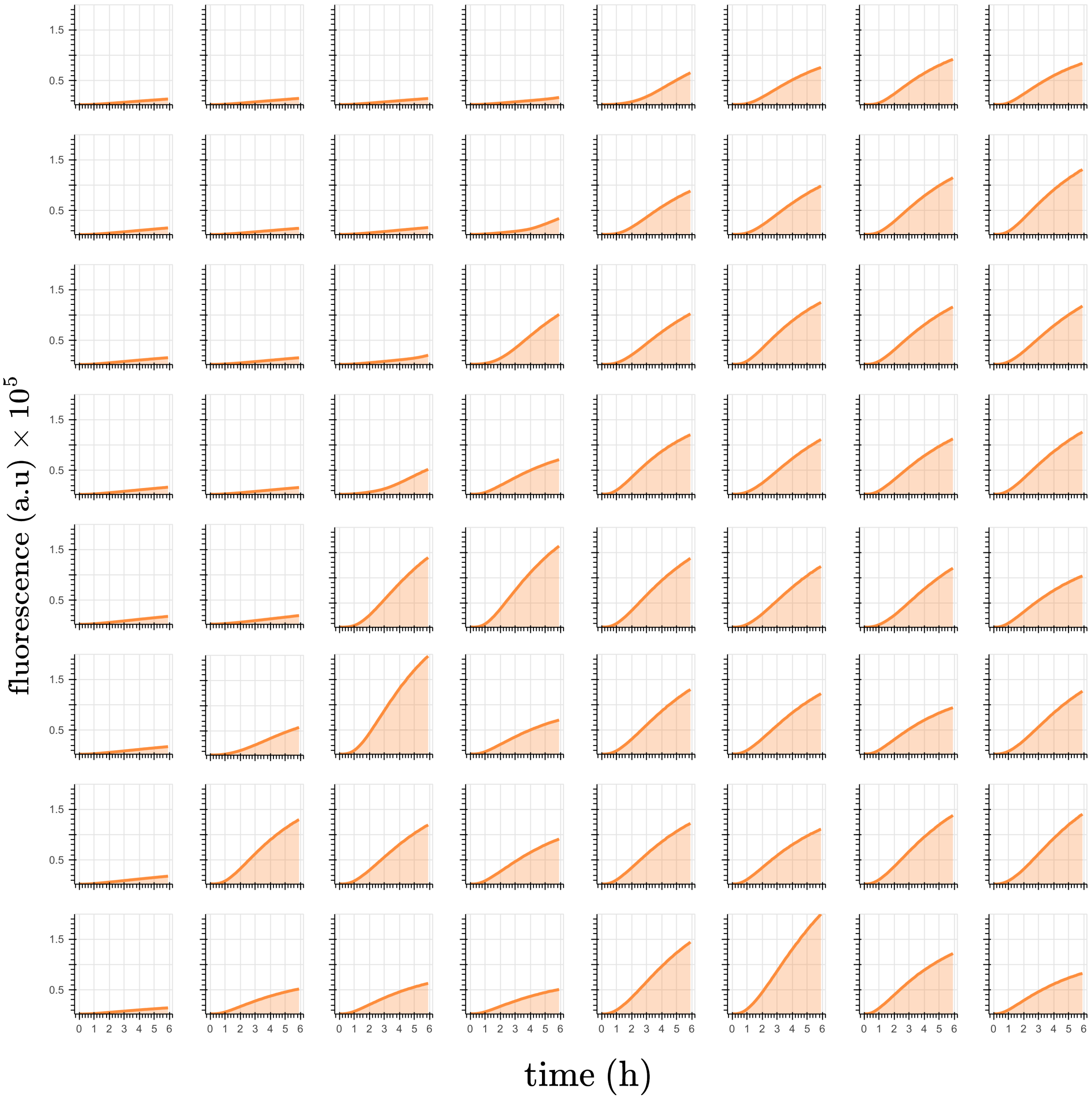

***Figure 9: Time-course measurements of the molecular sequestration reaction between $\sigma_w$ and anti-$\sigma_w$ pair showing an increase in the slope of the decision boundary****. The concentration of $\sigma_w$ increases from 0 nM on the left to 4 nM on the right, while the concentration of RsiW increases from 0 nM at the bottom to 2 nM at the top. To maintain a constant plasmid concentration of 4 nm, a dummy plasmid is added to the RsiW mix accordingly.*

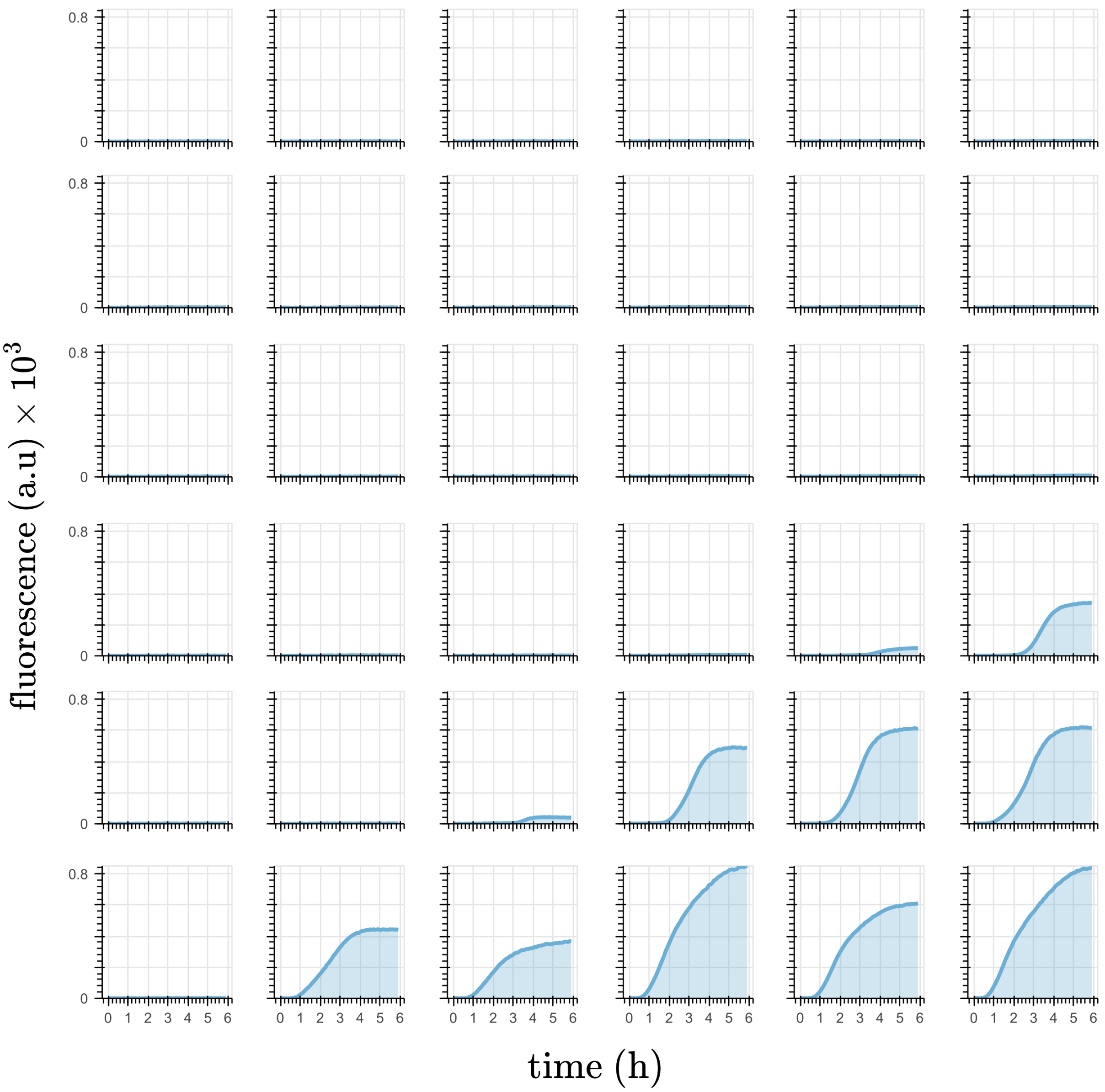

***Figure 10: Time-course measurements of the molecular sequestration reaction between $\sigma_{28}$ and FglM pair showing a decrease in the slope of the decision boundary**. The concentration of $\sigma_{28}$ increases from 0 nM on the left to 0.57 nM on the right, while the concentration of FlgM increases from 0 nM at the bottom to 0.6 nM at the top. To maintan a total plasmid concentration of 0.6 nM, a dummy plasmid is added to the $\sigma_{28}$ mix accordingly.*

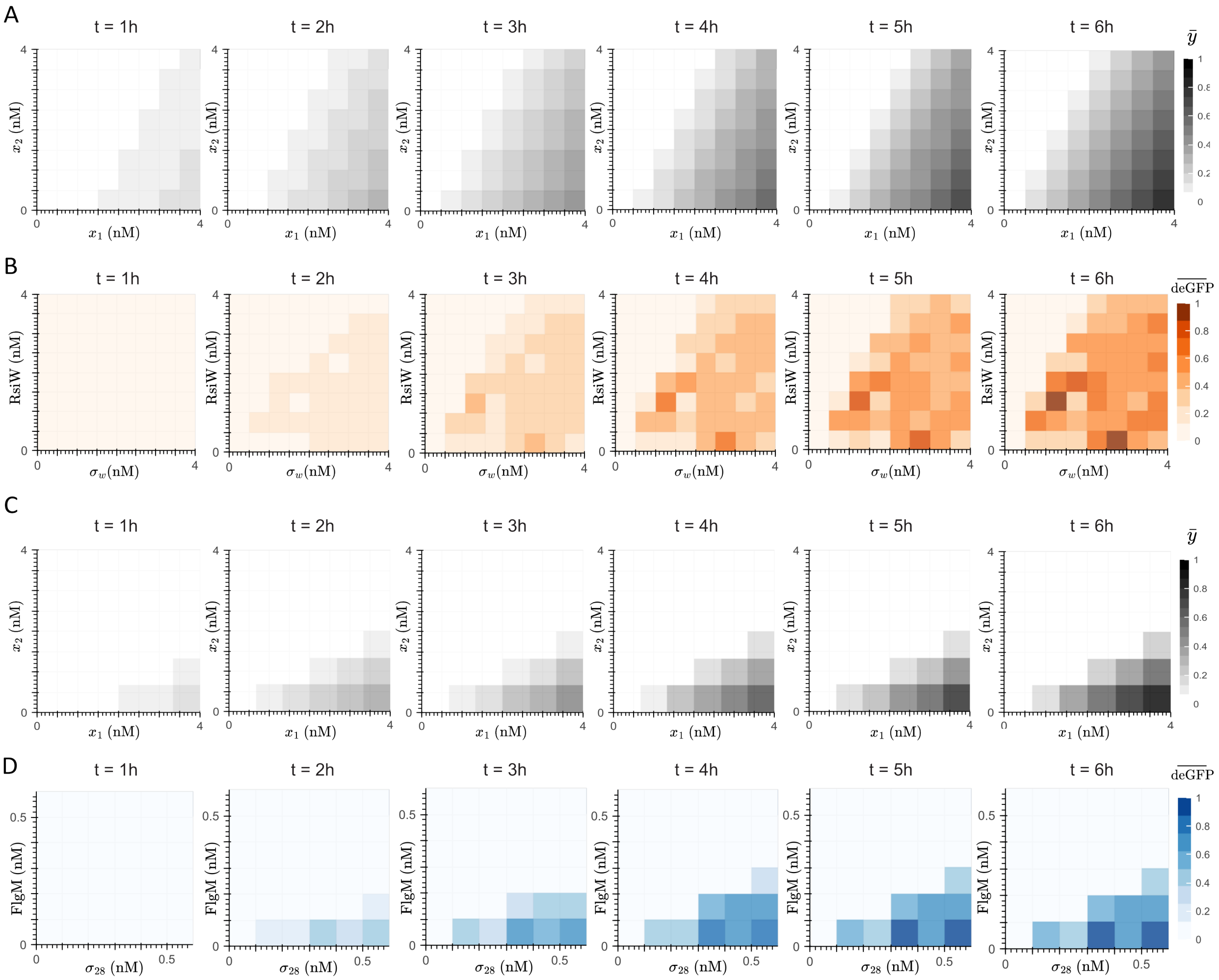

***Figure 11: Temporal pattern recognition with adjusted weights.*** *For the biochemical implementation based on $\sigma_w$ and RsiW proteins, Panel A shows the heatmaps at 6 time points obtained by numerically solving equations* (62) *to* (64) *with an 8x8 resolution, considering $w_1 = 1$ (/h) and $w_2 = 0.5$ (/h). Panel B shows the corresponding experimental results. For the biochemical implementation based on $\sigma_{28}$ and FlgM proteins, Panel C shows the heatmaps obtained by solving the aforementioned equations with an 6x6 resolution, considering $w_1 = 0.5$ (/h) and $w_2 = 1$ (/h). Panel D shows the corresponding experimental results. For both biochemical implementations, the fluorescence signals in the heatmaps were normalized using the maximum fluorescence value recorded at the 6-hour time point. Kinetic parameter values for the numerical simulations are as follows: $a = 100$ (/h/$\mu$M) and $d = 1$ (/h).*

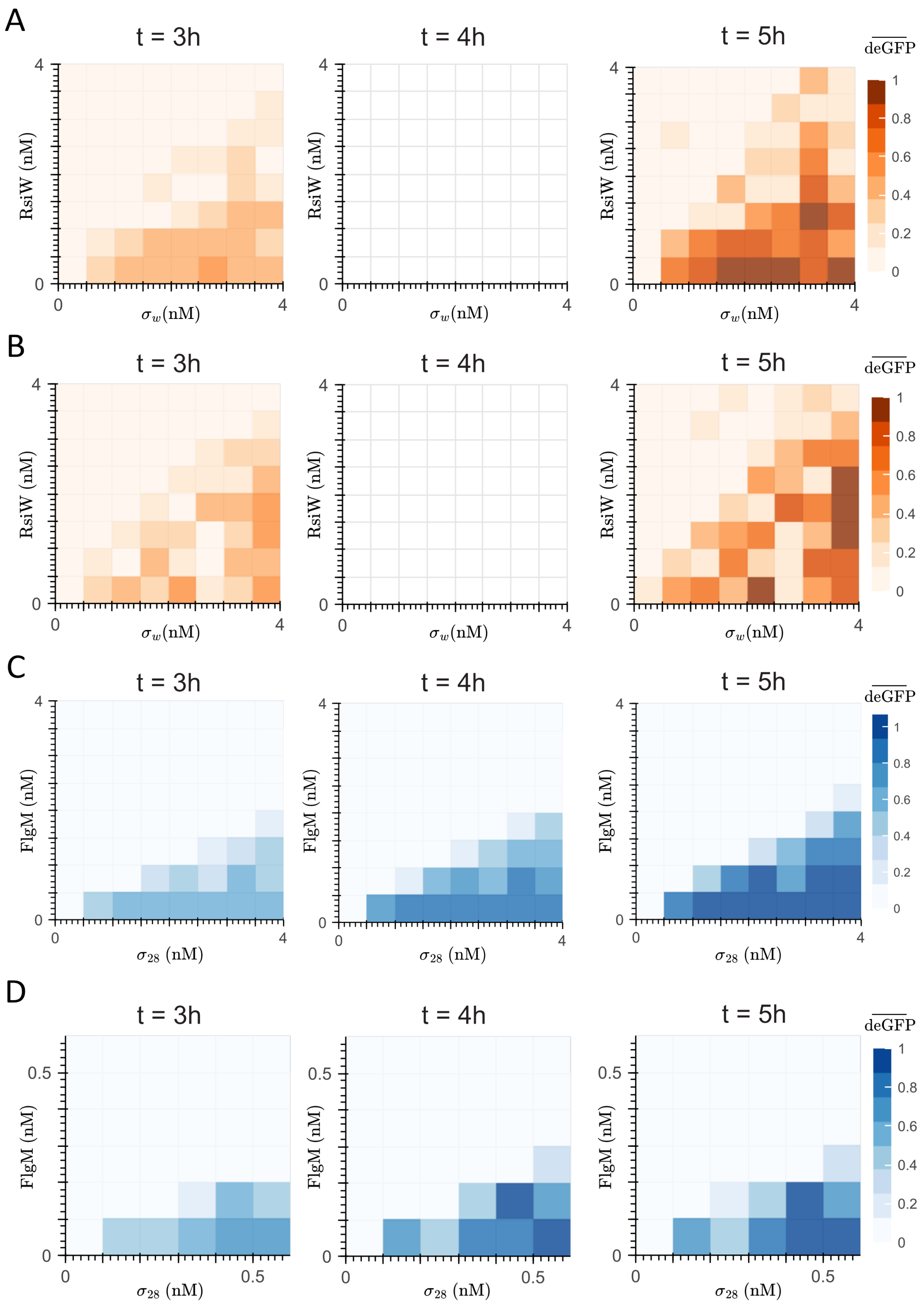

***Figure 12: Weight tuning of the slope defined by the decision boundary - Replicates***. *Panel A and B show the heatmaps for the biochemical implementation based on $\sigma_w$ and RsiW proteins. There was no measurement for $t = 4$ hours. Panel C and D show the biochemical implementation based on $\sigma_{28}$ and FlgM proteins. For every implementation, the fluorescence signals in the heatmap were normalized using the maximum fluorescence value recorded at the 5-hour time point.*

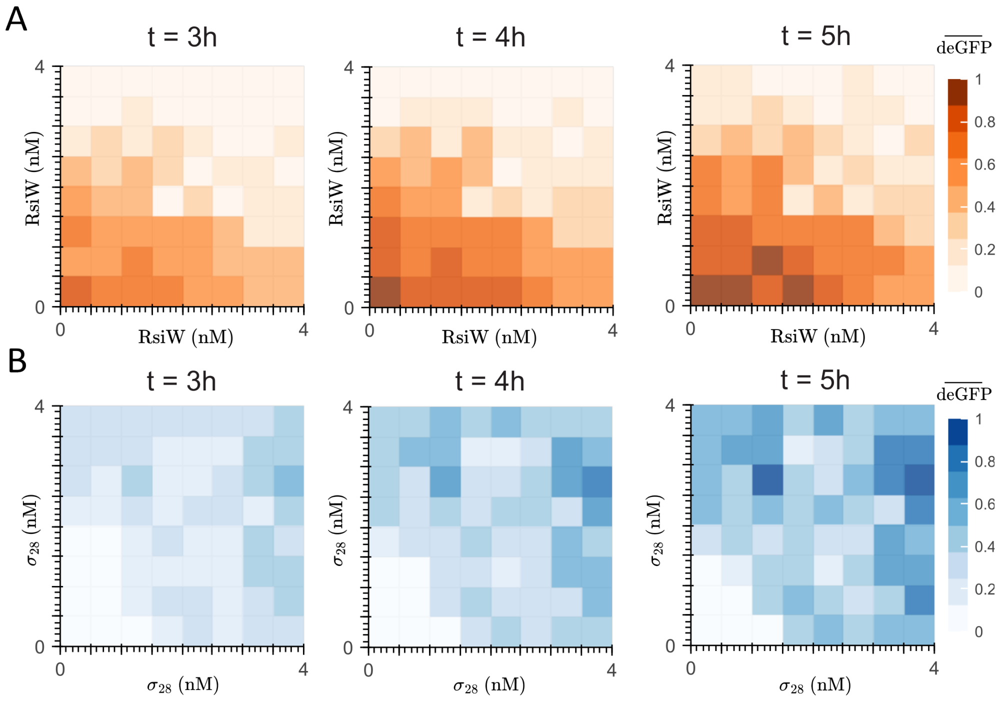

***Figure 13: Flipping the decision boundary and inverting the classification region - Replicates**. Panel A shows the heatmaps for the 3-input implementation based on $\sigma_w$ and RsiW that enables the flipping of the decision boundary. Panel B shows the heatmaps for the 3-input implementation based on $\sigma_{28}$ and FlgM factors that enables the inversion of the classification region. For every implementation, the fluorescence signals in the heatmap were normalized using the maximum fluorescence value recorded at the 5-hour time point.*

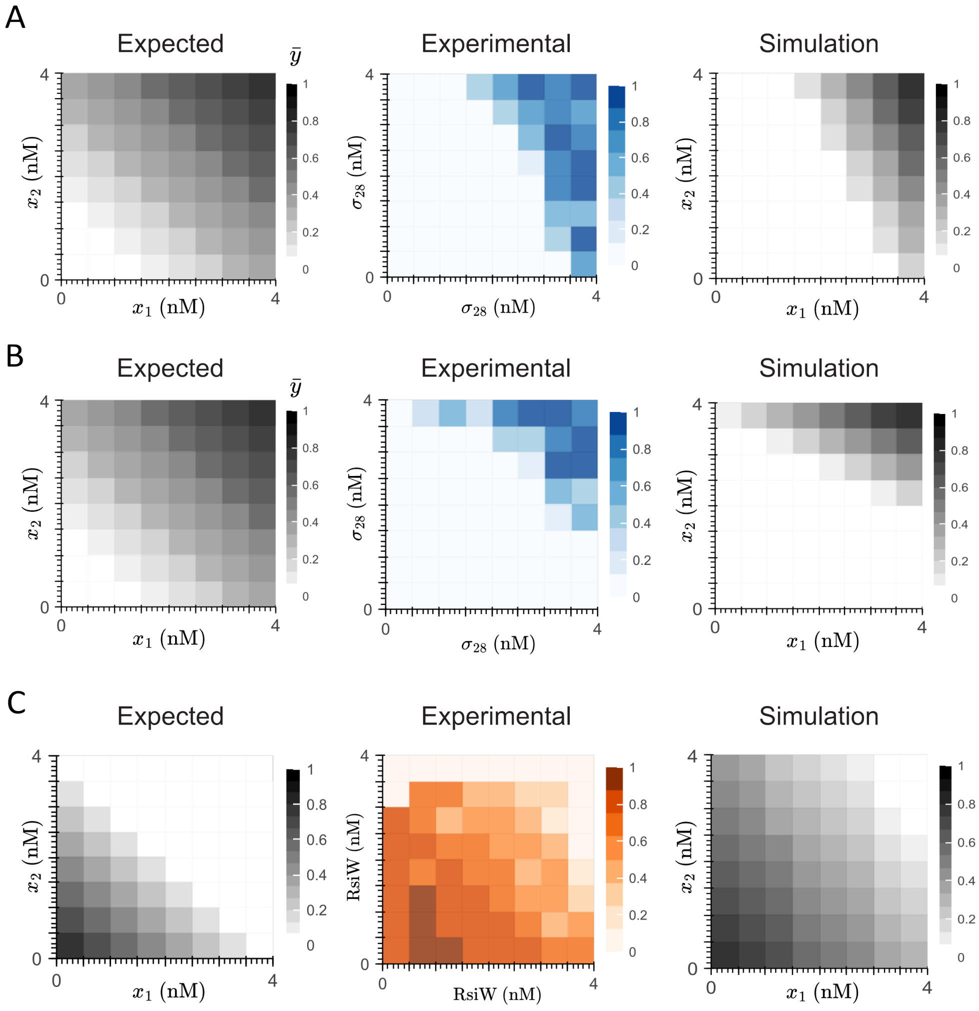

***Figure 14: Model-guided troubleshooting of experiments**. Panels A–C show three representative cases where unexpected patterns deviating from the expected behavior were observed. These deviations were captured by adjusting the weights of the $X_1$ and $X_2$ species and/or the concentration of the bias species $X_0$. Kinetic parameter values for the numerical simulations are as follows: $a = 100$ (/h/μM) and $d = 1$ (/h).*

### 8 Experimental details of in vivo implementation

#### 8.1 Internal ribosome entry site (IRES)-mediated co-expression

As described in the Main Text, we implemented the molecular perceptron using two plasmids, each encoding a sequestering species and co-expressing a fluorescent reporter to estimate its production rate. In summary, one plasmid contains a constitutive promoter (hEF1$\alpha$) driving the expression of mNeonGreen, which includes a cleavage domain at its 3' end, and co-expresses tagBFP as a fluorescent marker. The second plasmid, also under the same constitutive promoter, produces the CRISPR endoRNase Csy4, which binds to the cleavage site on mNeonGreen and mediates its transcript degradation, while co-expressing iRFP720 as its fluorescent marker. To enable independent expression of both genes in each plasmid, an internal ribosome entry site (IRES) separates tagged mNeonGreen from tagBFP in the first plasmid and Csy4 from iRFP720 in the second.

Since the fluorescent genes serve as proxies for production rates, we aimed to achieve similar stoichiometry in co-expression. To this end, we compared the co-expression of tagBFP and mNeonGreen using an IRES against a 2A linker, which enables nearly 1:1 stoichiometry between the upstream and downstream genes [4]. However, we did not use a 2A linker for co-expression because our circuit operates at the post-transcriptional level, whereas a 2A linker requires translational or post-translational regulation. Furthermore, unlike previous implementations (i.e., DNA hybridization and TXTL), where inputs were defined and controlled, here we infer them from fluorescent markers. We generate a wide range of inputs by leveraging the inherent heterogeneity in plasmid copy number from transient lipotransfection and then estimate them based on fluorescence readouts from flow cytometry [5]. Because this approach introduces uncertainty in input estimation, we quantified it by analyzing the "spread" of the tagBFP-to-mNeonGreen expression ratio and further assessed it by comparing the IRES-based co-expression with a co-transfection strategy, where mNeonGreen and tagBFP were expressed from separate plasmids, each under the same constitutive promoter.

For this experiment, 75,000–95,000 cells were collected per sample. Figure 15-A shows the fluorescence thresholds (black dashed lines) captured in the culture without plasmid transfection, which served as a control. After removing saturated events and applying density gating [6], we constructed a normalized histogram (with respect to the maximum count) for each experiment to evaluate uncertainty based on the histogram's spread (Figure 15-B). We visually ranked the experiments by their distribution spread, with the 2A linker exhibiting the least spread (blue), followed by the IRES (orange), and finally the co-transfection, which had the highest spread (purple). To quantify this ranking, we calculated the interquartile range (IR) and median absolute deviation (MAD) for each experiment. The observed differences in spread reflect the variability in co-expression efficiency, with tighter distributions indicating more consistent expression ratios across the population. Figure 15-C shows a strong correlation between tagBFP and mNeonGreen expression when using the 2A linker, with minimal spread. Figure 15-D also shows a high correlation between the two proteins using the IRES, but with greater spread, reflected in higher IR and MAD values. Finally, Figure 15-E displays the co-transfection results, where expression correlation is still present, but with the highest IR and MAD, indicating the most uncertainty.

#### 8.2 Circuit characterization

Once we validated IRES-mediated co-expression, we tested the cleavage of the target sequence upstream of mNeonGreen, which is recognized and cleaved by Csy4 endoRNase, as well as the effect of adding a triple-helix stabilizing sequence from the MALAT1 gene [7]. We compared three constructs: (1) tagBFP and mNeonGreen co-expressed via an IRES (Figure 16-A, Part 1), (2) the same construct with a cleavage

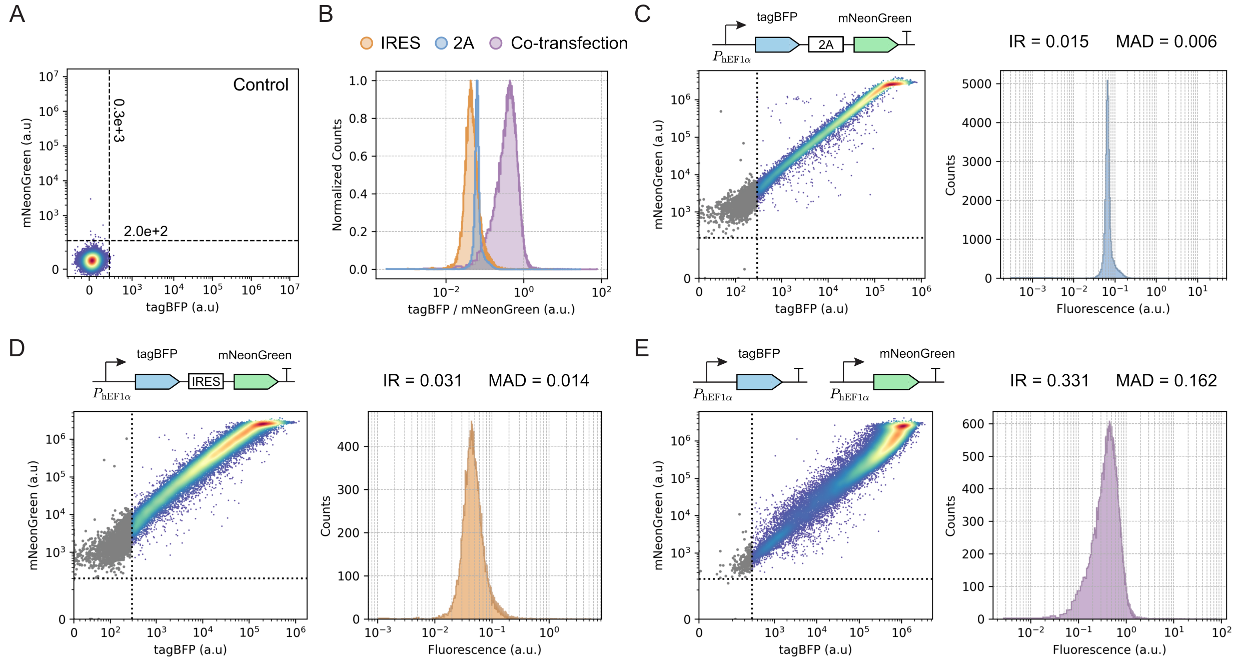

***Figure 15: Characterization of IRES-mediated co-expression in a single construct.*** *Panel A shows the fluorescence thresholds (black dashed lines) for tagBFP* ($0.3 \times 10^3$) *and mNeonGreen* ($2 \times 10^2$)*, estimated from a control culture without plasmid transfection. Panel B presents the normalized histograms of the tagBFP to mNeonGreen fluorescence ratio for the 2A linker (orange), IRES (blue), and co-transfection (purple) experiments, using a bin size of* $10^4$*. Histograms were normalized to their respective maximum event counts. Panels C–E display scatter plots (left) and corresponding histograms (right) for the 2A linker (C), IRES (D), and co-transfection (E) conditions, with interquartile range (IR) and median absolute deviation (MAD) shown. Grey points in the scatter plots correspond to events with fluorescence below the control thresholds, while the colormap represents event density, ranging from low (blue) to high (red).*

domain near the 5' end of the mNeonGreen gene (Figure 16-A, Part 2), and (3) this construct with an additional triple-helix stabilizing sequence (Figure 16-A, Part 3). For the last two constructs, we tested co-expression of tagBFP and mNeonGreen both in the absence and presence of Csy4 by co-transfecting (or not) with a plasmid encoding Csy4 (Figure 16-A, Part 4).

For this experiment, 75,000–110,000 cells were collected per sample. Figure 16-B shows fluorescence thresholds (black dashed lines) in a culture without plasmid transfection, serving as a fluorescence control. Figure 16-C presents the scatter plot for IRES-mediated co-expression of tagBFP and mNeonGreen as a reference. When Csy4 binds and cleaves its target sequence upstream of mNeonGreen, the scatter plot should exhibit a lower slope than this reference, as mNeonGreen levels decrease while tagBFP, positioned upstream of the cleavage site, remains—especially with the triple-helix stabilizing sequence present.

When comparing tagBFP and mNeonGreen co-expression in the absence (top) and presence (bottom) of Csy4 in Figure 16-D, we did not observe the expected decrease in slope relative to Figure 16-C. Instead, a subset of the population showed reduced expression of both fluorescent proteins, suggesting that Csy4 cleavage inhibits mNeonGreen as expected but also affects tagBFP, likely due to transcript destabilization [7]. This effect is further evident in Figure 16-F: on line (2), both fluorescent proteins have similar expression levels and range, while on line (2+4), their distributions shift leftward, indicating a reduction in fluorescence for both. This confirms that Csy4-mediated cleavage effectively reduces mNeonGreen expression while also impacting tagBFP.

On the other hand, when the stabilizing sequence is included in the construct, Figure 16-E shows a subset of the population with a decreased slope relative to Figure 16-C. This effect is further evident in Figure 16-F: on line (3), both fluorescent proteins have similar expression levels and range, while on line (3+4), only the mNeonGreen distribution shifts leftward, indicating a reduction in its fluorescence. Importantly, when comparing Figures 16-C, 16-D (top), and 16-E (top), neither the cleavage sequence nor the stabilizing sequence affect the expression of either protein in the absence of Csy4.

#### 8.3 Linear classification with time-varying inputs

As predicted in equation (110), the reaction does not need to reach steady state to exhibit the decision boundary. We validated this statement by numerically solving equations (79) to (81) and constructing heatmaps in the $x_1$-$x_2$ plane. To match the resolution of the experimental results, we considered a $16\times16$ grid and evaluated the reaction's output at six representative time points (T1 to T6), without loss of generality regarding the reaction timescale, which is expected to be on the order of days for the HEK293 cell line [5]. Figure 17-A shows that the decision boundary is present at every time point, with increasing magnitude over time. To mimic the biological context of our experiments, where the transfection of the sequestering species (i.e., mNeonGreen, modeled by species $X_1$ and produced at rate $w_1$, and the Csy4 endoRNase, modeled by species $X_2$ and produced at rate $w_2$) was transient, we treated the inputs as time-varying. In other words, in equations (79) and (80), we considered $x_1 = x_1(t)$ and $x_2 = x_2(t)$. For comparison, in the TXTL implementation, where no cell division occurs, the inputs were assumed to be constant. However, in vivo, the plasmids encoding the sequestering species undergo dilution due to cell division and are not replicated. To account for this, we repeated the numerical simulations shown in Figure 17-A, but with inputs $x_1$ and $x_2$ modeled as exponentially decaying functions of the form $x(t) = xe^{-\lambda t}$, where $x$ represents the initial plasmid concentration.

Figure 17-B shows that the decision boundary remains detectable as the inputs decay, up to time point T5, as long as plasmid dilution has not yet become significant. While this model does not provide quantitative predictions—unlike models used for DNA hybridization or TXTL—it effectively captures

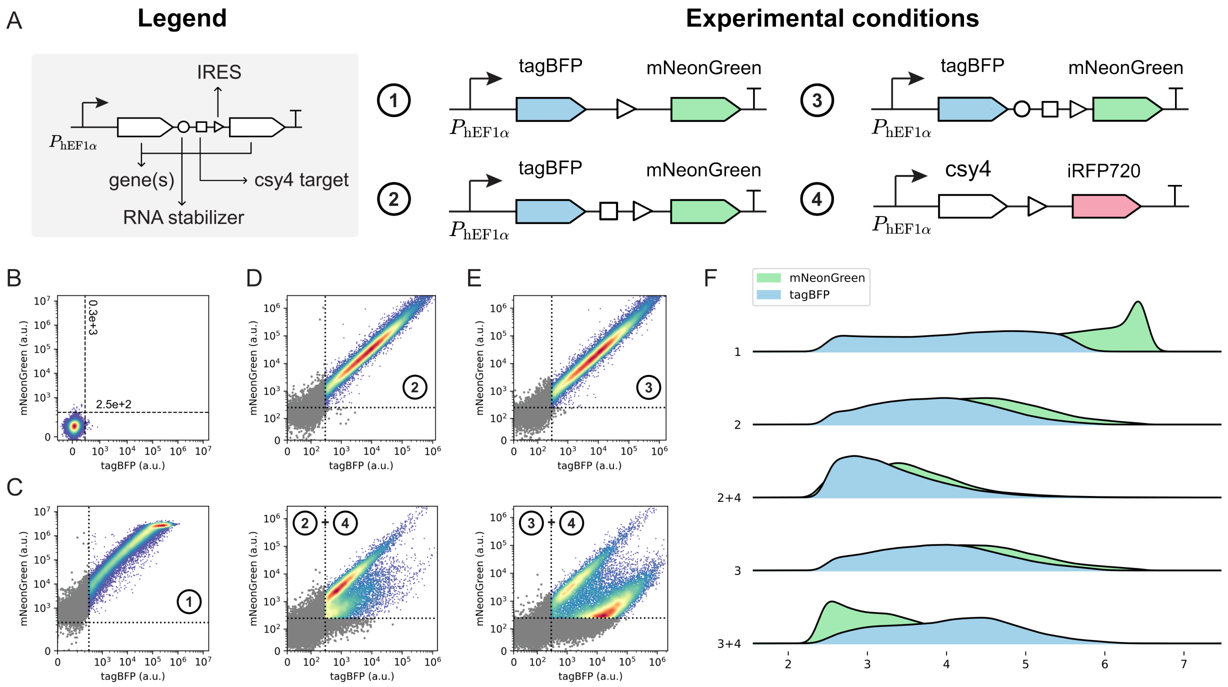

***Figure 16: Effect of Csy4-mediated cleavage and transcript stabilization.*** *Panel A summarizes the four experimental conditions: (1) IRES-mediated co-expression of tagBFP and mNeonGreen, (2) the same construct as (1) with a cleavage domain upstream of the IRES element, (3) the same construct as (2) with a triple-helix stabilizing sequence [7] upstream of the cleavage domain, and (4) IRES-mediated co-expression of Csy4 endoRNase and the iRFP720 fluorescent marker. Panel B shows fluorescence thresholds (dashed lines) for tagBFP* $(0.3 \times 10^3)$ *and mNeonGreen* $(2.5 \times 10^2)$*, estimated from a control culture without plasmid transfection. Panel C presents the scatter plot for the reference IRES-mediated construct (1). Panel D shows scatter plots for the construct with the cleavage domain (2), without (top) and with (bottom) co-transfection of Csy4 (4). Panel E displays scatter plots for the construct with both the cleavage domain and the stabilizing sequence (3), without (top) and with (bottom) co-transfection of Csy4 (4). Panel F shows the fluorescence distribution of tagBFP (blue) and mNeonGreen (green) on a logarithmic scale for each condition. In scatter plots, grey points represent events below control fluorescence thresholds, while the colormap indicates event density from low (blue) to high (red).*

the temporal pattern recognition feature, even under time-varying inputs with exponentially decaying dynamics.

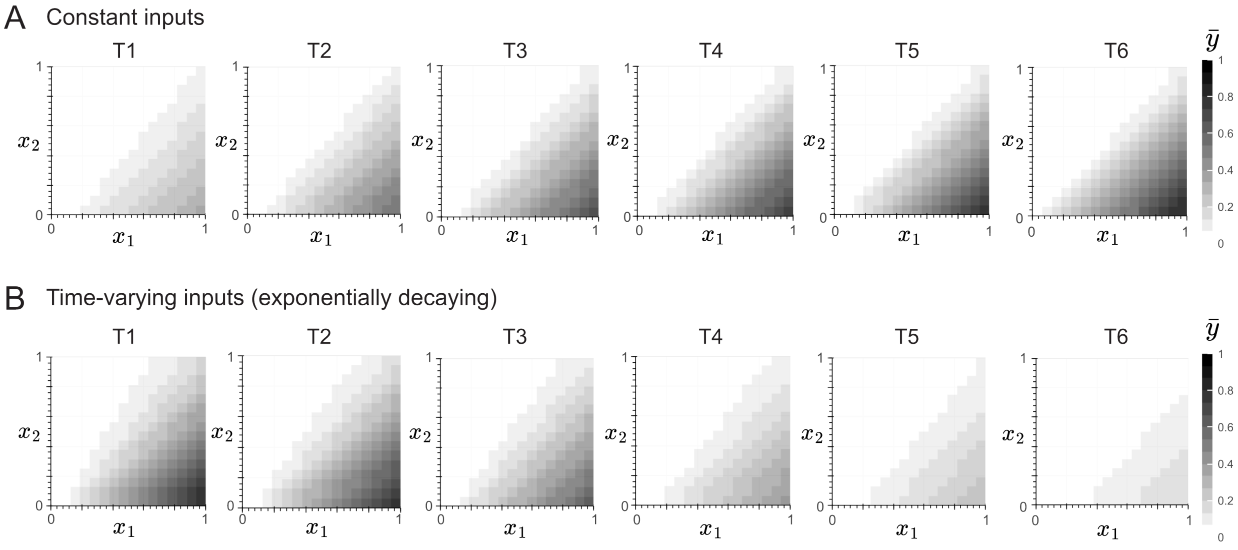

***Figure 17: Effect of exponentially decaying inputs on the decision boundary****. Panels A and B show heatmaps at 6 time points, obtained by solving equations* (79) *to* (81) *with a 16x16 resolution and the following production rates: $w_1 = w_2 = 1$ (/h). Panel A uses constant inputs $x_1$ and $x_2$ (0–1 $\mu$M), while Panel B uses exponentially decaying inputs $x(t) = xe^{-\lambda t}$ with $\lambda = 1$ (/h) and $x = 1$ $\mu$M. Kinetic parameter values for the numerical simulations are as follows: $a = 100$ (/h/$\mu$M), $d = 1$ (/h), $\delta = 0.5$ (/h), and $\phi = 1$ (/h)*

#### 8.4 Activation function

Equation (110) predicts a time-dependent Rectified Linear Unit (ReLU)-like activation function, assuming a sufficiently small dissociation constant, similar to the TXTL system. In fact, with minimal degradation ($\delta \to 0$), we recover the activation function expression derived for the TXTL system (compare equations (75) and (109)). Following the TXTL system analysis (Section 7.1, Activation Function), we aimed to reconstruct the activation function by plotting the fluorescence response (mNeonGreen) against the normalized x-axis ($x_1 w_1$ relative to $x_2 w_2$). Since we used tagBFP as a proxy for $x_1 w_1$ and iRFP720 for $x_2 w_2$, binning was applied before constructing the function, as illustrated in Figure 18. This produced a 16$\times$16 grid, where we computed the median tagBFP, iRFP720, and mNeonGreen fluorescence for each bin. We then plotted the difference between tagBFP and iRFP720 against mNeonGreen, as shown in the Main Text.

Figure 18-A shows fluorescence thresholds (black dashed lines) from a culture without plasmid transfection, serving as a control. No threshold was imposed on mNeonGreen fluorescence to increase the sample size near the tagBFP $\sim$ iRFP720 threshold. The flow cytometer scatterplot is used to estimate the $x_1$-$x_2$ heatmap presented for both the DNA hybridization and TXTL implementations. We focused on the upper-right quadrant (Figure 18-B), which contains cells with fluorescence above both thresholds. Data was filtered accordingly and further binned: first into 16 bins by iRFP720 fluorescence (horizontal, Figure 18-C), then into 16 bins by tagBFP fluorescence (vertical, Figure 18-D), forming a 16$\times$16 grid. We also tested 8$\times$8 and 32$\times$32 grids, but 16$\times$16 provided a balance between sample size and resolution.

The 8×8 grid lacked sufficient sample size to properly observe the activation function, while the 32×32 grid introduced sparsity, with bins containing nearly zero events (data not shown).

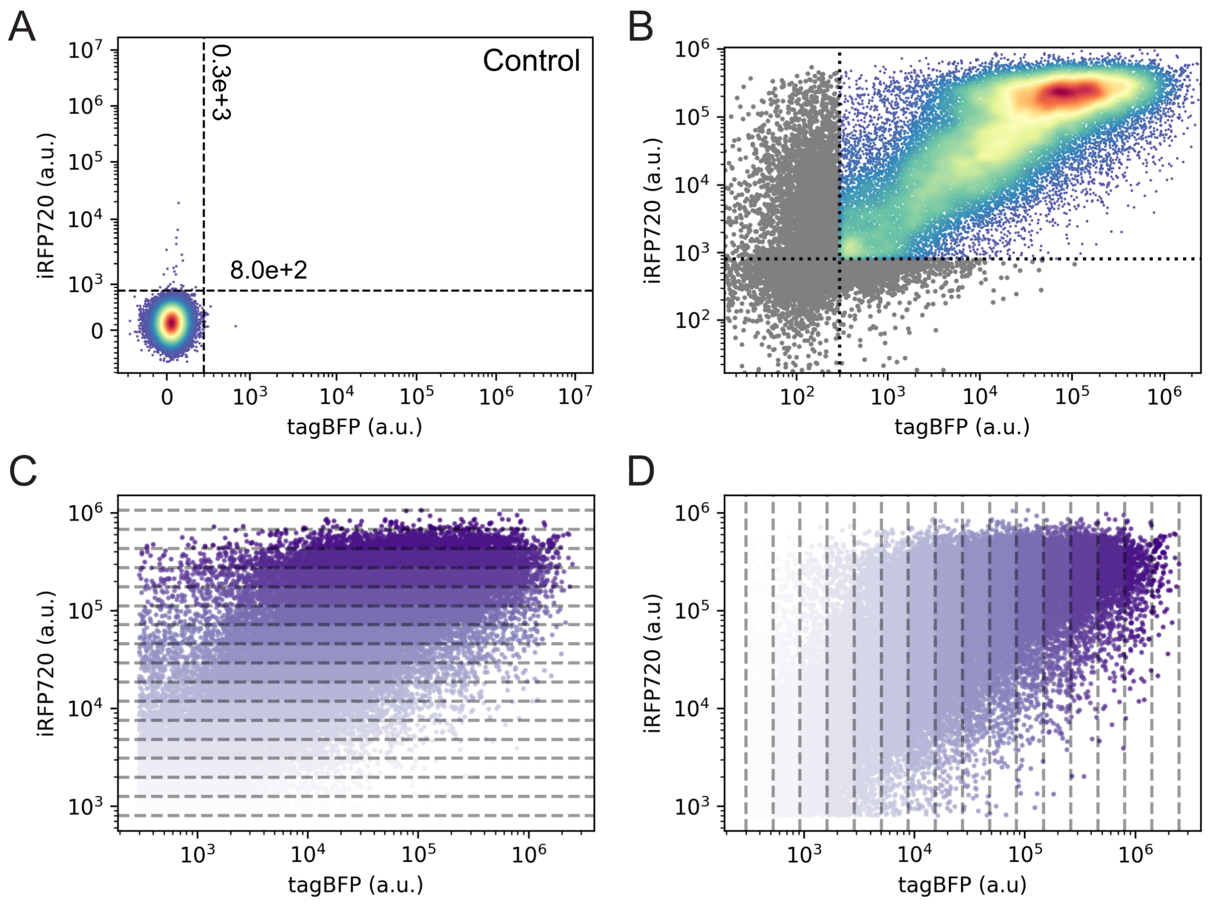

***Figure 18: Binning**. Panel A shows the fluorescence thresholds (black dashed lines) for tagBFP* $(0.3 \times 10^3)$ *and iRFP720* $(8 \times 10^2)$*, estimated from a control without plasmid transfection. Panel B presents the scatter plot of the experiment used to estimate the activation function (see Section 8.1 Internal ribosome entry site (IRES)-medianted co-expression). Panels C and D show the data from the upper-right quadrant of Panel B, binned by iRFP720 fluorescence (horizontal, C) and by tagBFP fluorescence (vertical, D). Bins are numbered from bottom to top and left to right, from 1 to 16, and color-coded from light purple (bin = 1) to dark purple (bin = 16).*

#### 8.5 Deconvoluting fluorescence measurements for input estimation

Since our poly-transfection protocol did not achieve 100% efficiency in covering the input space, we selected a subset of data that maximized the dynamic range of tagBFP and iRFP720 fluorescence. The dynamic range is defined as the spread between the lowest and highest fluorescence values after filtering, which was based on the control's thresholds. We focused on maximizing this range while ensuring enough events per bin to avoid sparsity, so we can confidently reconstruct the heatmaps in the $x_1$-$x_2$ plane. To do this, we developed a method detailed below:

1. Data is filtered (Figure 20-A) using the thresholds defined with the control experiment (e.g., Figures 18-A). In this example, the upper-right quadrant of the scatterplot shows a tagBFP range

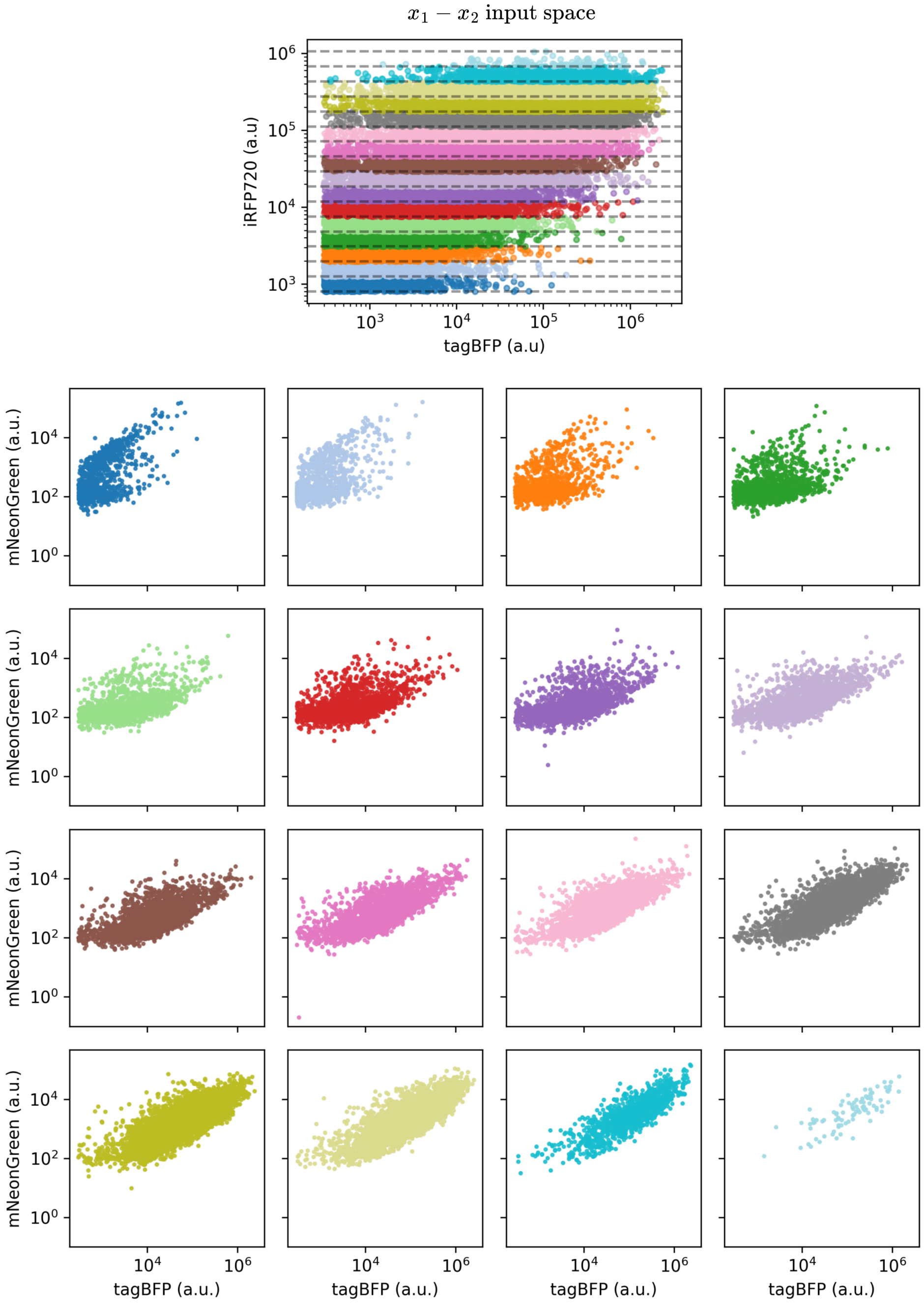

***Figure 19: Input/output map***. *According to Section 8.4, after filtering the data and divide it in 16 bins based on the iRFP720 fluorescence, the input (tagBFP)-output(mNeonGreen) can be reconstructed.*

of $0.3 \times 10^3$ to $2.5 \times 10^6$ and an iRFP720 range of $8 \times 10^2$ to $1.1 \times 10^6$

2. We divided the upper-right quadrant of the scatterplot into rectangular regions, each labeled by the percentage of the dynamic range covered (Figure 20-B). For example, the '25% rectangle' is defined by selecting 25% of the maximum tagBFP and iRFP720 values, respectively, and adding them to the minimum values after filtering (e.g., leftmost region in Figure 20-B).

3. Since poly-transfection—specifically, the number of plasmid copies a cell receives—is a random process, we approximate it using a Poisson model, as supported by prior studies [5, 8]. Given that we lack direct measurements of plasmid counts per cell but assume that, after filtering, each cell exhibits at least one copy of any plasmid, we instead model the number of cells per bin as a Poisson-distributed variable.

   When dividing the data into a 16x16 grid, we computed the expected number of cells per bin as:

$$\lambda = \frac{\text{Total number of cells}}{16^2}$$

   To quantify how many bins contain a statistically significant number of cells, we define a minimum count threshold using a one-sided 95% confidence bound under a normal approximation to the Poisson distribution, given by

$$\text{min count} = \lceil \lambda - k \cdot \sqrt{\lambda} \rceil$$

   Here, $\lceil \ \rceil$ corresponds to the "ceil" operation to ensure an integer number of cells per bin, while $k = 1.645$ corresponds to the 95th percentile of the standard normal distribution.

4. We began with the 2% rectangle and counted how many bins contained more events than the previously estimated min count. This process was repeated in 2% increments until the entire dynamic range was covered, as shown in Figure 20-C.

5. We computed the cumulative sum (CUSUM) of the counts across the dynamic range and determined the percentile at which the maximum CUSUM occurs. In this example, the maximum is observed at 44%, as shown in Figure 20-D.

From the percentile calculated in Step 5 (e.g., Figure 20-E), the area is binned as described in Figure 18, and heatmaps are constructed by taking the median mNeonGreen fluorescence per bin. Additionally, Figure 20-F shows the input coverage calculated in Step 5, color-coded by the number of events in each bin, while Figure 20-G compares the mNeonGreen fluorescence distribution across the entire dynamic range (black) with the calculated percentile (green).

#### 8.6 Reconstructed heatmaps

Following the method outlined in Section 8.5 Deconvoluting fluorescence measurements for input estimation, we constructed heatmaps in the $x_1$-$x_2$ plane for the endoRNase-based in vivo implementation of the molecular perceptron, as shown in Figure 21 for three different plasmid stoichiometries. Specifically, Figure 21-A corresponds to a dynamic range of 44%, Figure 21-B to 40%, and Figure 21-C to 53%.

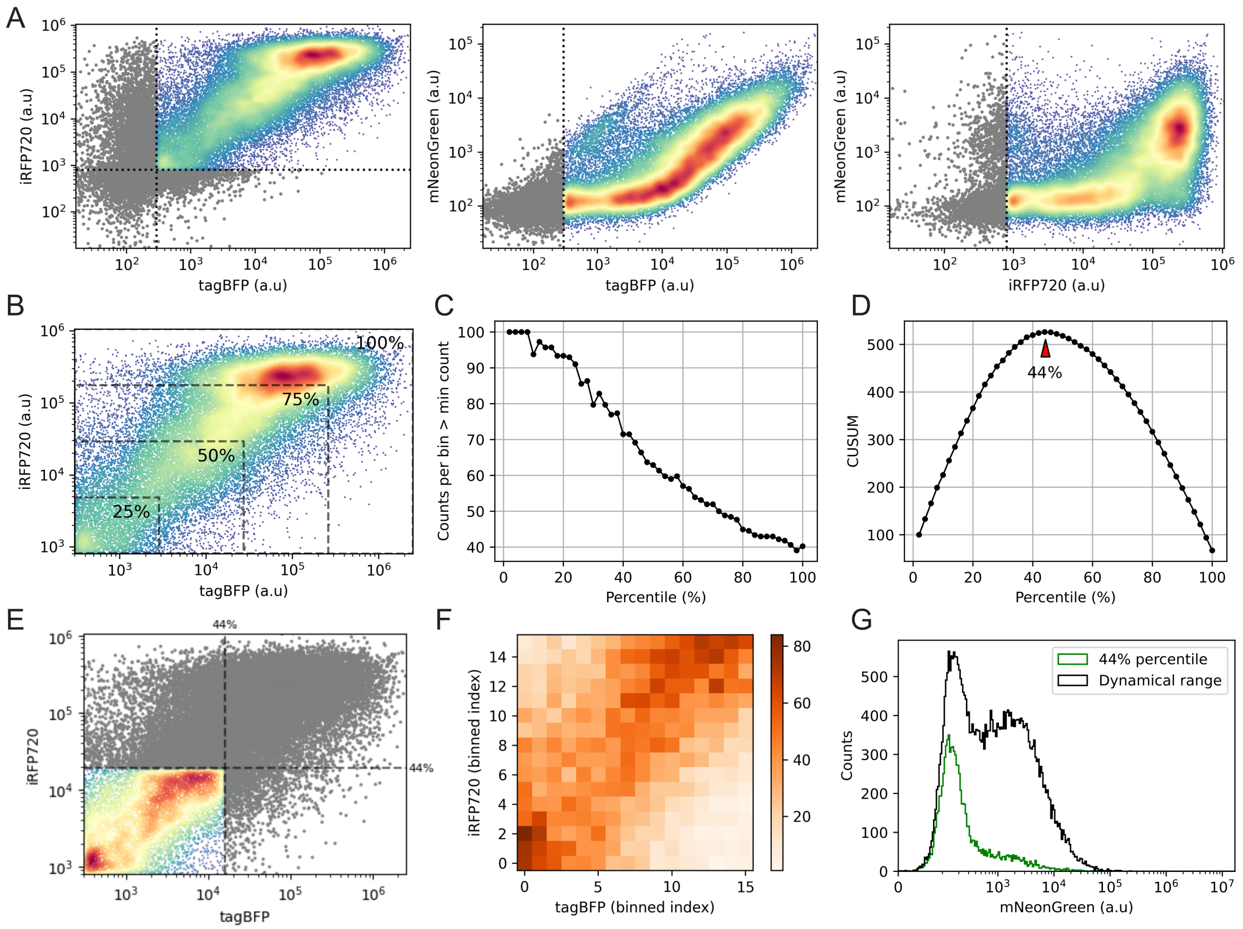

***Figure 20: Method to maximize the dynamical range for constructing the heatmaps in the $x_1$-$x_2$ plane.*** *Panel A displays scatter plots of the data filtered by fluorescence thresholds, showing the following combinations (from left to right): tagBFP versus iRFP720 (input space), tagBFP versus mNeonGreen (input-output map with respect to $x_1$), and iRFP720 versus mNeonGreen (input-output map with respect to $x_2$). Panel B illustrates the 'percentile rectangles.' Panel C quantifies the number of bins containing a statistically significant number of cells based on a minimum count threshold. Panel D presents the cumulative sum from Panel C, highlighting the percentile that maximizes the dynamic range, which is further illustrated in Panel E. Panel F shows a heatmap of cell counts per bin, with color coding from low (light orange) to high (dark orange). Panel G compares the mNeonGreen fluorescence distribution for the entire dynamic range (upper-right quadrant from Panel B) with the 44% percentile calculated in Panel D.*

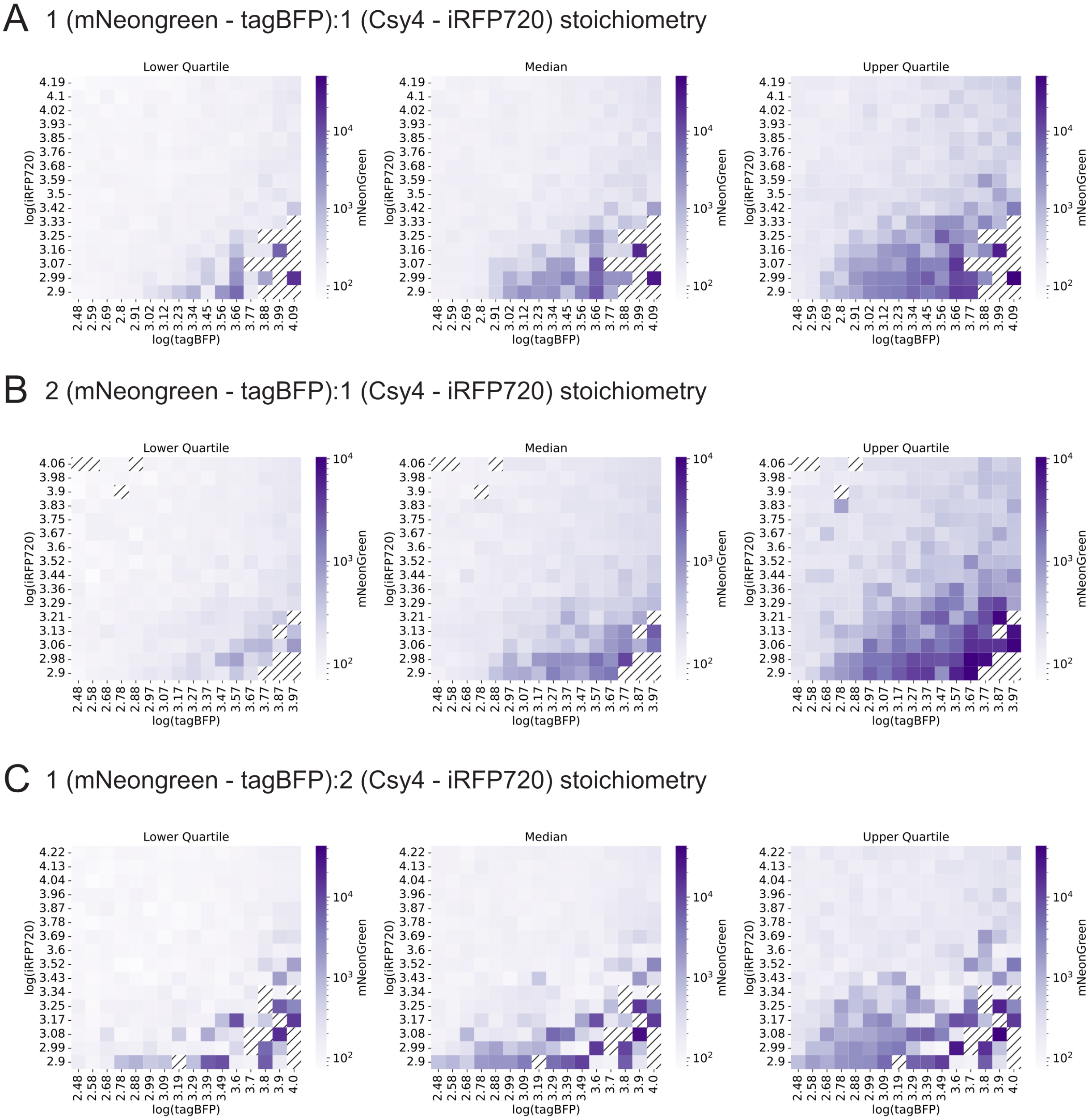

***Figure 21: The endoRNAse-based implementation of the molecular perceptron exhibits a decision boundary**. From top to bottom, each panel represents different plasmid stoichiometries: (A) 1:1, (B) 1:2, and (C) 2:1, where '1' corresponds to a plasmid concentration of 400 ng/$\mu$L and '2' corresponds to 800 ng/$\mu$L. From left to right, the mNeonGreen fluorescence per bin is estimated using the lower quartile (25th percentile, left column), the median (50th percentile, middle column), and the upper quartile (75th percentile, right column). tagBFP, iRFP720, and mNeonGreen fluorescence are shown on a logarithmic scale, with color coding from light purple (low) to dark purple (high). Black stripes indicate bins without a statistically significant number of cells.*

#### 8.7 Weight tuning

##### 8.7.1 Production rate and degradation-based tuning

From equation (110), we observe a similar mechanism for adjusting the slope of the decision boundary to the TXTL system (Section 7.3 Weight tuning): adjusting the production rates associated with the $X_1$ and $X_2$ species ($w_1$ and $w_2$, respectively). In Figure 22-A, we increase the slope of the decision boundary by decreasing the production rate associated with the $X_2$ species ($w_2$, corresponding to the production of the endoRNAse), and Figure 22-B shows that this property also holds for exponentially decaying inputs. Furthermore, equation (110) shows that the magnitude of $y(t)$ can be adjusted through the degradation rate, resulting in a rescaled version of the decision boundary. Figures 22-C and 22-D illustrate how increasing the degradation rate of the $Z$ species (i.e., the endoRNAse) affects the decision boundary for constant and exponentially decaying inputs, respectively.

##### 8.7.2 Characterization of uORFs

Following the results from the numerical simulations in Figure 22-B, we reduce the production rate of endoRNase (e.g., $w_2$) by using upstream open reading frames (uORFs), which suppress protein production by reducing the translation rate [9]. The more uORFs are repeated, the stronger the inhibition, leading to a greater increase in the slope of the decision boundary. We tested 1, 3, and 5 repetitions of uORFs upstream of a mNeonGreen gene, followed by an IRES sequence that enables co-expression of tagBFP. With this setup, we expect to see a decrease in mNeonGreen expression as more uORFs are added.

For this experiment, 77,000 – 110,000 cells were collected per sample. Figure 23-A shows fluorescence thresholds (black dashed lines) from a control culture with no plasmid transfection. For each circuit (no uORFs, 1, 3, and 5 repetitions), we calculated the slope of the filtered data in scatterplots of tagBFP versus mNeonGreen, along with the $R^2$ value to assess linearity. Figure 23-B shows the scatter plot for IRES-mediated co-expression of tagBFP and mNeonGreen as a reference, since it doesn't include uORFs, along with the corresponding slope and $R^2$. Figures 23-C, 23-D, and 23-E show the scatter plots for 1, 3, and 5 uORF repetitions upstream of mNeonGreen, respectively. Upon comparison, we observe how the addition of uORFs reduces the slope (calculated as tagBFP over mNeonGreen), where the highest slope corresponds to no uORFs and the lowest to 5 repetitions. Finally, in Figure 24, we compute the distribution of the tagBFP/mNeonGreen ratio for each condition. Adding uORFs shifts the distribution to the right, and combined with the scatter plots in Figure 23, this suggests a reduction in mNeonGreen expression.

##### 8.7.3 Reconstructed heatmaps: 5xuORFs

Following the method described in Section 8.5 Deconvoluting fluorescence measurements for input estimation, we constructed heatmaps in the $x_1$-$x_2$ plane for a construct based on the one described in Section 8.7.3, but including a PEST degron downstream of the Csy4 endoRNase gene. The effect of this degron on the half-life of the upstream protein has been previously characterized [10]. We tested three different plasmid stoichiometries, as shown in Figure 26. Specifically, Figure 26-A corresponds to a dynamic range of 38%, Figure 26-B to 40%, and Figure 26-C to 41%.

##### 8.7.4 Reconstructed heatmaps: PEST degron

Following the method described in Section 8.5 Deconvoluting fluorescence measurements for input estimation and building on the circuit proposed in Section 8.7.3 Reconstructed heatmaps: 5xuORFs, we constructed heatmaps in the $x_1$-$x_2$ plane by adding a PEST degron downstream of the Csy4 endoRNase

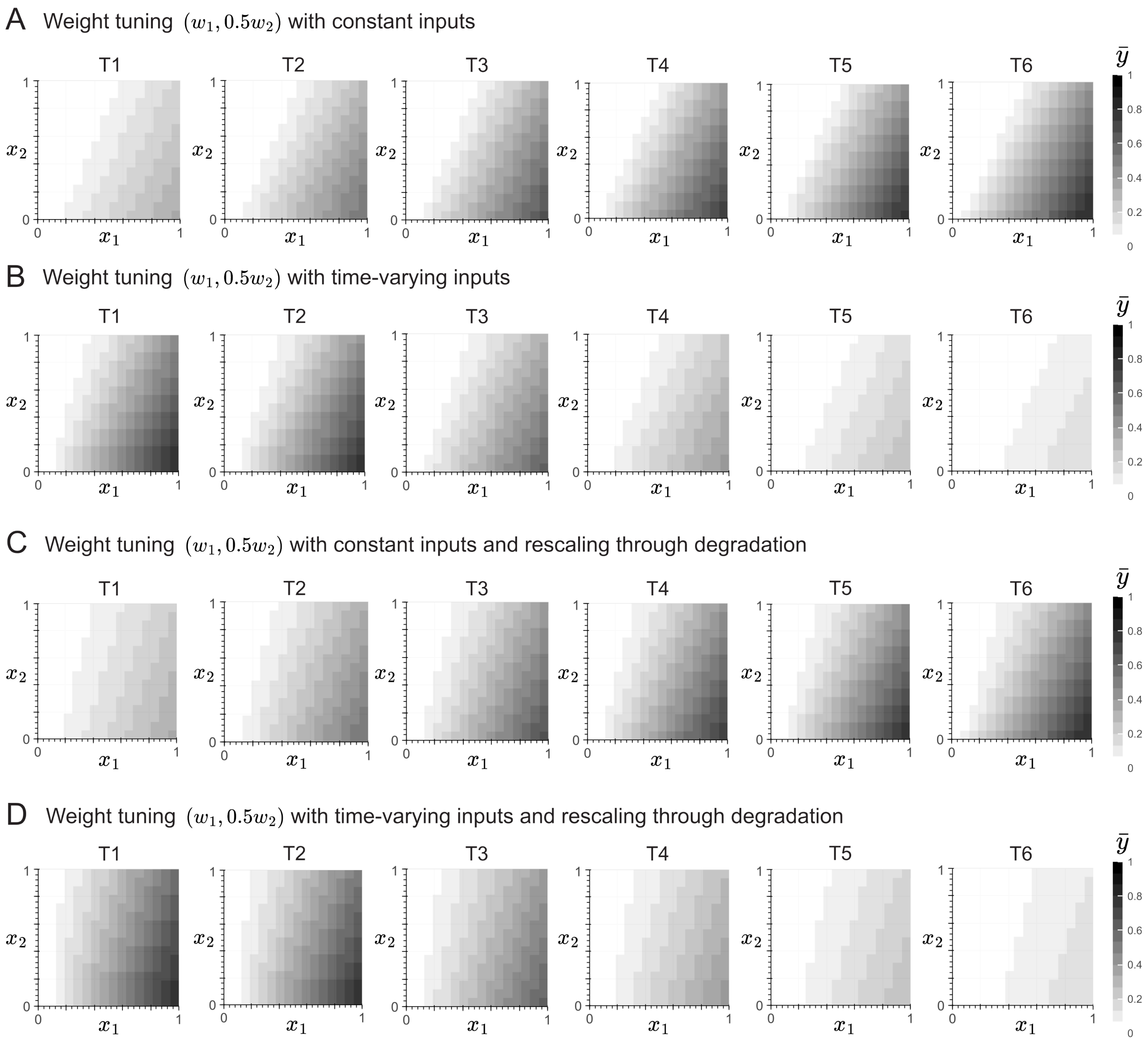

***Figure 22: Weight tuning via production rate adjustment and degradation rate for constant and exponentially decaying inputs.***. *Panels A to D show heatmaps at 6 time points, obtained by solving equations* (79) *to* (81) *with a 16x16 resolution. Panels A and B use the following nominal parameters to simulate an increase in the decision boundary:* $w_1 = 1$ *(/h)* $w_2 = 0.5$, $a = 100$ *(/h/*$\mu$*M),* $d = 1$ *(/h),* $\delta = 0.5$ *(/h), and* $\phi = 1$ *(/h). Panel C and D use the same nominal parameters, except for* $\phi = 10$ *(/h) to simulate the re-scaling of the decision boundary. Both Panel A and C use constant inputs* $x_1$ *and* $x_2$ *(0-1* $\mu$*M), while Panel B and D use exponentiall decaying inputs* $x(t) = xe^{-\lambda t}$ *with* $\lambda = 1$ *(/h) and* $x = 1$ $\mu$*M.*

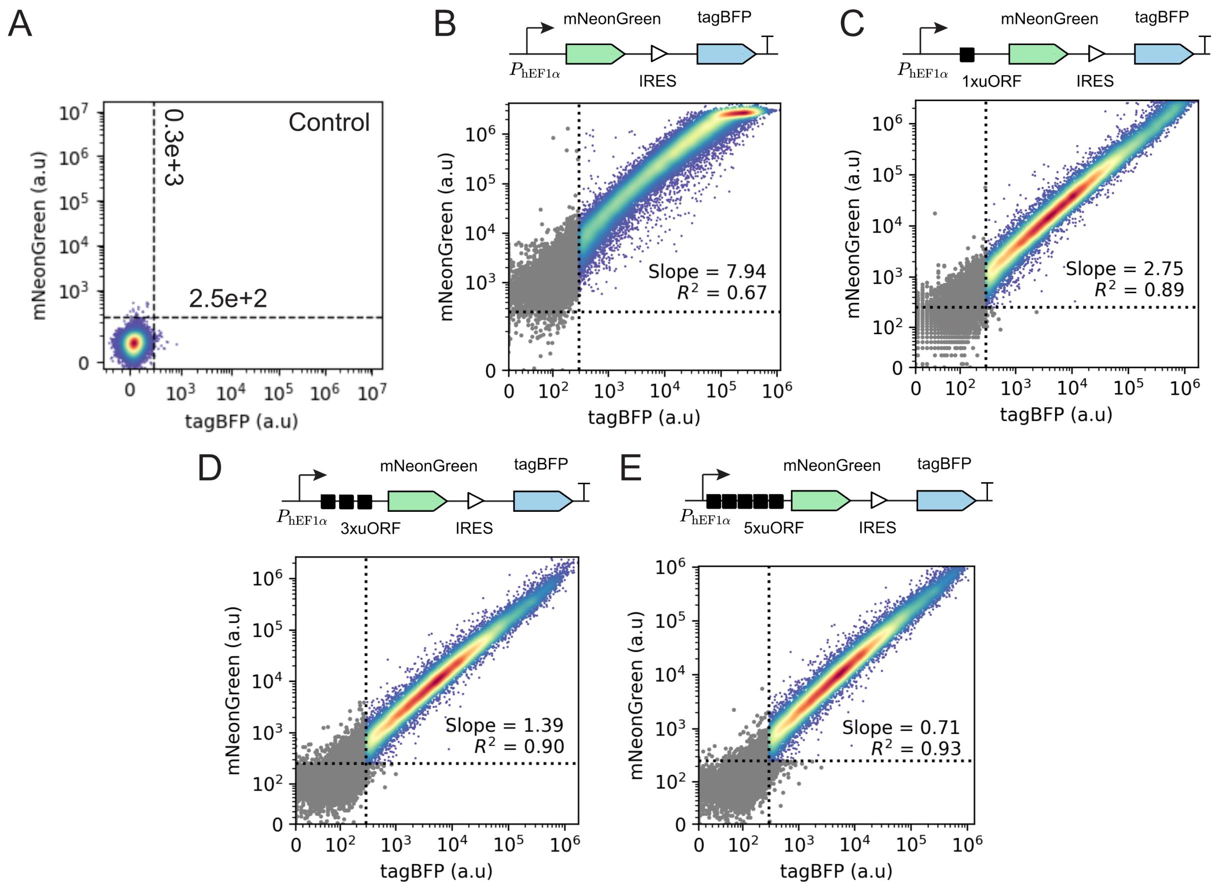

***Figure 23: Characterization of uORFs for weight tuning - Scatter plots..*** *Panel A shows the fluorescence thresholds (black dashed lines) for tagBFP* $(0.3 \times 10^3)$ *and mNeonGreen* $(2.5 \times 10^2)$*, estimated from a control culture without plasmid transfection. Panel B presents the scatter plot for the reference IRES-mediated construct. Panels C to E show the scatter plots for the construct with 1 (C), 3 (D) and 5 (E) repetitions of the open reading frame (uORF). The slope and associated* $R^2$ *value are calculated per scatter plot. In all scatter plots, grey points represent events below control fluorescence thresholds, while the colormap indicates event density from low (blue) to high (red)*

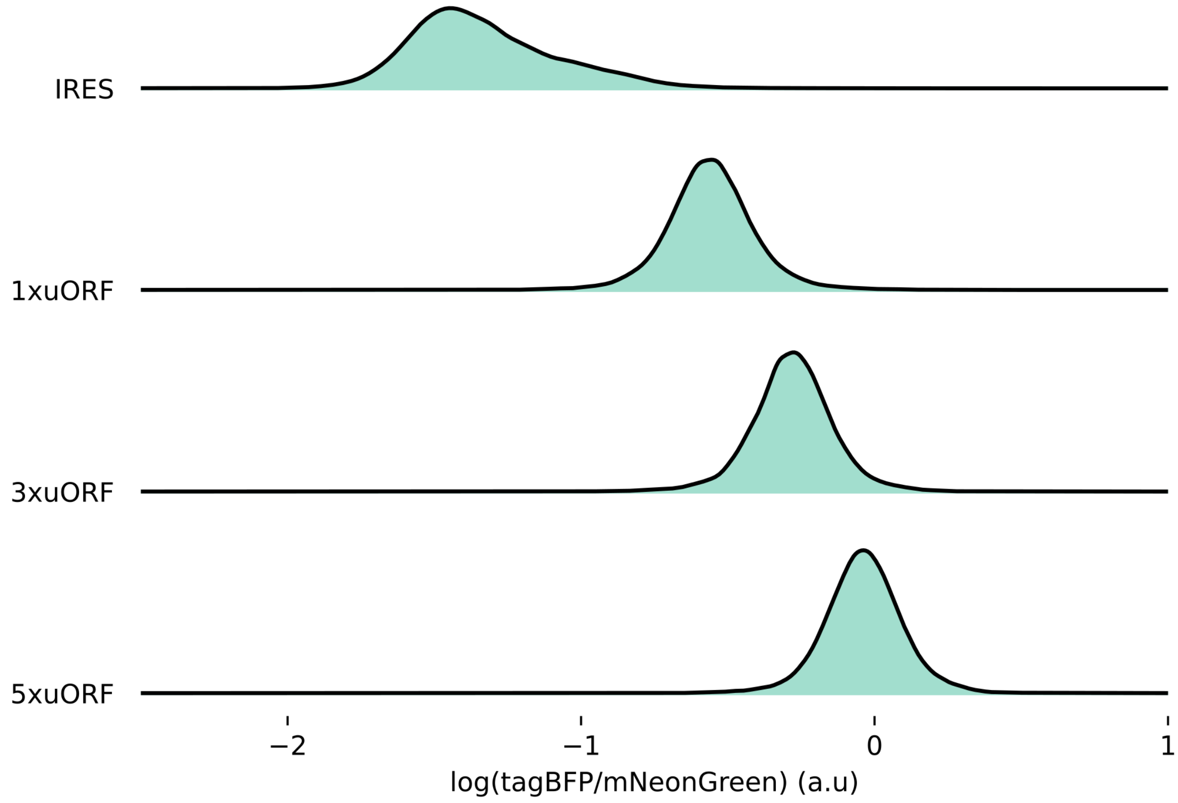

***Figure 24: Characterization of uORFs for weight tuning - Distributions.***. *The fluorescence distribution of the ratio between tagBFP and mNeonGreen on a logarithmic scale for each condition. From top to bottom, the (reference) IRES-mediated co-expression of tagBFP and iRFP720, 1, 3, and 5 repetitions of the uORFs*

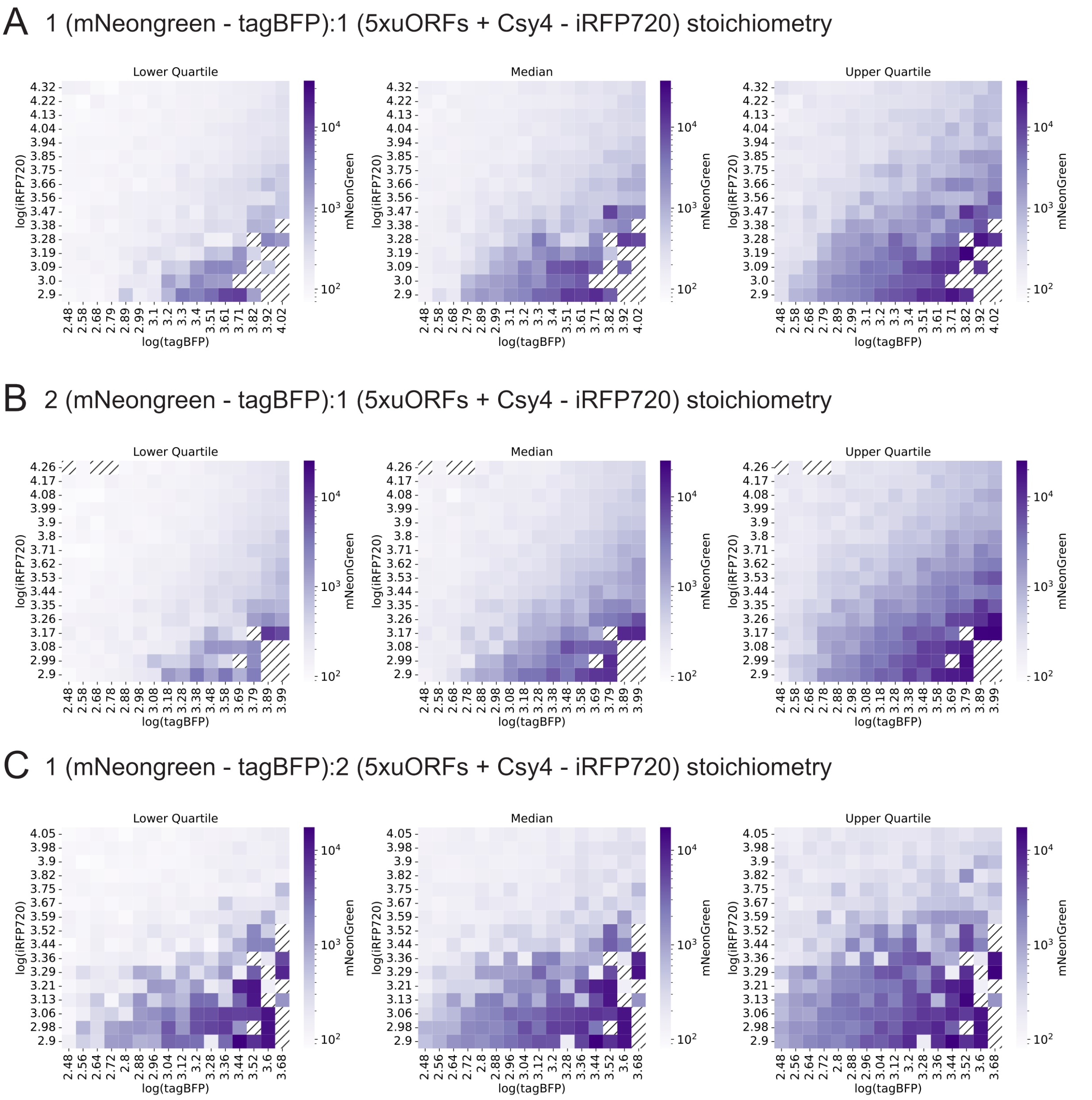

***Figure 25: Weight tuning using 5xuORFs***. *From top to bottom, each panel represents different plasmid stoichiometries: (A) 1:1, (B) 1:2, and (C) 2:1, where '1' corresponds to a plasmid concentration of 400 ng/$\mu$L and '2' corresponds to 800 ng/$\mu$L. From left to right, the mNeonGreen fluorescence per bin is estimated using the lower quartile (25th percentile, left column), the median (50th percentile, middle column), and the upper quartile (75th percentile, right column). tagBFP, iRFP720, and mNeonGreen fluorescence are shown on a logarithmic scale, with color coding from light purple (low) to dark purple (high). Black stripes indicate bins without a statistically significant number of cells.*

gene. The effect of the degron on the half live of the protein that is encoded upstream to it has been previosly characterized [10]. We tested three different plasmid stoichiometries, as shown in Figure 26. Speficially, Figure 26-A corresponds to a dynamic range of 38%, Figure 26-B to 40%, and Figure 26-C to 41%.

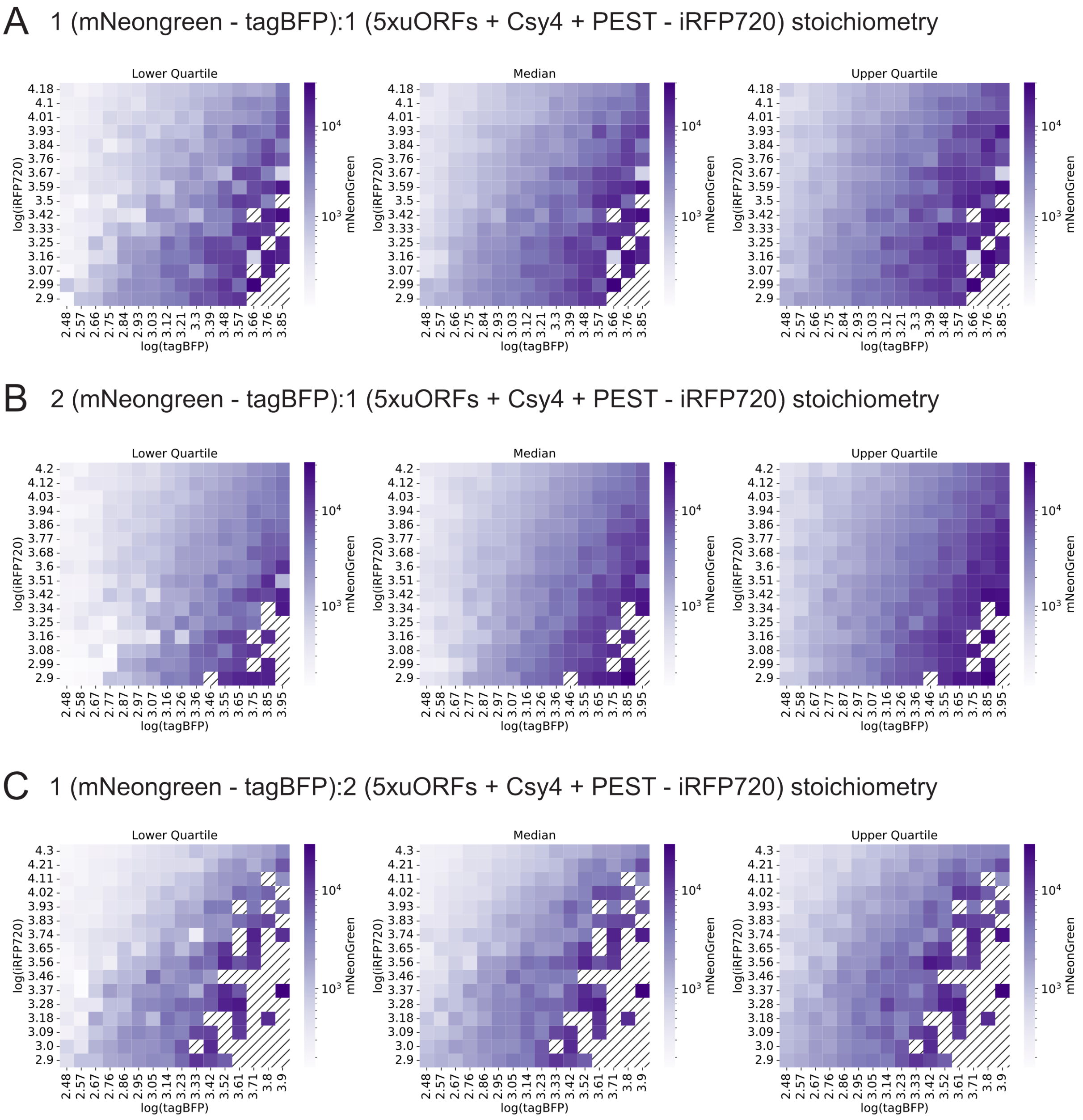

***Figure 26: Re-scaling the decision boundary by changing the degradation rate through a PEST degron****.* *From top to bottom, each panel represents different plasmid stoichiometries: (A) 1:1, (B) 1:2, and (C) 2:1, where '1' corresponds to a plasmid concentration of 400 ng/$\mu$L and '2' corresponds to 800 ng/$\mu$L. From left to right, the mNeonGreen fluorescence per bin is estimated using the lower quartile (25th percentile, left column), the median (50th percentile, middle column), and the upper quartile (75th percentile, right column). tagBFP, iRFP720, and mNeonGreen fluorescence are shown on a logarithmic scale, with color coding from light purple (low) to dark purple (high). Black stripes indicate bins without a statistically significant number of cells.*
